# Transcriptome and hormone regulations shape drought stress-dependent Fusarium Head Blight susceptibility in different barley genotypes

**DOI:** 10.1101/2025.11.23.689882

**Authors:** Felix Hoheneder, Christina Elisabeth Steidele, Michael Gigl, Corinna Dawid, Ralph Hückelhoven

**Affiliations:** Chair of Phytopathology, TUM School of Life Sciences, HEF World Agricultural Systems Center, Technical University of Munich, Emil-Ramann Str. 2, 85354 Freising-Weihenstephan, Germany; Junior Research Group Food Processing and Health, ZIEL Institute for Food and Health, Technical University of Munich, Lise-Meitner-Straße 34, 85354 Freising-Weihenstephan, Germany; Professorship for Chemosensory Food Systems, TUM School of Life Sciences, Technical University of Munich, Lise-Meitner-Straße 34, 85354 Freising-Weihenstephan, Germany; Leibniz Institute for Food Systems Biology at the Technical University of Munich, Lise-Meitner-Straße 34, 85354 Freising-Weihenstephan, Germany

**Keywords:** Fusarium Head Blight, Barley, Drought, Transcriptomics, Abscisic acid, Stress combination

## Abstract

Little is known about regulatory mechanisms that crop plants use to respond to combinations of abiotic and biotic stress. We analysed four barley genotypes under simultaneous *Fusarium culmorum* infection and drought stress by phenotyping of Fusarium Head Blight (FHB) disease, hormone profiling, and transcriptome analysis. FHB severity was host genotype-dependent, with moderately resistant cultivars Avalon and Barke showing increased FHB under drought, while drought did not further increase FHB severity in highly susceptible Morex and Palmella Blue. Transcriptome analysis revealed largely additive effects of single stresses, with drought-dominated regulation with increasing drought severity at later stages. Co-expression analysis connected abscisic acid and auxin to gene expression modules functionally enriched with stress-specific physiological responses. Stress-response genes, uniformly expressed across genotypes, were linked to pathogen defence, detoxification, and drought adaptation, whereas a cluster of hundreds of moderately *Fusarium*-responsive genes was limited in up-regulation under combined stress, possibly explaining enhanced FHB severity under drought. A multiple linear regression model accurately predicted combined stress expression from single stress responses, demonstrating modularity of barley’s stress regulation under combined drought and *Fusarium* stress.

**Graphical Abstract:** Simplified working model illustrating drought effects on abscisic acid (ABA) levels and susceptibility to Fusarium Head Blight (FHB) of barley, and key processes such as storage and desiccation responses, photosynthesis, cell wall formation, metabolism, and transferase activity.

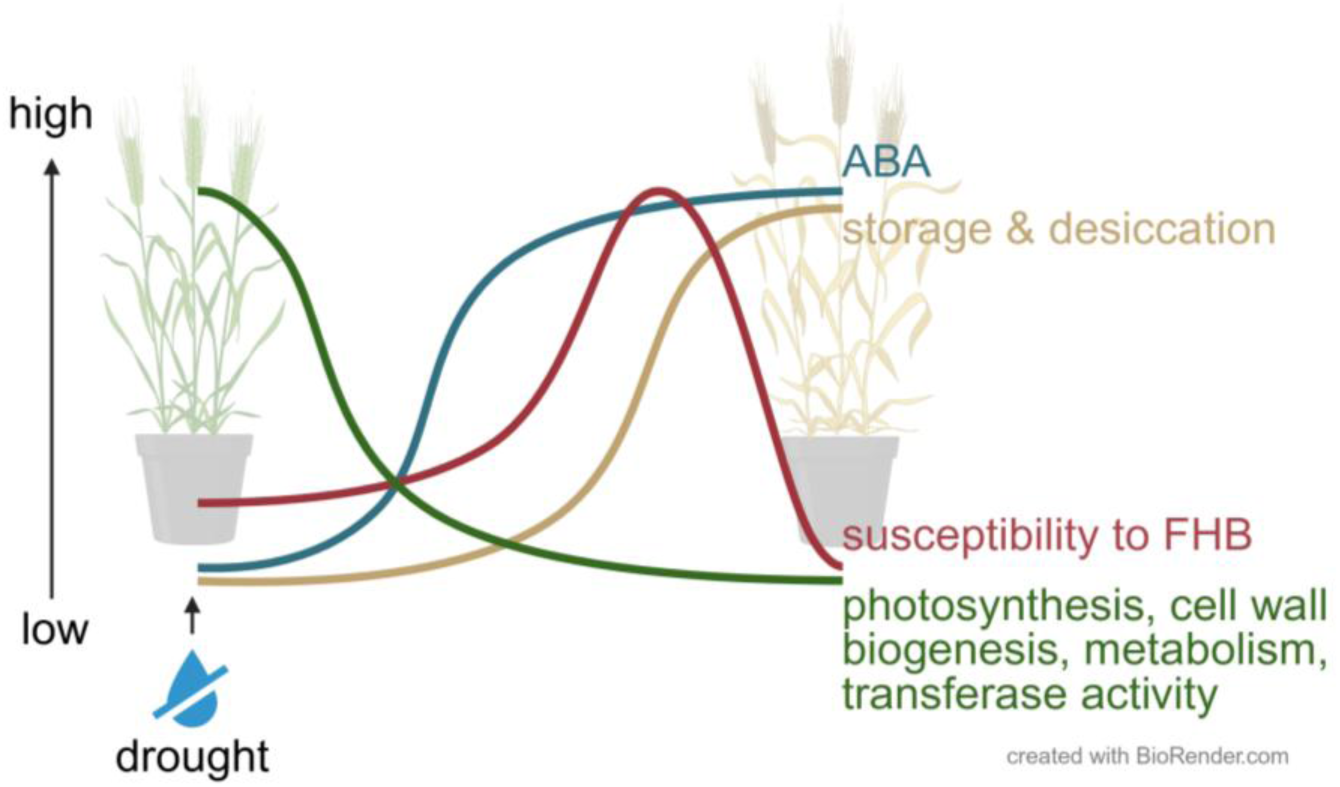

## Introduction

During the preparation of this study, the year 2024 marked a significant milestone, with the global average temperature exceeding the pre-industrial baseline by 1.5 °C for one full year for the first time (Copernicus Climate Change Service 2025). This threshold is widely regarded as a critical limit for irreversible environmental damage (Doelman et al. 2019). Europe is warming faster than the global average (van der Schrier et al. 2013) experiencing more frequent droughts (Hanel et al. 2018; Spinoni et al. 2018), which significantly threaten cereal production (Naumann et al. 2015; Brás et al. 2021; Martin et al. 2025). Simultaneously, climate change is intensifying plant disease dynamics by altering pathogen abundance, distribution, and host susceptibility (Chakraborty and Newton 2011; Delgado-Baquerizo et al. 2020). Rising temperatures, air humidity, and atmospheric CO₂ levels are driving changes in pathogen abundance, geographic distribution, and host susceptibility, creating increasingly complex stress environments and weather events. For pathogenic *Fusarium* spp., climate-driven shifts in species and chemotypes are changing mycotoxin profiles, posing risks to food safety (Nnadi and Carter 2021; Johns et al. 2022).

While the individual impacts of drought and pathogens on plants are well studied, their simultaneous occurrence triggers complex and often unpredictable plant responses (Gupta et al. 2016; Zandalinas et al. 2021). These stress interactions involve signal cross-talk, resource trade-offs, and altered defense strategies (Karasov et al. 2017; Leisner et al. 2023). Even mild stresses, when combined, can have disproportionate effects on plant health and yield, depending on the timing, sequence, and dominance of each stressor (Pandey and Senthil-Kumar 2019; Zandalinas et al. 2021). This complexity highlights that the physiological and molecular mechanisms underlying plant responses to multiple stresses remain poorly understood. Although there is growing research on stress interactions, most studies focus on isolated stressors or use model plants, limiting agricultural relevance (Prasch and Sonnewald 2013; Zandalinas et al. 2021; Tan et al. 2023).

In cereals, drought alters host susceptibility (Liu and Liu 2016; Hoheneder et al. 2023) and pathogen dynamics of *Fusarium* spp., the causal agents of Fusarium Head Blight (FHB), making it difficult to predict disease pressure and complicating control strategies (Lahlali et al. 2024). This is particularly concerning in cereals, where FHB reduces yield and contaminates grain with harmful mycotoxins (Wegulo et al. 2015). Despite the evident risks, crop breeding efforts have rather neglected combined stress responses, likely because of their complexity.

Here, we investigated how simultaneous *Fusarium* infection and drought stress affect FHB severity, stress-related gene expression, and hormonal regulation in barley. We reveal a genotype-dependent increase in FHB severity under drought, accompanied by strong additive transcriptomic responses. Notably, we identified core sets of differentially expressed genes shared across individual and combined stress conditions, and used a linear regression model to predict gene expression under combined stress from single stress profiles.

## Materials and Methods

### Greenhouse experiment

Four spring barley genotypes (Avalon, Barke, Morex, and Palmella Blue) were cultivated under controlled glasshouse conditions at the Greenhouses and Phytochambers Unit of the Plant Technology Center, Technical University of Munich. These genotypes showed varying levels of quantitative resistance to FHB previously determined in field (Hoheneder et al. 2022) and greenhouse experiments (Hoheneder et al. 2023).

Five grains were sown per 3-litre pot, each containing 900 g of peat substrate (Einheitserde C700, Stender, Germany). 18 pots per genotype and treatment group ensured an adequate number of individual plants. Supplemental lighting for 16 hours per day provided long-day conditions. The temperature was 18 °C during the day and 16 °C at night, with a relative humidity of 60%.

Drought stress was applied to half of the pots starting from mid flowering stage (growth stage 65) by a stop of automatic daily watering. These drought-stressed pots were weighed to monitor water loss at 0, 1, 2, 4, and 8 days post-watering cessation (dpw) and manually supplemented with 75 ml of water at 4 dpw and 50 ml at 8 dpw. Severe drought stress was achieved by reducing the relative pot weight to 40% after 8 dpw until growth stage 90 (Fig. S1). Plants under drought conditions visually exhibited reduced turgor pressure and partial loss of green leaf area, while the watered plants displayed normal growth and ripening. At mid-flowering, half of the irrigated plants and half of those designated for subsequent drought stress were sprayed till run-off with either a *Fusarium culmorum* (Fc) spore solution or a mock solution as described in detail by (Hoheneder et al. 2023), resulting in the following treatments: WM (watered+mock), WFc (watered+infected), DM (drought+mock), and DFc (drought+infected). At 2, 4, and 7 dpi, four pots were randomly selected, and individual spikes (four per sample) were cut off and flash-frozen in liquid nitrogen, obtaining four biological replicates.

### Preparation of *F. culmorum* inoculum and mock solution

The fungal *Fusarium culmorum* inoculum (isolates Fc002, Fc03, and Fc06; from the culture collection of the Chair of Phytopathology, Technical University of Munich) was cultured as outlined by (Linkmeyer et al. 2013). The spore solution was prepared as described by (Hoheneder et al. 2023).

### Measurement of phytohormones and metabolites

Spike samples (150 mg ground tissue, identical to those used for DNA/RNA extraction) were used to quantify abscisic acid (ABA), abscisic acid glucoside (ABA-Glc), phaseic acid (PA), dihydrophaseic acid (DPA), auxin (IAA), jasmonic acid (JA), salicylic acid (SA), salicylic acid glucoside (SA-Glc), and the osmolyte proline as described by (Chaudhary et al. 2020). Samples were supplemented with internal standards, extracted in ethyl acetate, and the filtered supernatant was analyzed by LC−MS/MS in multiple reaction-monitoring mode. Data were acquired using Analyst 1.6.3 (Sciex, Darmstadt, Germany).

Gaseous ethylene (ET) production in spikes was measured in four replicates per treatment at 2, 4 and 7 dpi. Four spikes per treatment were sampled in glass vials containing 1 ml of sterile water to prevent the spikes from desiccation by soaking in water. The vials were closed with gas-tight rubber lids and incubated for 5 hours under daylight at room temperature. 1 ml of the headspace was injected into a gas chromatograph (Varian Aerograph 3300, Shimadzu, Japan) equipped with a deactivated aluminium oxide column and combined with an Integrator (C-R6A Chromatopac, Shimadzu, Japan). ET amounts (pmol/ml air) were calculated by dividing peak areas by 214 (internal calibration: 1 ml pure ethylene gas = 214 pmol at 1 bar) (Kruedener et al. 1995) and normalized to spike fresh weight.

### Area under the curve analyses

Measured phytohormone and proline quantities were used to calculate the area under the curve (AUC, 4-7 dpi) following (Shaner and Finney 1977). Similarly, area under the disease progression curve (AUDPC) values were calculated for FHB symptoms and fungal DNA in infected and drought-plus-infected samples. Log_2_ fold changes of AUCs were computed relative to watered mock-treated plants (WFc, DM, DFc). Hierarchical clustering (average linkage) grouped phytohormone responses across cultivars and treatments using MultiExperiment Viewer 4.9.0 (Howe et al. 2011).

### Measurement of chlorophyll contents and stomatal conductance

Physiological responses were assessed by measuring chlorophyll content and stomatal conductance of upper leaves (flag leaf [F] and flag leaf-1 [F-1]). Chlorophyll was recorded as SPAD values on 18 leaves per stage at 2, 4, and 7 dpw using a SPAD502 meter (Konica Minolta, Japan). Stomatal conductance was measured on 15 leaves per stage at 3, 5, and 8 dpw using an SC-1 Leaf Porometer (Meter Group, Germany).

### DNA and RNA extraction

Spike samples were ground in liquid nitrogen for DNA extraction following (Fraaije et al. 1999). DNA was diluted to 20 ng/μl. RNA was extracted using the GeneMATRIX Universal RNA Purification Kit (EURx, Poland) with DNA digestion, quantified with a Qubit 2.0 fluorometer, and diluted to 100 ng/µl.

### Quantification of fungal DNA

*F. culmorum* and barley DNA in spike samples were quantified using the qPCR protocol and species-specific primers from (Nicolaisen et al. 2009) as described in detail by (Hoheneder et al. 2023).

### Library preparation and 3’RNA-sequencing

Libraries for 3’-RNA sequencing were prepared using the QuantSeq 3’mRNA-Seq Library Prep Kit (Lexogen, Austria) following the manufacturer’s protocol (Moll et al. 2014). Library concentrations were quantified with a Qubit 2.0 and Qubit DNA HS Assay, and pool concentrations determined by qPCR. Library integrity and fragment size were assessed with an Agilent 2010 Bioanalyzer (HS DNA Assay). Sequencing was performed on an Illumina NovaSeq 6000.

### Read quality check, trimming, mapping and annotation

3’mRNAseq reads were processed with nf-core/rnaseq (3.17.0) using Nextflow (24.04.4) (Ewels et al. 2020). The quality of the raw reads was checked using FastQC (0.12.1), adapter trimming and filtering was performed with Trim Galore (v0.6.10) and Cutadapt (4.9). STAR (2.7.11b) was used for read alignment against the barley reference genome Morex v3 (Mascher et al. 2021) merged with the *F. culmorum* genome FusCulm_01 (GenBank assembly GCA_003033665.1) and Salmon (1.10.3) for transcript quantification. The 3’UTR regions of each gene were extended by 3 kb or until the next gene to improve mapping rates.

### Differential gene expression analysis and WGCNA

Differential gene expression analyses were performed with edgeR v4.0.16 (Robinson et al. 2010). For each time point (4, 7 dpi), differential expression was calculated as variety_condition – variety_watered-mock, where variety = Avalon, Barke, Morex, or Palmella Blue and condition = watered-infected, drought-mock, or drought-infected, yielding 15,094 DEGs. Significant DEGs were filtered by FDR-corrected p < 0.01 and log_2_FC (log_2_FC < 0: down-regulated; log_2_FC > 0: up-regulated) (Fig. S2). Co-expression analysis was performed using the WGCNA R package (Langfelder and Horvath 2008, 2012) on 6,558 significant DEGs, using their normalized counts per million as input. Modules of co-expressed genes were identified and correlated with phenotypic traits (quantified phytohormones, proline, fungal DNA contents and FHB symptoms; see Suppl. Dataset_P). Network construction used the following parameters: blockwiseModules(mydatExpr, power = 8, TOMType = “signed”, minModuleSize = 30, reassignThreshold = 0, mergeCutHeight = 0.25, numericLabels = TRUE, pamRespectsDendro = FALSE, saveTOMs = TRUE, saveTOMFileBase = “mydatTOM”, verbose = 3).

### DEG Cluster analysis

To cluster DEGs by expression pattern across treatments, a self-organizing tree algorithm (SOTA) was computed in MultiExperiment Viewer v4.9.0 (Howe et al. 2011) using 6,558 significant DEGs (FDR-corrected p < 0.01; |log_2_FC| > 1) in at least one genotype.

### Gene Ontology enrichment analysis

Gene Ontology (GO) enrichment analysis was performed using g:Profiler (Kolberg et al. 2023) with default parameters, except term size, which was limited to at least 5 genes. The most enriched terms for biological process (BP), molecular function (MF), and cellular component (CC) were visualized using ‘ggplot2’ and ‘dplyr’ in R Studio. The visualizations display the most significant GO terms (based on the -log10 adjusted p-value), the respective gene counts (DEGs per GO term), and gene ratios (DEGs per GO term/total DEGs).

### Similarity Score analysis

To compare gene sets regulated under single or combined stress, a Similarity score (Jaccard index) was calculated following (Tan et al. 2023) as the ratio of the intersection to the union of two DEG sets. Therefore, Venn diagrams for up- and down-regulated DEGs per treatment, genotype, and time point were generated (Fig. S3) using Venny 2.1.0 (Oliveros 2007).

### Similarity of gene expression profiles under single and combined stress

To assess similarity of gene expression profiles under combined stress (log_2_FC DFc) versus individual stresses (log_2_FC WFc, log_2_FC DM), absolute differences |log_2_FC DFc − log_2_FC WFc| and |log_2_FC DFc − log_2_FC DM| were calculated for each DEG. Each gene was assigned to the stress with the smaller difference, and a color-coded scatterplot was generated in R studio (‘ggplot2’) to visualize the similarity. This identified whether a gene’s expression under combined stress resembled one of the individual stresses more closely.

### Multiple Linear Regression analysis

Multiple Linear Regression (MLR) models were defined to predict gene regulation under combination stress based on the assumption of additive effects from individual stresses, as observed in this study and previously (Hoheneder et al. 2023). The additive model was complemented with a two-way interaction term between single stresses, accounting joint effects on gene expression under stress-combination, which improved the prediction accuracy.

Gene expression datasets were compiled for: all regulated DEGs (n = 15,094), significantly regulated DEGs under combined stress per genotype (n = 914–4,700), and DEGs significantly regulated in at least one genotype (n = 6,558). For each DEG, average log_2_FC under combined stress was predicted from the averages under individual stresses. Across all datasets and models, multicollinearity was tested between independent variables, indicating the degree to which the variances in the regression estimates are increased due to high correlation between independent variables. Variance inflation factors remained low (< 3.5) among the tested models and datasets. Model selections were based on the goodness of fit (R^2^) and comparison of the Akaike information criterion as an estimator of prediction error of different models. Homoscedasticity was checked in residual plots. Quantile-quantile plots were generated to graph the distributions of residuals. Normality was not strictly required, as the analysis involved large gene expression datasets and linear models are robust to mild distributional deviations under the Central Limit Theorem (Lumley et al. 2002). Differences between R^2^ and adjusted R^2^ values were compared, indicating model overfitting. The regression model maximizing R^2^ and minimizing the Root Mean Square Error has the following equation:

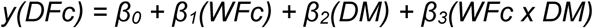

Where:

y(DFc) = predicted expression under combined stress β₀ = intercept

β₁(WFc) = effect of infection β₂(DM) = effect of drought

β₃(WFc × DM) = interaction between infection and drought.

Regression coefficients (β₀–β₃) and corresponding p-values were extracted to estimate the contribution of each predictor to gene expression under combined stress.

### Data availability

All data supporting the findings of this study are available throughout the paper and online accessible: #available when published. The raw sequencing data has been deposited at the NCBI Gene Expression Omnibus (accession GSE310570) and will be available when published.

## Results

### Drought stress supports FHB disease in moderately resistant cultivars

To understand better the quantitative FHB resistance of barley under complex stress, the present study determined the differences in FHB severity and stress responses under simultaneous drought stress and infection by *F. culmorum* (DFc) in comparison to single stresses (WFc, watered *Fc*-infected; DM, drought-stressed mock-infected) and non-stressed controls (WM, watered mock-infected; Fig. 1a).

**Figure 1:**
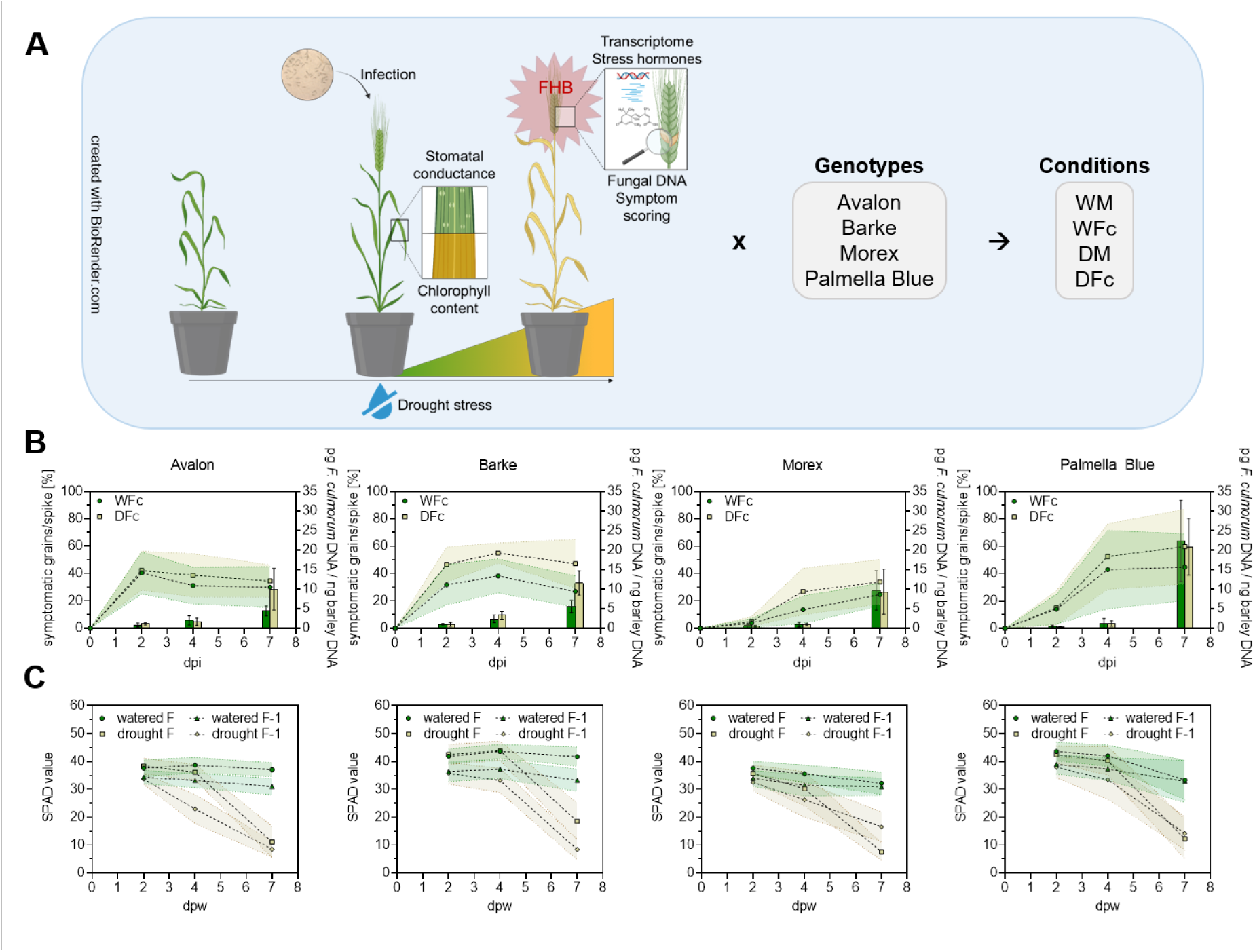
Genotype-Specific Responses to Drought and Infection. (A) Barley cultivars Avalon (Av), Barke (Ba), Morex (Mo), and Palmella Blue (Pb) were grown under controlled greenhouse conditions. Drought stress was applied by stopping irrigation at mid-anthesis. Plants were either watered (W) or drought-stressed (D) and inoculated with *F. culmorum* (Fc) or mock (M). Disease symptoms were recorded at 2, 4, and 7 dpi; spikes were collected for fungal DNA, transcriptome, and hormone analyses. Leaf chlorophyll was measured to monitor ripening and physiological responses. (B) FHB severity (% symptomatic grains per spike) and fungal DNA contents. Progression curves show symptom means; shaded areas indicate SD. Bars display mean fungal DNA at 2, 4, and 7 dpi with SD. (C) Mean SPAD (n = 18, F and F-1 leaves) from 2–7 dpw in watered and drought-stressed plants; shaded areas show SD. Differences between treatment groups were tested using Mann-Whitney U-test (table S1-3).

We assessed FHB severity visually and via fungal DNA quantification in the four barley cultivars (Fig. 1a, b). Both watered and drought-stressed plants showed infection by 2 dpi, with increasing symptoms and fungal DNA. From 4 dpi onwards, drought stress enhanced FHB symptoms, notably in Palmella Blue, Barke, and Morex, with visual differences most pronounced at 7 dpi. Avalon showed only a slight symptom increase under drought, but fungal DNA content rose significantly (*p = 0.0286). Barke displayed a strong symptom increase (**p = 0.0019) and higher fungal DNA (*p = 0.0286) under drought. Morex and Palmella Blue showed no DNA differences at 7 dpi, with Palmella Blue being the most susceptible (table S1-2).

Chlorophyll contents and stomatal conductance reflected plant ripening and drought response. Irrigated plants maintained stable chlorophyll, whereas SPAD-values of flag leaf and flag leaf-1 decreased significantly under drought (****p < 0.001; Fig. 1c), reflecting accelerated senescence. Transpiration declined under limited water, too, indicated by reduced stomatal conductance (Fig. S4).

### Plant hormone contents reflect drought, FHB responses and gene expression patterns

To examine pathogenesis-related and physiological changes with regard to quantitative FHB resistance in more detail, we measured hormones and metabolites by LC-MS/MS or gas chromatography. Global gene expression was measured by 3’RNA-sequencing detecting 50,267 different gene transcripts from spike tissues. We performed hierarchical clustering of log_2_ fold changes in AUC values for phytohormones measured between 4–7 dpi, alongside AUDPC values for fungal DNA and visible symptoms from 0–7 dpi. Pearson correlations visualized relationships between plant hormones and disease severity (Fig. 2a, b). WGCNA clustered 6,558 DEGs (|log_2_FC| > 1; FDR-corrected p < 0.01) into 18 co-expression modules to identify associations with measured physiological traits (Fig. 2c).

**Figure 2:**
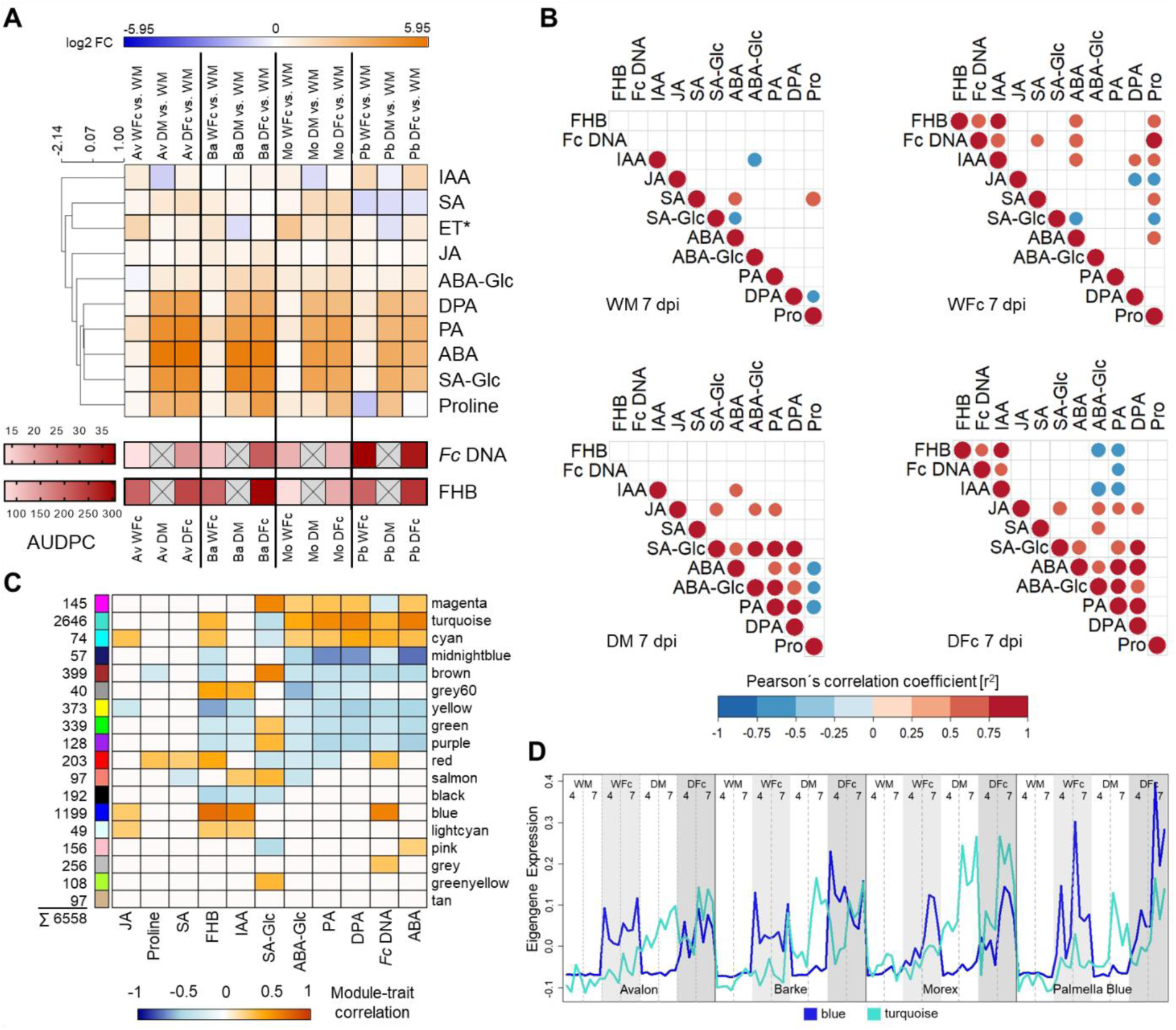
Phytohormone regulation, correlation and WGCNA in drought- and pathogen-stressed barley. (A) log_2_ fold changes in phytohormone levels (AUC from 4 to 7 dpi) for WM vs. WFc, DM, and DFc treatments. Hierarchical clustering highlights treatment-specific hormone responses. The respective AUDPC values for FHB symptoms and fungal DNA indicate disease severity across cultivars. *ET: The gaseous ethylene was measured on additional spikes sampled in parallel to those used for fungal DNA and phytohormone quantification. (B) Pearson correlation matrices showing significant (p < 0.05) relationships among phytohormone levels, symptomatic grain proportions, and fungal DNA at 7 dpi. (C) WGCNA module-trait relationships with significant eigengene correlations (coloured boxes: FDR-corrected p < 0.01) to measured stress parameters (D) Line plots of eigengene expression profiles for the two largest modules (turquoise, blue) across genotypes, treatments, and time points.

Hierarchical clustering grouped several drought-responsive metabolites showing a positive log_2_ fold change under drought or combined stress. The strong positive regulation of ABA-Glc, DPA, PA, ABA, SA-Glc and proline reflects the accumulation of these stress-related hormones/metabolites, which confirms the successful application of drought stress in all four genotypes (Fig. 2a; Fig. 1c, Fig S4). Avalon and Barke showed highest regulation of ABA under drought and combination stress, but also highest absolute values, demonstrating that these genotypes most strongly responded to the drought (Fig. 2a, Fig. S5). The stress hormones SA, ET and JA showed diverse patterns or slight up- or down-regulation, depending on individual stress treatments and genotypes. In addition, auxin showed down-regulation under drought stress in Avalon, Morex and Palmella Blue. In contrast, infection and drought plus infection led to an up-regulation of auxin. This was most pronounced in susceptible Palmella Blue and hardly detectable in less susceptible Barke, in which slightly positive fold changes of auxin remained relatively uniform across all treatments. In addition, the AUDPC integrated the progression of FHB symptoms and fungal DNA contents to display overall effects of drought stress or irrigation on disease severity (Fig. 2a).

Pearsońs correlations suggested associations between multiple stress-associated parameters over all genotypes (Fig. 2b). Under control conditions at 7 dpi, phytohormones show a positive correlation between ABA and SA (r^2^ = 0.80) and a negative correlation with SA-Glc (r^2^ = -0.53). Proline and SA were in a positive relationship (r^2^ = 0.72), while proline and DPA were negatively correlated (r^2^ = -0.52). In infected samples, auxin showed a positive correlation with the scored *Fusarium* symptoms and fungal DNA contents in WFc or DFc at 7 dpi (Fig. 2b) and 4 dpi (Fig. S6). Similar correlations were also found when taking all infected samples into account (WFc+DFc, 4+7 dpi) (Fig. S6). The correlations suggest a strong relationship between auxin levels and FHB infection. Strong positive correlations between proline and *Fusarium* symptoms (r^2^ = 0.6) or fungal DNA contents (r^2^ = 0.8) were also found in watered plants at 7 dpi (Fig. 2b). Auxin and ABA positively correlated under infection or drought stress alone, but this was resolved under combined stress. In line with the positive regulations of ABA, ABA-Glc, PA and DPA under drought stress (Fig 2a), ABA and ABA-Glc positively correlated with ABA-catabolites PA and DPA under similar conditions (DM, DFc, 7 dpi), but not under infection stress (WFc, 7 dpi; Fig. 2b). A similar pattern was found at 4 dpi (DM, DFc; Fig. S6), which confirms the close relationship of these hormones with drought.

We carried out a WGCNA to associate gene co-expression modules with stress-associated phytohormones and disease parameters (Fig. 2c). The turquoise co-expression module revealed the strongest positive associations with ABA levels (r^2^ = 0.66), DPA (r^2^ = 0.64), PA (r^2^ = 0.60) and ABA-glucoside (r^2^ = 0.46), but also with *Fusarium* symptoms (r^2^ = 0.37) and fungal DNA contents (r^2^ = 0.38). The respective eigengene expression profile supports the association with drought stress by increased expression values in all DM and DFc samples (Fig. 2d). A Gene ontology enrichment analysis for 2646 DEGs in the turquoise module significantly enriched ‘carbohydrate metabolic process’, ‘catabolic process’, ‘response to water’, ‘response to acid chemical’, and further ‘response to water deprivation’ and ‘response to endogenous stimulus’ as enriched biological processes reflecting the drought stress. In addition, ‘phosphoric ester hydrolase activity’, ‘hydrolase activity’, ‘phosphatase activity’, and ‘nutrient reservoir activity’ were the most significantly enriched molecular functions, indicating stress responses, metabolism and signal regulation (Fig. S7). In support of this, the list of DEGs in the turquoise module contains numerous stress- and maturity-related functions, namely 16 ‘peroxidases’, seven ’abscisic-associated and receptors proteins’, 16 ‘stress-’ and 15 ‘senescence’-related, nine ‘dehydration’, 34 ‘late embryogenesis abundant’ and nine ‘seed maturation’ genes (supplemental Dataset W17). Hence, the turquoise module represents a large drought-responsive gene cluster, reflecting stress-accelerated ripening and ABA regulation (Fig. 2b).

The second largest blue module (1199 DEGs) showed strong positive associations with FHB symptoms (r^2^ = 0.74), fungal DNA (r^2^ = 0.64), auxin (r^2^ = 0.63) and to a lesser extend with JA (r^2^ = 0.27). The respective eigengene expression profile showed strong up-regulation of genes in the infected (WFc, DFc) samples across all cultivars (Fig. 2d). This demonstrates a close module association with pathogenesis and infection stress. The GO analysis identified ‘glutathione metabolic process’ as the most significant biological process, and ‘glutathione transferase activity’, ‘transferase activity, transferring alkyl or aryl (other than methyl) groups’, ‘transferase activity’ and ‘glycosyltransferase activity’ as the most enriched molecular functions (Figure S8). Since the expression pattern of almost all DEGs in the blue module showed an up-regulation under infection and double stress and often no or down-regulation under drought stress (Dataset W2), the module predominantly contains typical infection-responsive DEGs with high gene-trait associations (see ‘gene-trait-significance’ values; Dataset W2). Prominent examples are a disease resistance protein (TIR-only class, HORVU.MOREX.r3.2HG0134190) and 33 ‘Glutathione S-transferases’ (e. g. HORVU.MOREX.r3.5HG0424890; HORVU.MOREX.r3.7HG0665160) as described to be involved in DON-detoxification (Gardiner et al. 2010). In addition, the DON-detoxifying UDP-glycosyltransferase *Hv*UGT13248 (HORVU.MOREX.r3.5HG0464880) (Schweiger et al. 2010; Mandalà et al. 2019) was highly expressed under infection and combination stress, but not under drought. *Hv*UGT13248 was already previously identified as highly *Fusarium*-responsive hub-gene with a stable expression under both infection and drought-pre-stressed barley infection (Hoheneder et al. 2023). In addition, eight ‘2-oxoglutarate (2OG) and Fe(II)-dependent oxygenase superfamily proteins’ showed high regulations under infection and combination stress. These genes represent putative immune regulators facilitating *Fusarium* susceptibility (Low et al. 2020). With regard to defence, eleven ‘pathogenesis-related proteins’ (including ten PR-1), six ‘glucanases’, seven ‘chitinases’, four ‘thaumatin-like proteins’ and ten ‘peroxidases’ were strongly up-regulated under infection (WFc, DFc) displaying high basal defence activity (Dataset W2).

The brown module (399 DEGs) positively correlated with SA-glucoside (r² = 0.64) and negatively with fungal DNA, FHB symptoms, ABA and its derivatives (ABA-Glc, PA, DPA), and proline (Fig. 2c). GO terms indicate general pathogen and drought stress regulation, including ‘regulation’ and accumulated several ‘metabolic processes’ (Fig. S9). The yellow module reflects drought effects on photosynthesis and energy supply (Fig. 2c; Fig. S10; Dataset W18). The red module correlated with FHB symptoms, fungal DNA, proline, and SA (Fig. 2c). Magenta and cyan modules showed patterns similar to turquoise, except for JA or SA-Glc, while midnightblue displayed opposite associations (Fig. 2c). The magenta eigengene expression distinguished Avalon and Barke from Morex and Palmella Blue, reflecting genotype-specific regulation (Fig. S11).

Taken together, the different modules indicate that barley employs diverse gene sets with distinct functions to regulate complex stresses, each linked to stress-specific hormonal regulations or disease severity.

### Drought and FHB additively contribute to combined stress responses

We compared global trends of gene expression under individual and combined stress conditions across cultivars and time points (Fig. 3a; Fig. S2). DEG comparisons using the Jaccard Index (Fig. 3b) revealed quantitative differences between treatments, with combined stress inducing the highest number of DEGs (4 dpi: 918–1454; 7 dpi: 3780–4704), reflecting stress severity indicated by increasing fungal DNA and ABA levels (Fig. 1b, 2a). Drought stress alone induced the second highest DEG numbers at 7 dpi (2299–3363), with a strong and consistent increase from 4 dpi (317–1405) across cultivars.

**Figure 3:**
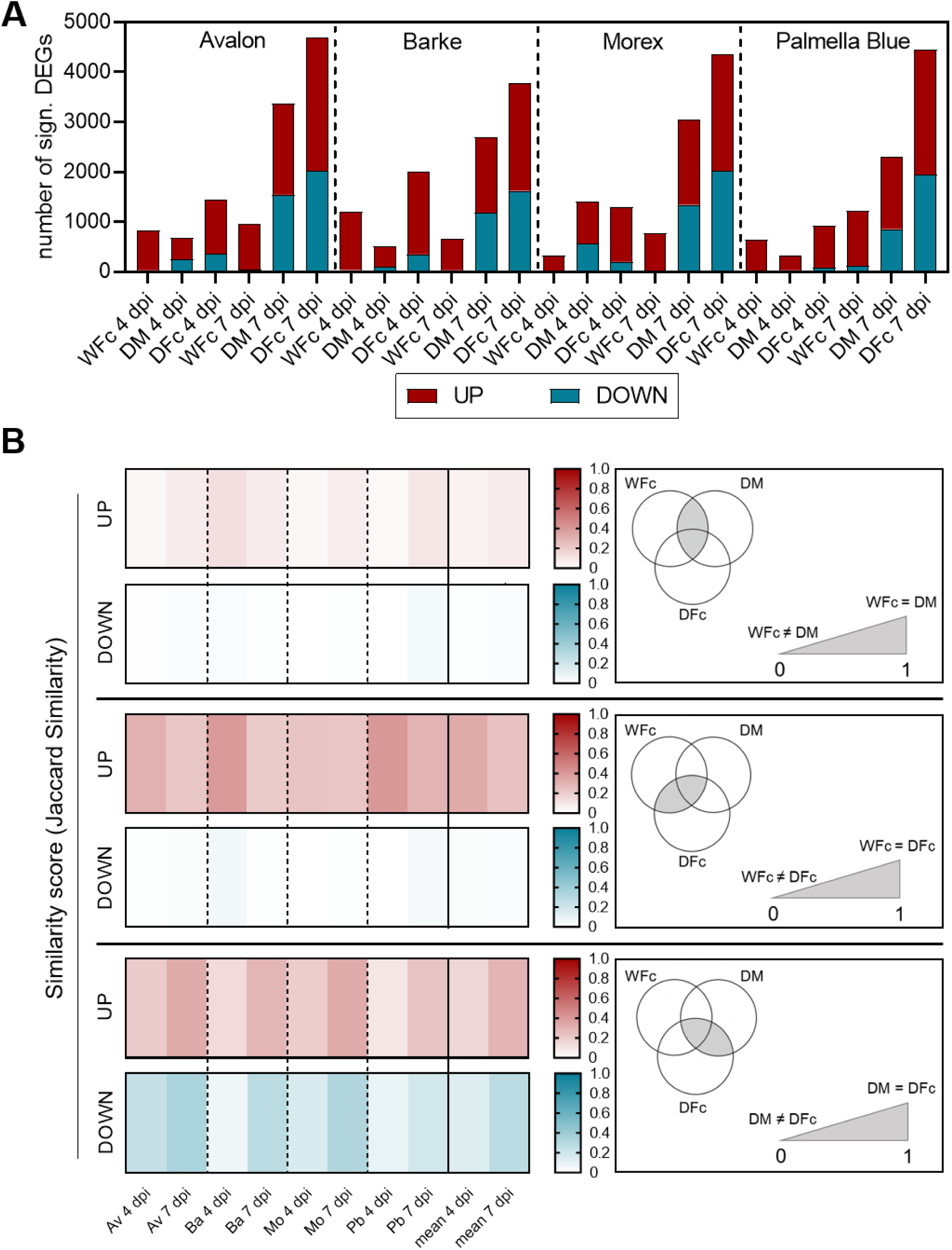
Significant DEG numbers and similarity scores between gene sets regulated under single or combination stress. (A) Significantly regulated DEGs (FDR-corrected p < 0.01) across genotypes and conditions at 4 and 7 dpi. Red bars: up-regulated; blue bars: down-regulated DEGs. (B) Similarity scores between DEGs under single and combined stress across genotypes and time points, calculated separately for up- and down-regulated genes. Venn diagrams show overlapping gene sets corresponding to the similarity scores.

Across treatments, more genes were up- than down-regulated. Infection alone led to several hundred up-regulated DEGs per cultivar but relatively few down-regulated DEGs (max. 113 in Pb at 7 dpi). In contrast, drought and combined stress caused substantial down-regulation of DEGs, increasing from 4 dpi (DM: 31–559; DFc: 81–357) to 7 dpi (DM: 855–1541; DFc: 1624–2019), accounting for nearly half of total DEGs under these conditions.

Combined stress generally followed an additive pattern: up-regulated DEGs largely matched the sum of single stress responses, while down-regulated DEGs exceeded additive expectations, except in Morex at 4 dpi (Fig. 3a, Fig S2). DEG similarity between infection and drought stress remained low (Fig. 3b), indicating separated transcriptional responses.

Infection and combined stress shared on average 29% of up-regulated DEGs, with a time-dependent decrease in overlap from 4 to 7 dpi across most cultivars. Overlap in down-regulated DEGs was minimal due to the weak down-regulatory effect of infection alone (Fig. 3b). In contrast, DEG overlap between drought and combined stress increased toward 7 dpi across all cultivars, reflecting both the rising number of drought-responsive genes and their consistent expression under co-occurring FHB-stress (Fig. 3b; Dataset D1).

### Core stress response genes are stably expressed under single and combined stress

To assess whether and how single stress responses account for gene expression under combined stress, we subtracted single stress regulation from combined stress regulation values and visualized all DEGs (Fig. 4). In the corresponding similarity plots, DEGs near the y-axis reflect infection-like expression under combined stress, those near the x-axis reflect drought-like expression, and those near the diagonal indicate expression values under combined stress that are less well explainable by single stress effects. Figure 4a shows that many DEGs actually plot close to the y- axes at 4 dpi and thus their regulation can be largely explained by pathogenesis-related expression patterns. At 7 dpi, more genes plot close to the diagonal and the x-axis, which can be explained by additive or solely drought-related expression patterns, which is likely caused by the increasing drought severity (Fig. 5b). DEGs from Avalon or Morex show stronger drought effect already visible at 4 dpi (Fig 4a).

**Figure 4:**
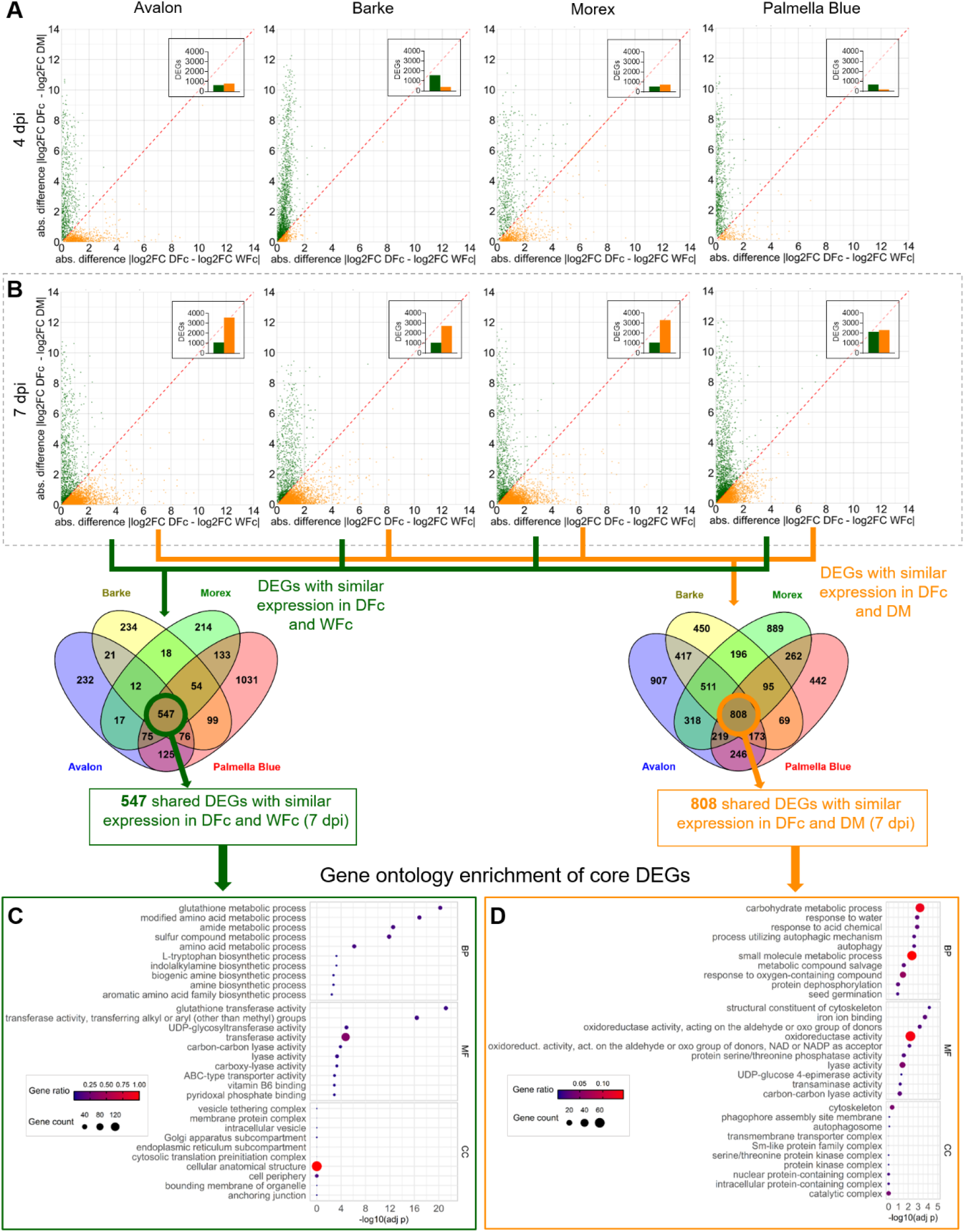
Expression Similarities and GO enrichment analyses of shared core DEGs. (A) Expression similarity plots for significantly (FDR-corrected p < 0.01) regulated DEGs at 4 dpi and (B) at 7 dpi. DEGs with similar expression under infection and combined stress are shown as green dots; those with similar expression under drought and combined stress are shown in orange. Bar graphs within each plot indicate total DEG counts. Venn diagrams show the number of shared DEGs with similar expression between DFc and WFc or DFc and DM across all four genotypes at 7 dpi. (C) GO enrichment for DEGs with similar expression in DFc and WFc. (D) GO enrichment for DEGs with similar expression under DFc and DM for biological process (BP), molecular function (MF) and chemical component (CC).

**Figure 5:**
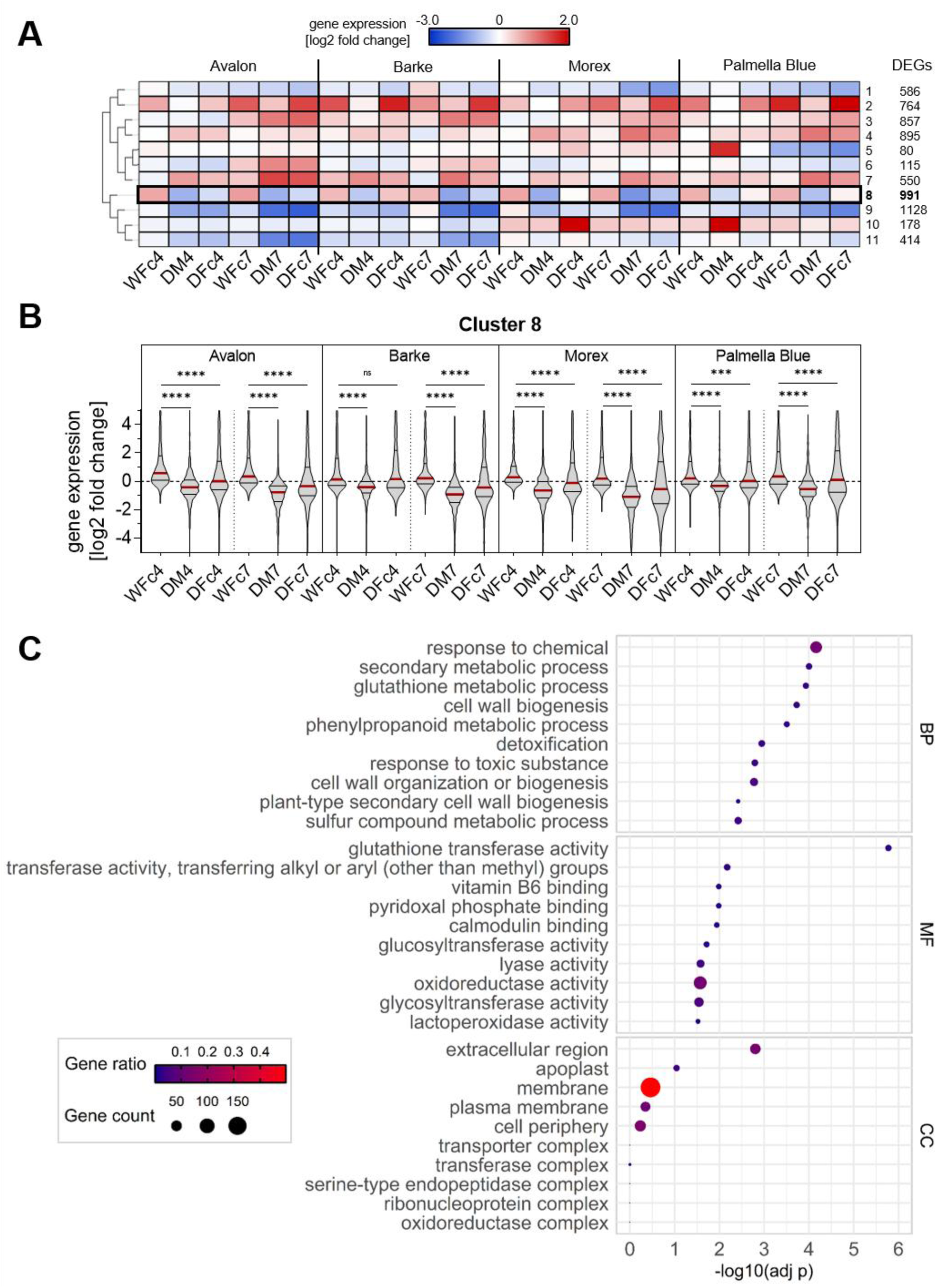
DEG and GO enrichment analyses targeting *Fusarium*-responsive DEGs under infection stress and weaker regulation under combination stress. (A) 6,556 significantly (FDR-corrected p < 0.01) regulated DEGs with |log_2_FC| > 1 sorted by a self-organizing tree algorithm (SOTA) into 11 clusters with similar expression pattern. The heat map displays the mean log_2_ fold changes across all genotypes, treatments and time points per each cluster. (B) Violin plots represent the frequency distribution of DEGs per treatment, time point and genotype found in cluster 8 according to their magnitude of gene expression. Violins show the median (red solid line) and upper and lower quartile (black lines). For better visualization, the data within a range of |log_2_FC| < 5 is shown. Significant differences between medians were tested with Kruskal-Wallis test and Dunńs multiple comparisons test at a significance level of 0.05. (ns = not significant; ***p < 0.001; ****p < 0.0001). (C) GO enrichment for DEGs found in cluster 8 for biological process (BP), molecular function (MF) and chemical compound (CC). GO enrichment analyses of the remaining clusters are shown in supplementary figures S13-22.

To identify robust and general stress responses, we intersected DEGs showing similar expression between DFc and WFc or DFc and DM in all genotypes. This revealed 547 core DEGs related to infection and 808 drought-related DEGs, shared across single and combined stress conditions (Fig. 4c, d). Among infection-related DEGs, enriched GO terms at 4 or 7 dpi, respectively, included ‘sulfur compound metabolic process’ and ‘glutathione metabolic process’, underscoring the role of glutathione S-transferase metabolism in detoxification during *Fusarium*- or DON-induced stress (Gardiner et al. 2010; Gullner et al. 2018) (Fig. 4c; Fig. S12). Enriched molecular functions like ‘transferase activity’, ‘glutathione transferase activity’, and ‘UDP-glycosyltransferase activity’ further highlight detoxification-related enzyme activity. Additionally, the enriched GO term ‘L-tryptophan biosynthetic process’ support the role of the tryptophan-derived defense pathway in barley (Ishihara et al. 2017; Powell et al. 2017; Hein et al. 2025) and its involvement in auxin biosynthesis linked to *Fusarium*-susceptibility (Brauer et al. 2019; Su et al. 2021). Highly expressed genes in these DEG sets include ‘glutathione S-transferases’, ‘tryptophan decarboxylases’, ‘Fusarium resistance orphan protein’, and ‘disease resistance protein’ (Dataset E1-2).

Among 51 (4 dpi; Dataset E3) and 808 (7 dpi; Dataset E4) DEGs with similar expression under DM and DFc, several were linked to drought, senescence, and maturation (e.g. ‘dehydrins’, ‘late embryogenesis abundant proteins’, and ‘ripening-related proteins’), supported by GO enrichment indicating a basal drought response. The early time point (4 dpi; Fig. S12) enriched for signaling-related processes (‘dephosphorylation’, ‘phosphatase activity’) (Yuan et al. 2016; Zhang et al. 2023), while later responses (7 dpi; Fig. 4d) shifted toward important drought stress adaptation mechanisms including ‘response to water’, ‘response to oxygen-containing compound’, and ‘autophagy’ (Liu et al. 2009; Tang and Bassham 2022).

Thus, the similarly regulated genes represent core sets of likely important stress-responsive genes expressed across diverse cultivars in barley (Fig. 4c, d; Dataset E1-4).

### Drought response limits FHB-related gene up-regulation

Overall, DEG profiles under combined stress appeared largely additive, but this does not readily explain the drought-induced increase in FHB severity. Many *Fusarium*-responsive DEGs were up-regulated under infection but down-regulated under drought, suggesting drought-attenuated defence responses. To capture such patterns, we applied a self-organizing tree algorithm, clustering 6,558 significant DEGs (|log_2_FC| > 1; FDR-corrected p < 0.01; Dataset D2) across cultivars, conditions, and time points into 11 co-expression groups (Fig. 5a). Cluster 8 comprised 991 DEGs consistently induced by infection but suppressed under drought and largely reduced under combined stress, except in Barke at 4 dpi (Fig. 5a, b). This cluster included stress-related glycosyltransferases, glutathione S-transferases, tryptophan decarboxylases, and pathogenesis-related proteins (Dataset S8), with GO enrichment for ‘metabolism’, ‘secondary metabolic processes’, ‘glutathione transferase activity’, ‘detoxification’, and ‘cell wall biogenesis’. The attenuation of those infection-related responses likely contribute to increased susceptibility of Avalon and Barke under drought (Fig. 5c).

### MLR model explains combined stress responses from single stress responses

Since the number of regulated DEGs and fold changes increased with stress intensity and under combined stress, our study and previous data (Hoheneder et al. 2023) suggest that gene regulation follows an additive pattern (Fig. 3a, c). To test this, we applied an MLR model to predict gene expression under combined stress from single infection and drought responses (Fig. 6). We used different DEG sets to optimize model performance. While predictions based on all DEGs or DEGs significant in at least one genotype were acceptable, the best fit was achieved when focusing on DEGs significant under combined stress (FDR-corrected p < 0.01). The final model, y(DFc) = β0 + β_1_(WFc) + β_2_(DM) + β_3_(WFc × DM), achieved high goodness of fit (R² = 0.58–0.98; RMSE = 0.44–1.65) across genotypes (Fig. 6a; Dataset M9), showing that expression under combined stress can be well predicted from single stress responses.

**Figure 6:**
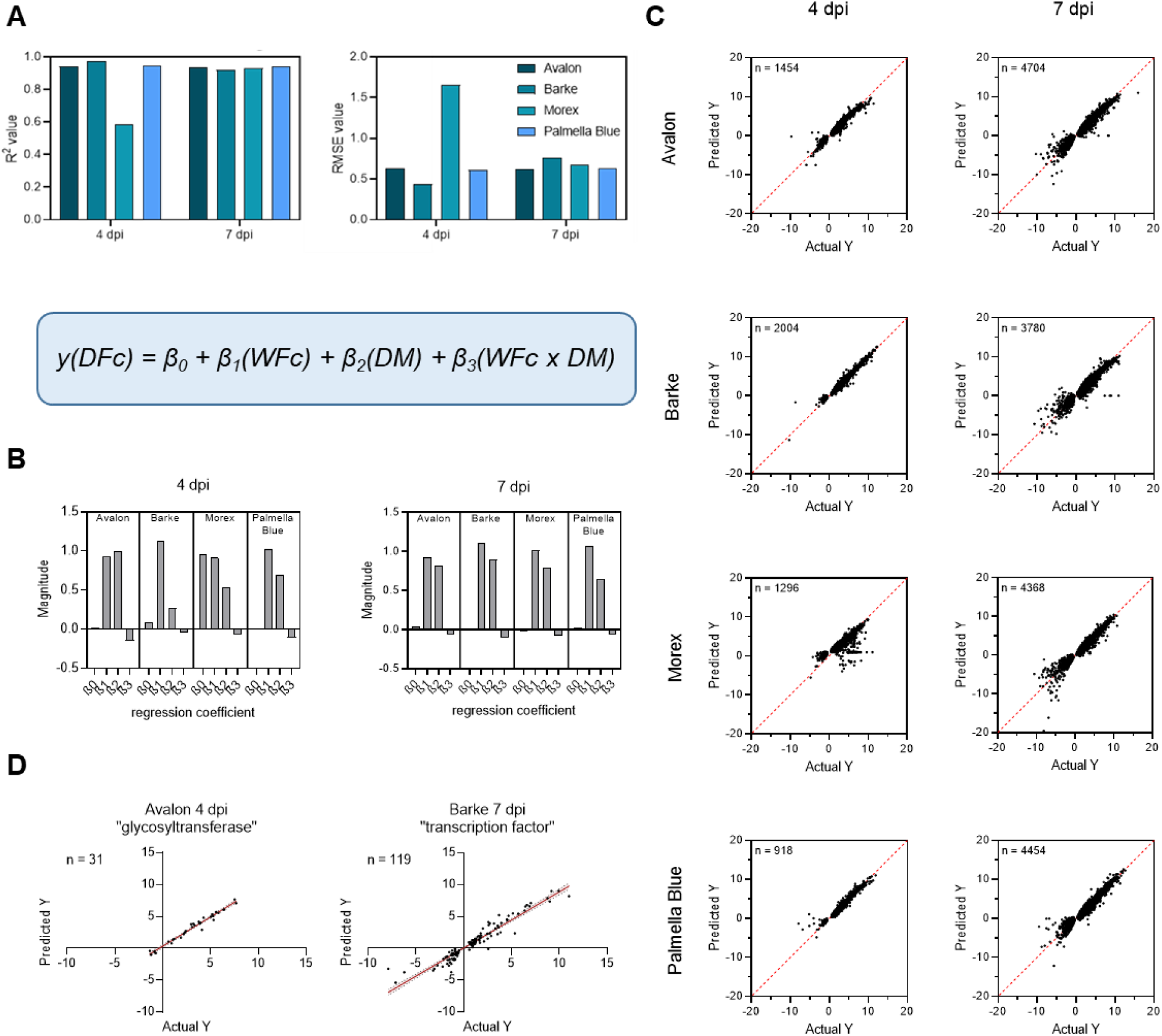
Multiple Linear Regression analysis for prediction of gene expression under combined stress. (A) Goodness of fit (R^2^) and respective Root Mean Squared Error obtained from the MLR model for significantly (FDR-corrected p < 0.01) regulated DEGs under combination stress per cultivar. The MLR model equation is given in the box. (B) Magnitude of regression coefficients for β_0_, β_1_ (main effect: WFc), β_2_ (main effect: DM) and β_3_ (interaction between WFc and DM) per cultivar and time point post-infection. (C) Actual and predicted log_2_ fold changes for significantly regulated DEGs in DFc obtained from the model for each cultivar and the two-time points post-infection. The dashed red line shows the line of identity. (D) Actual and predicted log_2_ fold changes in combination stress for exemplary gene sets selected by keyword search in the obtained DEG lists (Dataset M1-8) for ‘glycosyltransferase’ in Avalon 4 dpi and for ‘transcription factor’ in Barke 7 dpi. The red solid line displays the regression line, and dashed lines show the 95% confidence bands of the best-fit line.

Very high R² values were observed for Barke at 4 dpi (0.98) and all genotypes at 7 dpi (0.92–0.94). RMSE values remained below 0.76 except for Morex at 4 dpi, which showed lower accuracy (R² = 0.58; RMSE = 1.65) (Fig. 6a). Regression coefficients indicated strong positive effects of infection (β₁) and drought (β₂) on gene expression under combined stress. β₁ was high in Barke, Morex, and Palmella Blue at 4 dpi and in all cultivars at 7 dpi, while β₂ was greater than β₁ only in Avalon at 4 dpi and increased overall at 7 dpi, highlighting stronger drought influence at later times (Fig. 6b). All coefficients were significant (p < 0.0001), though β₀ was notably high in Morex at 4 dpi, reflecting lower model fit (Fig. 6b; Dataset M9).

The MLR model-predicted and measured log_2_ fold changes of DFc DEGs closely aligned with the identity line, confirming good model performance except for some deviations in Morex at 4 dpi (Fig. 6c). Exemplary regression predictions for selected gene sets, such as ‘glycosyltransferase’ in Avalon and ‘transcription factor’ in Barke, showed tight clustering around the regression line (Fig. 6d).

We applied the same model to published transcriptome data from a different experimental setup on *Fusarium*-infected and drought pre-stressed barley (Hoheneder et al. 2023). This yielded similarly high goodness-of-fit (R² = 0.90–0.97) and low RMSE (0.44–1.01), demonstrating robustness of the MLR model across barley cultivars and independent combined stress experiments (Fig. S23).

## Discussion

With more frequent extreme weather events due to climate change, the severity of stress affecting crops is gradually rising (Heino et al. 2023; Martin et al. 2025). Additionally, pathogen abundance and disease severity are likely increasing under these changing environmental conditions (Delgado-Baquerizo et al. 2020). This can result in more frequent exposure of plants to combined abiotic and biotic stresses. In this context, we assessed FHB severity and global gene expression responses in different barley genotypes under simultaneously increasing drought stress. Drought stress before infection reduced FHB severity in barley, likely due to accelerated ripening (Hoheneder et al. 2023). However, the complexity of plant stress regulations and the final outcome under combined stress cannot be generalized from individual experiments and largely depends on the genotype, stress severity, the timing and order of the stress applications (Suzuki et al. 2014; Zandalinas et al. 2021). The present study shows that simultaneous drought and *Fusarium* infection increase FHB severity, but this effect is depended on the plant genotype. For the transcriptome, we found a largely additive regulatory response under combined stress, showing an increasing dominance of drought-related regulation under lasting drought application. Additionally, we comprehensively measured multiple stress-related hormones and found associations with disease parameters and gene co-expression modules suggesting ABA and auxin regulation play important roles in limiting basal FHB resistance. A group of *Fusarium*-responsive genes showed trade-off effects under simultaneous drought (Fig. 5), which could have contributed to limited FHB resistance under drought.

Our data shows increased FHB severity in the two more resistant genotypes under drought. Barley regulated the highest number of genes under combined stress, indicating high physiological costs in complex stress situations (Fig. 2a, 2c, 3a). Hence, quantitative resistance might have been only partially functional under simultaneous drought stress, but this effect depended on the individual genotype (Fig. 1b). It should be noted that differences in flowering times between Avalon/Barke and Morex/Palmella Blue meant that simultaneous inoculation was not possible, thus limiting the full scope for data comparison. Nevertheless, all genotypes showed similar FHB severity at 4 dpi, and data show that the modern varieties Avalon and Barke are relatively robust and did not become fully susceptible under drought when compared to highly susceptible Palmella Blue. The drought-induced ABA levels were highest in these two genotypes (Fig. S5), thus they likely reacted strongly to drought. ABA was previously shown to promote FHB severity by interfering with defence signaling (Qi et al. 2016; Buhrow et al. 2021). However, ABA concentrations positively correlated with disease parameters under infection (Fig. 2b), but not under combined stress, likely because the upfront more susceptible varieties accumulated less ABA. Under double stress, ABA-glucoside, and phaseic acid showed negative correlations with FHB. In contrast, the more *Fusarium*-susceptible cultivars Morex and Palmella Blue (Hoheneder et al. 2022; Hoheneder et al. 2023) were strongly infected under irrigation, and drought did not further increase fungal DNA contents, although slightly more FHB symptoms were visible during drought in all genotypes. It is therefore possible that these two genotypes were less responsive to drought, and ABA regulation hence did not have a negative effect on basal resistance. However, we found several ABA-related and positively co-upregulated ABA-metabolites (ABA-Glc, PA, DPA) and SA-glucoside in drought stressed samples (Fig. 2a, b; Fig. S5), which may have influenced resistance, but their exact role in relation to FHB in barley is unclear and needs future investigation. Together, our data suggest an association of ABA with drought-supported FHB susceptibility during ongoing pathogenesis in Avalon and Barke.

Importantly, the two largest WGCNA co-expression modules (Fig. 2c, blue and turquoise) each associated with respective infection- or drought-stress-related stress parameters, respectively. Similarly, also smaller modules add additional stress- or genotype-specific gene clusters with specific links to stress-related hormone data. Examining DEG lists and GO terms enabled us to describe in detail the drought, infection, and combination stress responses and link gene sets to measured stress parameters in the four barley genotypes (Fig. 2c; Fig. S7-10; S24-37; Dataset W1-20). Overall, the DEG clustering and hormone data provide a global and detailed insight into barleýs response to combined stress, but do not explain increased FHB severity under drought in Avalon and Barke. Therefore, we required a detailed analysis of patterns of differentially gene expression.

The overall transcriptional response shows that additive effects occurred under combined stress. Barley regulated independent sets of pathogen- and drought stress-responsive genes (Fig. 3a), which added up during the double stress response (Fig. 3a). The additivity for up-regulated genes under combined stress appears clearly fulfilled, but less so for down-regulated genes (Fig. 3b). The similarity scores reveal that the single stress responses show only a small overlap with each other and are thus clearly separated (Fig. 3b). There is a large overlap between WFc and DFc, especially in up-regulated DEGs. This similarity decreased at 7 dpi, likely due to the increasing impact of drought stress, which causes partially opposite regulation to the infection effect (see below). The response to infection mainly caused an up-regulation of DEGs, and was partially maintained under DFc, with a clear addition and integration of the drought response. By contrast, for both up- and down-regulated genes, the overlap of DEGs regulated in DM and DFc increased between 4 and 7 dpi. Here, too, the drought stress response increasingly dominates. This manifests also for down-regulated genes in this comparison, because barley regulates many more genes down under drought than infection (Fig. 3a-b). This suggests that under drought stress, which affects the entire plant, many biological functions are down regulated, whereas local infection response mainly causes an up-regulation of DEGs. Drought stress, for example, strongly affected energy metabolism (yellow module; Figure S10) and enriched for hydrolase and phosphatase enzyme activities involved in stress signalling and adaptation (turquoise module; Figure S7) (Schweighofer et al. 2004; Fu et al. 2024), while infection was strongly associated with transferase activities and secondary metabolism (blue module; Figure S8), representing a core metabolic activity against *Fusarium* stress (Bethke et al. 2023; Gardiner et al. 2010; Hein et al. 2025).

We further looked into gene expression patterns that illustrate the individual stress responses and dynamics of regulation (Fig. 4a, b). The subtraction of regulatory responses at the single-gene level allowed us to split the population of DEGs in those DEGs with similar expression between DFc and WFc or between DFc and DM (Fig. 4a, b). Such genes, similarly regulated across all genotypes, show clearly separable enriched biological functions (GOs) important for pathogen defence or drought stress mitigation. Interestingly, these genotype-spanning “core DEG sets” still consist of a few hundred genes (Fig. 4b, c, d). Here, the DEGs involved in pathogen defence again show strong enrichment in transferase activities, including glutathione-S-transferases and UDP-glycosyltransferases (Fig. 4c). This supports that oxidative stress balance and DON detoxification are important factors in the *Fusarium* response and are maintained even under combination stress. Notably, a recent study found in wild barley accession several UDP-glycosyltransferases as crucial in drought tolerance by modulating hormone homeostasis and secondary metabolism (Feng et al. 2024). It needs to be analysed which individual UDP-glycosyltransferases might fulfil a double function in pathogen and drought stress resistance. “Core DEGs” show an overall up-regulation of many pathogen-responsive genes during infection, often down-regulation under drought, but again up-regulation under combined stress. Among the 6,558 significantly regulated DEGs, many pathogen-induced DEGs show similar fold changes in WFc and DFc (compare cluster 2 in Fig. 5; Dataset D2). Conversely, we found cluster number 8 of 991 infection stress-related DEGs showing mostly attenuated or opposite regulation under combined stress compared to infection stress alone (Fig. 5; Dataset S-8). This includes again UDP-glycosyltransferases, glutathione S-transferases, tryptophan decarboxylases, pathogenesis-related proteins, and enriched functions in cell-wall biogenesis, together likely important for mycotoxin detoxification and basal defense against FHB (Kang and Buchenauer 2002; Boddu et al. 2006; Gardiner et al. 2010; Tucker et al. 2021; Bethke et al. 2023). This suggests an additive but contrasting regulation of DEGs in this cluster, which could reflect a trade-off between *Fusarium* and drought responses. It further shows that drought can compromise infection-induced defence outputs by weakening expression of infection-responsive genes, thus supporting fungal success.

The attenuated expression of larger groups of defence genes implies a physiological network sensitive to additional drought stress. This only partially explains our findings, but it is likely that under double stress, defence was limited or superimposed. It is also possible that transcription of genes relevant for individual stress responses was functional, but simultaneous drought may have partially compromised the translation of defence genes into proteins. This could be explained by strong alterations of the translational machinery under drought (Lei et al. 2015) and likely a suppressive role of ABA on global translation efficiency, as reported for *Arabidopsis thaliana* (Zhang et al. 2025). As shown for differently drought tolerant rice genotypes, sensitive plants show reduced translation under drought (Dawane et al. 2024). We speculate that in combination of drought with the fungal protein translation inhibitor DON, the moderately resistant varieties were restricted in their *Fusarium* response, whereas susceptibility did not further increase in the already very susceptible genotypes. This suggests studying the impact of drought on translation level of defence genes decisive for FHB resistance.

Drought stress can also limit *Fusarium* success in barley ears, when applied before infection, which might limit fungal growth by accelerated ripening and grain maturity (Hoheneder et al. 2023). In contrast to our previous finding, here, the simultaneous onset of the two stresses may have led to conflicting responses that require a prioritisation of stress responses and partial suppression of the pathogen response, which led to a better establishment of the fungus. For example, the quantitatively resistant variety Barke (Hoheneder et al. 2022; Hoheneder et al. 2023) showed a strong and early response to the fungus through regulation of many genes under both WFc and DFc conditions at 4 dpi (Fig. 4a), but the regulation subsequently shifted towards a more drought-responsive pattern at 7 dpi, similar to the other genotypes (Fig. 4b). Drought might have hampered maintenance of this strong and early defence response of Barke thus supporting FHB. Drought negatively affected photosynthesis and energy supply as indicated by GO enrichment e.g. in the yellow module (Figure S10) that showed negative correlations with several up-regulated drought stress-related phytohormones (Fig. 2c). Thus, despite a clear *Fusarium* response, further signal processing was potentially not successfully implemented into effective defence due to a lack of energy and resources or contradictory stress signals under additional drought. This suggests that breeding for drought tolerance might indirectly support disease resistance.

When barley is facing combined *Fusarium* and drought stress, its response is composed of single stresses and lacks a unique combination stress response (Hoheneder et al. 2023). This modularity is evident in the global DEG counts and overlaps (Fig. 3a, b) and in expression similarity plots (Fig. 4a, b). There is little overlap between drought and infection DEG sets, yet both align with combination stress, integrating drought- and infection-responses across cultivars (Fig. 3b). Although combined abiotic and biotic responses can lead to unique stress responses (Gupta et al. 2016; Zandalinas et al. 2021), we successfully applied an additive MLR model to predict gene expression under combined stress using data from single stresses and two independent experiments (Fig. 6; Fig. S23; (Hoheneder et al. 2023)). The MLR analysis confirmed that modularity of gene expression generally applies across stresses, making it possible to predict gene expression under combined stress from single stress responses. It is likely that barley also exhibits a modular response to other stress combinations, as suggested by results from model plants (Tan et al. 2023). However, this needs to be tested in future meta-analyses using barley transcriptomic data from other stress combinations. It is interesting that the timing of the applied stresses seems to have no influence on the additivity of the stress response regulation but on the damage potential of combined stresses (compare this study and (Hoheneder et al. 2023)). This suggests that combination stress responses can likely be approximated as overlays of infection- and drought-responsive components in barley but the parasitic fungus can only profit from contrasting regulatory responses as long as ripening and senescence has not progressed too much.

## Conclusions

Barley responses to simultaneous *Fusarium* infection and drought stress are largely additive and consist of responses to each single stress. Barley hence lacks a genetic program that is specific for the combination of FHB and drought stress. While conserved DEG sets highlight robust defence and drought adaptation functions, we also identified a large infection-related cluster with lower gene expression under combined stress. This attenuation of the pathogen response, together with genotype-specific hormonal shifts such as elevated ABA in Avalon and Barke, likely contributed to increased FHB severity in these cultivars. Nevertheless, the tested genotypes partially maintained quantitative FHB resistance even under drought, offering breeding potential that stacks genetically likely independent drought tolerance and FHB resistance. Predictive regression modelling showed that combined stress responses can be reconstructed from single stress responses, supporting an additive stress regulation in barley. Such modularity, together with genotype-dependent defence constraints, provides new insights and is relevant for breeding robust and stress adapted cereal crops.

## Supporting information

Suppl. Dataset_D1-2_DEG lists

Suppl. Dataset_M1-9_MLR results

Suppl. Dataset_P_Phenotypic data

Suppl. Dataset_S1-11_SOTA Cluster

Suppl. Dataset_E1-4_Shared DEGs_Expression Similarity

Suppl. Dataset_W1-20_WGCNA modules and eigengene data

## Acknowledgements

We are grateful to R. Dittebrandt, V. Klingl and S. Hein for technical assistance in the laboratory and in the greenhouse, S. Zuber, K. Steinmetz and R. Heinisch at the Greenhouse and Phytochamber Unit of the Plant Technology Center at the Technical University of Munich (TUM) for maintaining optimal plant growth conditions in the greenhouse. We thank Dr. C. Wurmser (Chair of Animal Physiology and Immunology, TUM) for RNA library preparations. We further thank L. Barl (Chair of Plant Breeding, TUM) for providing a porometer.

## Authors’ contributions

Conceptualisation and experimental design: FH, RH; Funding acquisition and resources: RH, CD; Laboratory and greenhouse experiments: FH, CS; Data acquisition: FH, CS, MG, CD; Data analysis, curation and visualization: FH, CS, RH; Writing the original draft, editing and finalising of the manuscript: FH, RH; All authors have read and approved the manuscript.

## Supporting Information

Additional supporting information can be found online in the Supporting Information section.

