## Supplementary tables and figures for "Transcriptome and hormone regulations shape drought stress-dependent Fusarium Head Blight susceptibility in different barley genotypes"

**Tables:**

Table S1: Statistical results for fungal DNA amounts in sampled spikes under irrigation and drought stress shown in Figure 1. Differences between treatment groups were tested using Mann-Whitney U-test at significance level of 0.05. ns = not significant, *p < 0.05.

| genotype | dpi | n | p value | significance |
|---|---|---|---|---|
| Avalon | 2 | 4 | 0.3429 | ns |
| Avalon | 4 | 4 | 0.6857 | ns |
| Avalon | 7 | 4 | 0.0286 | * |
| Barke | 2 | 4 | > 0.9999 | ns |
| Barke | 4 | 4 | 0.2000 | ns |
| Barke | 7 | 4 | 0.0286 | * |
| Morex | 2 | 4 | 0.3429 | ns |
| Morex | 4 | 4 | 0.6286 | ns |
| Morex | 7 | 4 | > 0.9999 | ns |
| Palmella Blue | 2 | 4 | 0.3429 | ns |
| Palmella Blue | 4 | 4 | 0.8857 | ns |
| Palmella Blue | 7 | 4 | 0.8857 | ns |

Table S2: Statistical results for FHB symptoms assessed on spikes under irrigation and drought stress shown in Figure 1. Differences between treatment groups were tested using Mann-Whitney U-test at significance level of 0.05. ns = not significant, *p < 0.05, **p < 0.01, ***p < 0.001.

| genotype | dpi | n | p value | significance |
|---|---|---|---|---|
| Avalon | 2 | 12 | 0.9435 | ns |
| Avalon | 4 | 12 | 0.2977 | ns |
| Avalon | 7 | 12 | 0.4343 | ns |
| Barke | 2 | 12 | 0.0318 | * |
| Barke | 4 | 12 | 0.0004 | *** |
| Barke | 7 | 12 | 0.0019 | ** |
| Morex | 2 | 12 | 0.504 | ns |
| Morex | 4 | 12 | 0.0378 | * |
| Morex | 7 | 12 | 0.1734 | ns |
| Palmella Blue | 2 | 20 | 0.9626 | ns |
| Palmella Blue | 4 | 20 | 0.2339 | ns |
| Palmella Blue | 7 | 20 | 0.0797 | ns |

Table S3: Statistical results for SPAD values measured on flag [F] and flag-1 [F-1] leaves under irrigation and drought stress as shown in Figure 1. Differences between treatment groups were tested using Mann-Whitney U-test at significance level of 0.05. ns = not significant, *p < 0.05, **p < 0.01, ***p < 0.001, ****p < 0.0001.

| **genotype** | **dpi** | **leaf stage** | **n** | **p value** | **significance** |
|---|---|---|---|---|---|
| Avalon | 2 | F | 18 | 0.4762 | ns |
| Avalon | 4 | F | 18 | 0.0126 | * |
| Avalon | 7 | F | 18 | < 0.0001 | **** |
| Avalon | 2 | F-1 | 18 | 0.4017 | ns |
| Avalon | 4 | F-1 | 18 | < 0.0001 | **** |
| Avalon | 7 | F-1 | 18 | < 0.0001 | **** |
| Barke | 2 | F | 18 | 0.3268 | ns |
| Barke | 4 | F | 18 | 0.5059 | ns |
| Barke | 7 | F | 18 | < 0.0001 | **** |
| Barke | 2 | F-1 | 18 | 0.6004 | ns |
| Barke | 4 | F-1 | 18 | 0.004 | ** |
| Barke | 7 | F-1 | 18 | < 0.0001 | **** |
| Morex | 2 | F | 18 | 0.0410 | * |
| Morex | 4 | F | 18 | 0.0056 | ** |
| Morex | 7 | F | 18 | < 0.0001 | **** |
| Morex | 2 | F-1 | 18 | 0.2056 | ns |
| Morex | 4 | F-1 | 18 | 0.0249 | * |
| Morex | 7 | F-1 | 18 | < 0.0001 | **** |
| Palmella Blue | 2 | F | 18 | 0.3425 | ns |
| Palmella Blue | 4 | F | 18 | 0.1782 | ns |
| Palmella Blue | 7 | F | 18 | < 0.0001 | **** |
| Palmella Blue | 2 | F-1 | 18 | 0.4666 | ns |
| Palmella Blue | 4 | F-1 | 18 | 0.0559 | ns |
| Palmella Blue | 7 | F-1 | 18 | < 0.0001 | **** |

**Figures:**

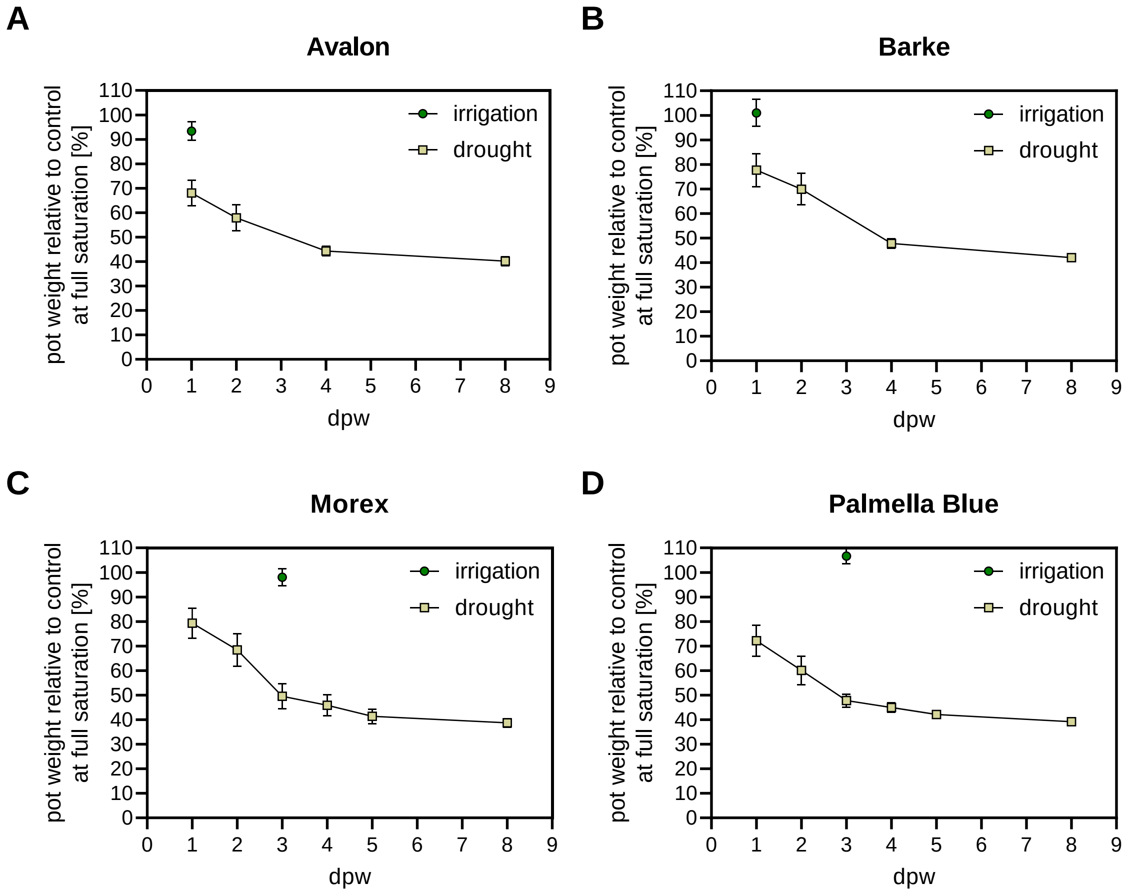


Figure S1: Mean pot weights relative to the control pots in (A) Avalon, (B) Barke, (C) Morex and (D) Palmella Blue. Pot weights are shown in relation to mean pot weights at full saturation (0 days post watering). The line plots show the means for 18 individual pots under irrigation or drought. Error bars show standard deviation.

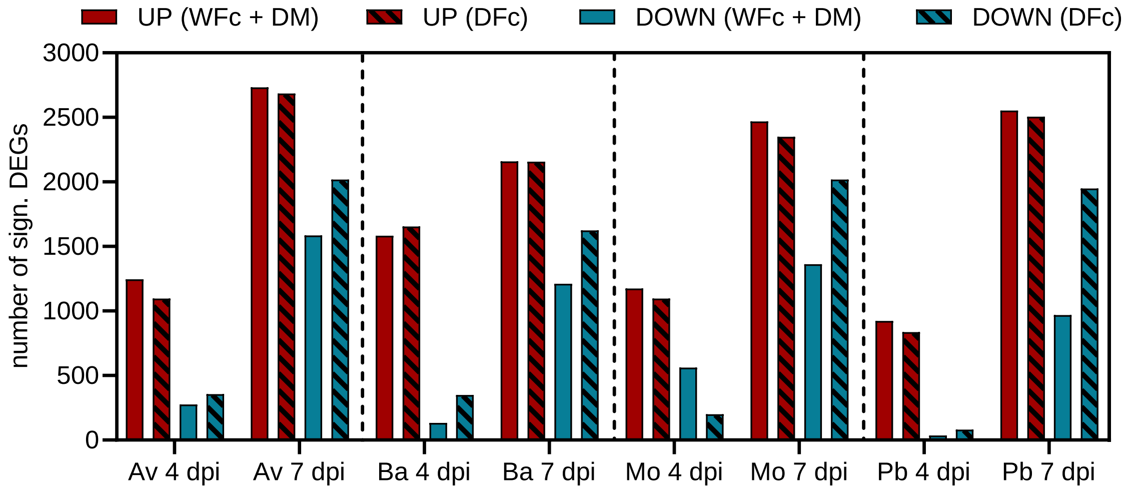


| Genotype | dpi | Treatment | DEGs [UP] | DEGs [DOWN] | Sum of DEGs [UP + DOWN] |
|---|---|---|---|---|---|
| Avalon | 4 | DFc | 1097 | 357 | 1454 |
| Avalon | 4 | WFc | 804 | 29 | 833 |
| Avalon | 4 | DM | 441 | 248 | 689 |
| Avalon | 7 | DFc | 2685 | 2019 | 4704 |
| Avalon | 7 | WFc | 910 | 45 | 955 |
| Avalon | 7 | DM | 1822 | 1541 | 3363 |
| Barke | 4 | DFc | 1655 | 349 | 2004 |
| Barke | 4 | WFc | 1169 | 37 | 1206 |
| Barke | 4 | DM | 415 | 96 | 511 |
| Barke | 7 | DFc | 2156 | 1624 | 3780 |
| Barke | 7 | WFc | 642 | 28 | 670 |
| Barke | 7 | DM | 1518 | 1183 | 2701 |
| Morex | 4 | DFc | 1096 | 200 | 1296 |
| Morex | 4 | WFc | 328 | 2 | 330 |
| Morex | 4 | DM | 846 | 559 | 1405 |
| Morex | 7 | DFc | 2349 | 2019 | 4368 |
| Morex | 7 | WFc | 749 | 22 | 771 |
| Morex | 7 | DM | 1719 | 1341 | 3060 |
| Palmella Blue | 4 | DFc | 837 | 81 | 918 |
| Palmella Blue | 4 | WFc | 638 | 5 | 643 |
| Palmella Blue | 4 | DM | 286 | 31 | 317 |
| Palmella Blue | 7 | DFc | 2505 | 1949 | 4454 |
| Palmella Blue | 7 | WFc | 1108 | 113 | 1221 |
| Palmella Blue | 7 | DM | 1444 | 855 | 2299 |

Figure S2: Sums of up- or down-regulated DEGs in single stresses and combination stress. The bars represent the sums of significantly (FDR-corrected $p < 0.01$) up- or down-regulated DEGs across the four tested genotypes under infection, drought and combined infection plus drought stress at 4 and 7 dpi. Red bars represent up-regulated DEGs, blue bars represent down-regulated DEGs. The respective data is summarized in the table below.

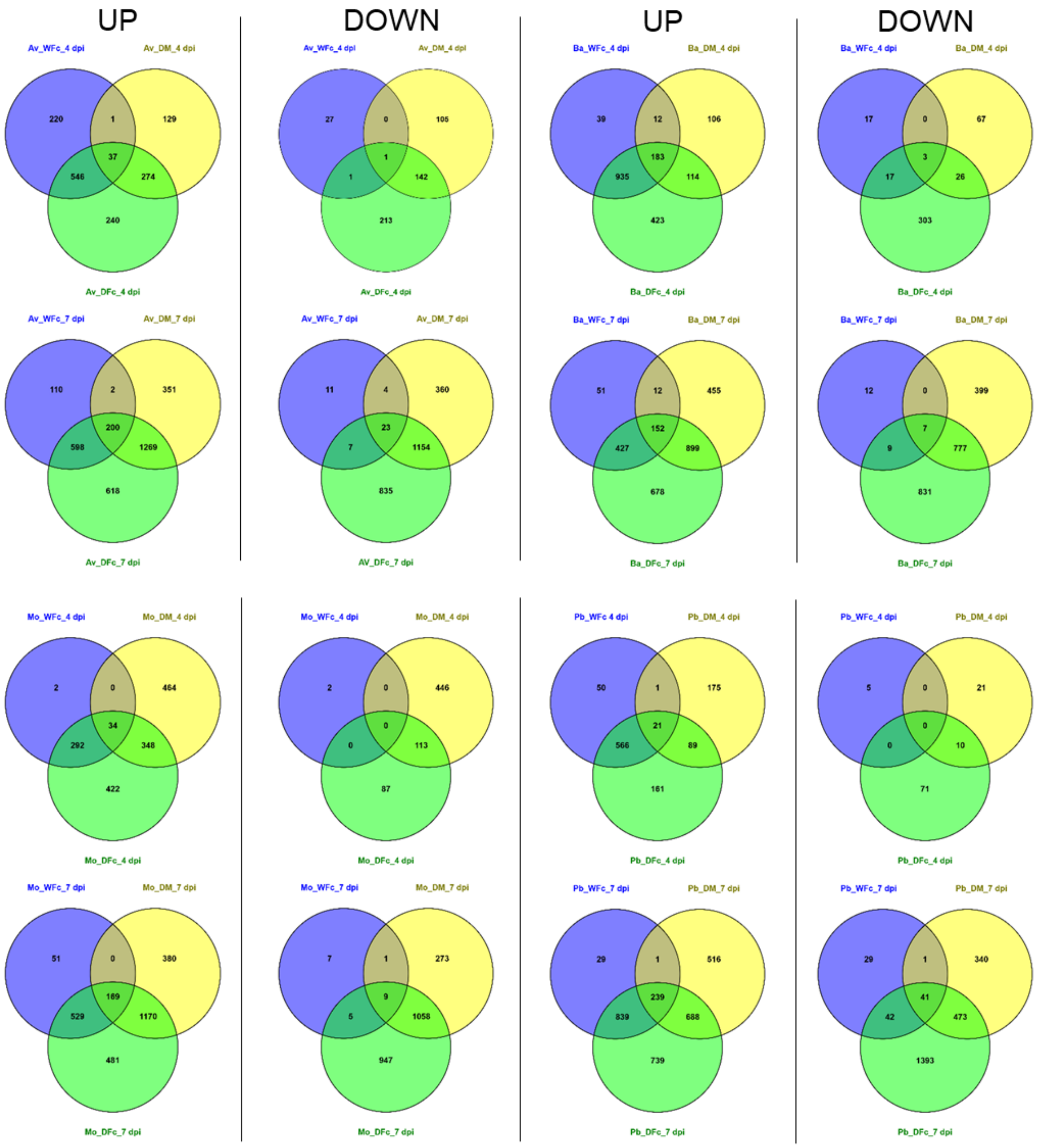


Figure S3: Venn diagrams for significantly (FDR < 0.01) up- or down-regulated differential expressed genes (DEGs) displaying overlapping DEG numbers between single stresses (WFc, DM) and combination stress (DFc) per cultivar (Av = Avalon, Ba = Barke, Mo = Morex, Pb = Palmella Blue) and time point.

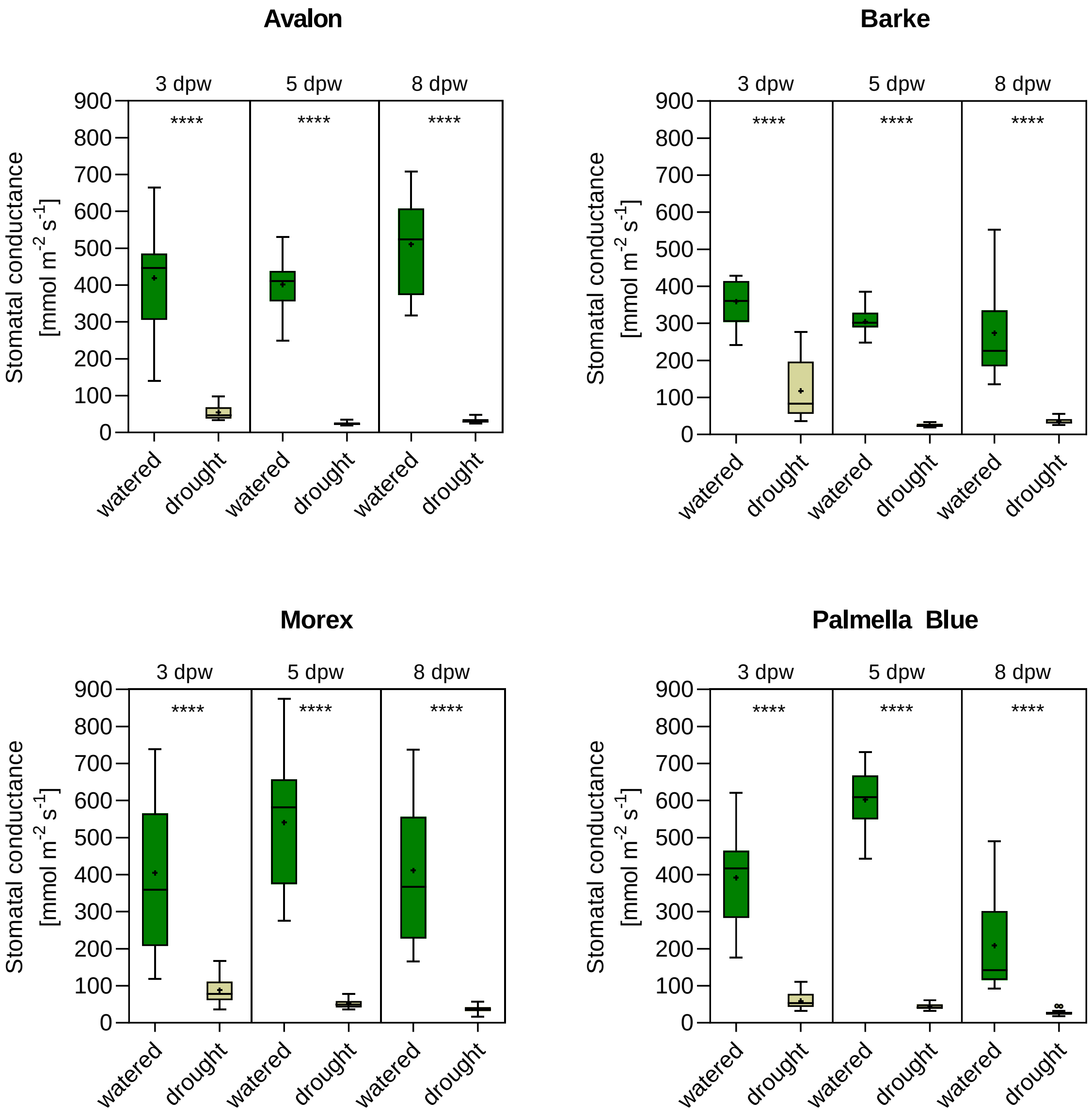


Figure S4: The graphs display the measured stomatal conductance (SC, mmol $m^{-2}$ $s^{-1}$) of upper leaf stages at 3, 5 and 8 days post-watering (dpw) in drought-stressed and watered plants for Avalon (A), Barke (B), Morex (C), and Palmella Blue (D). Data of 15 individual F and F-1 leaves was pooled for each treatment and time point. The box plots depict the median, mean (+), interquartile range and data distribution (whiskers) of SC. Statistical significance was tested using the Mann-Whitney U-test (**** $p < 0.0001$).

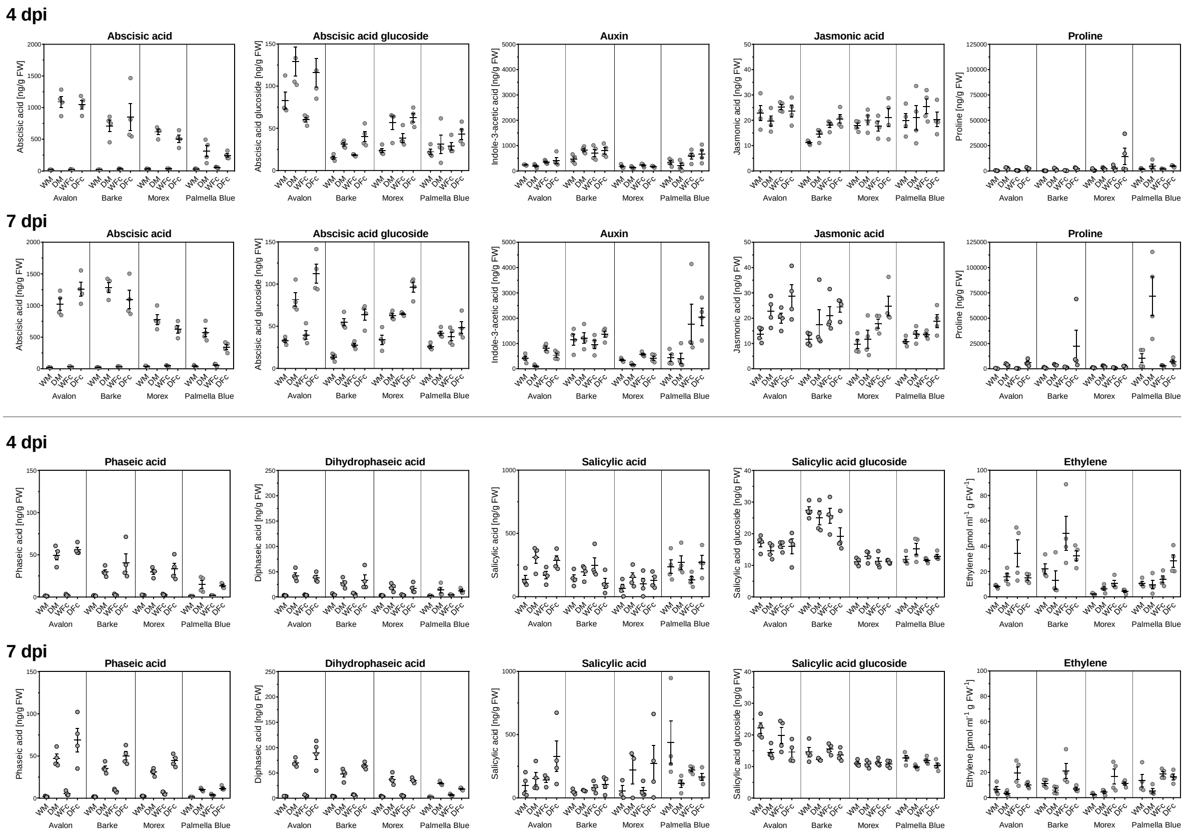


Figure S5: Quantities of phytohormones and proline measured in watered, infected, drought-stressed and drought plus infected cultivars Avalon, Barke, Morex and Palmella Blue at 4 and 7 dpi. The scatter dot plots show the individual quantities measured in four individual spike samples. The quantities of phytohormones and the osmolyte proline are presented in ng per gram fresh weight. The gaseous ethylene production was measured on additional barley spikes, which were sampled in parallel to those spikes used for fungal DNA and phytohormone quantification. Ethylene amounts are displayed in picomol per ml air and gram fresh weight. Bars indicate the mean, and error bars represent the standard deviation of the mean.

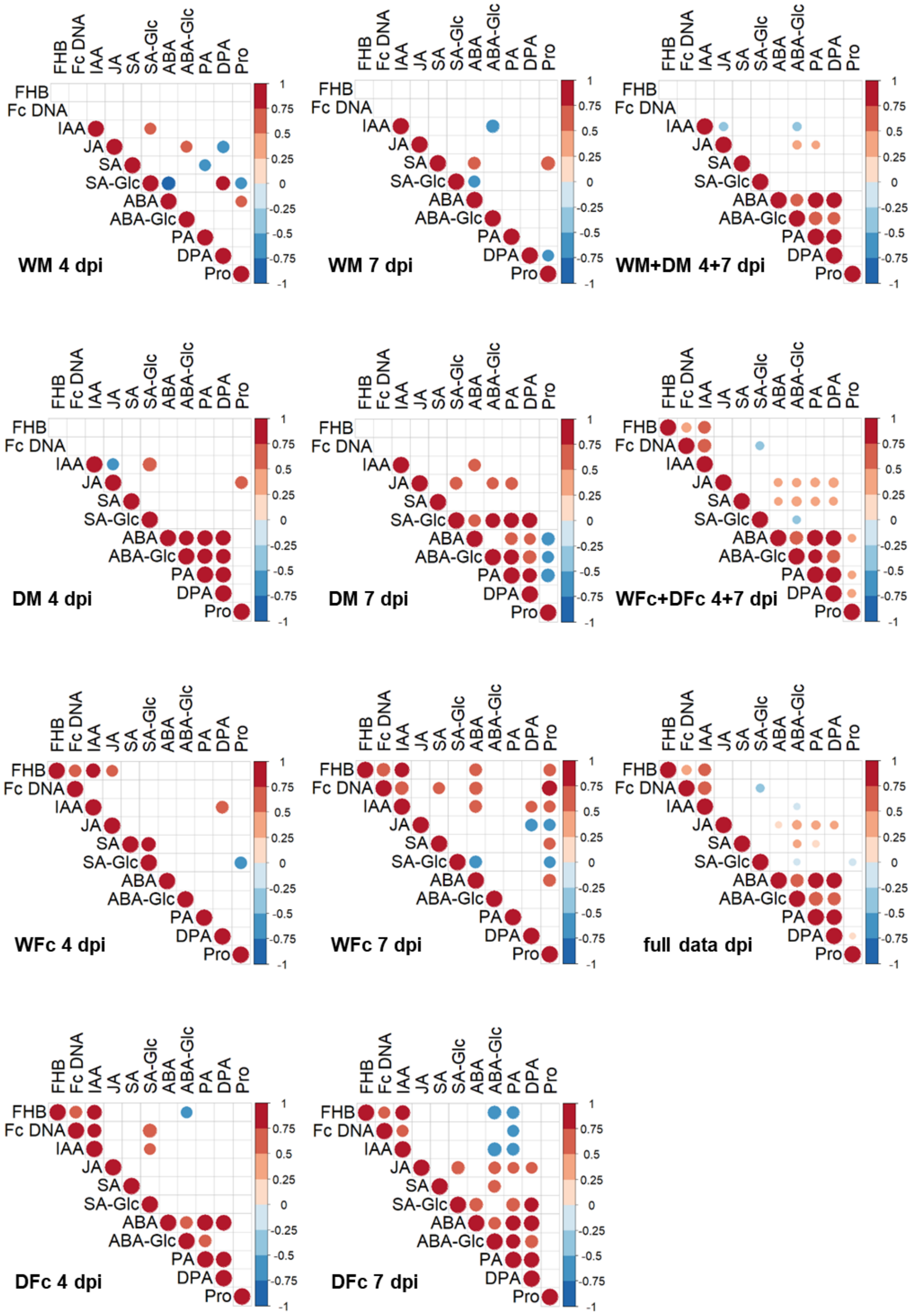


Figure S6: Pearson's correlation ($r^2$) matrices showing general relationships between mean phytohormone levels, Fusarium head blight (FHB) severity (proportion of symptomatic grains), and fungal DNA content in barley spikes at 4 and 7 dpi across the four tested barley genotypes and different treatments (WM, WFc, DM, DFc). In addition, correlation matrices are shown using the data for all mock-treated (WM + DM) or all infected (WFc + DFc) samples. Significant correlations ($p < 0.01$) are shown with coloured circles indicating the magnitude of regression coefficients.

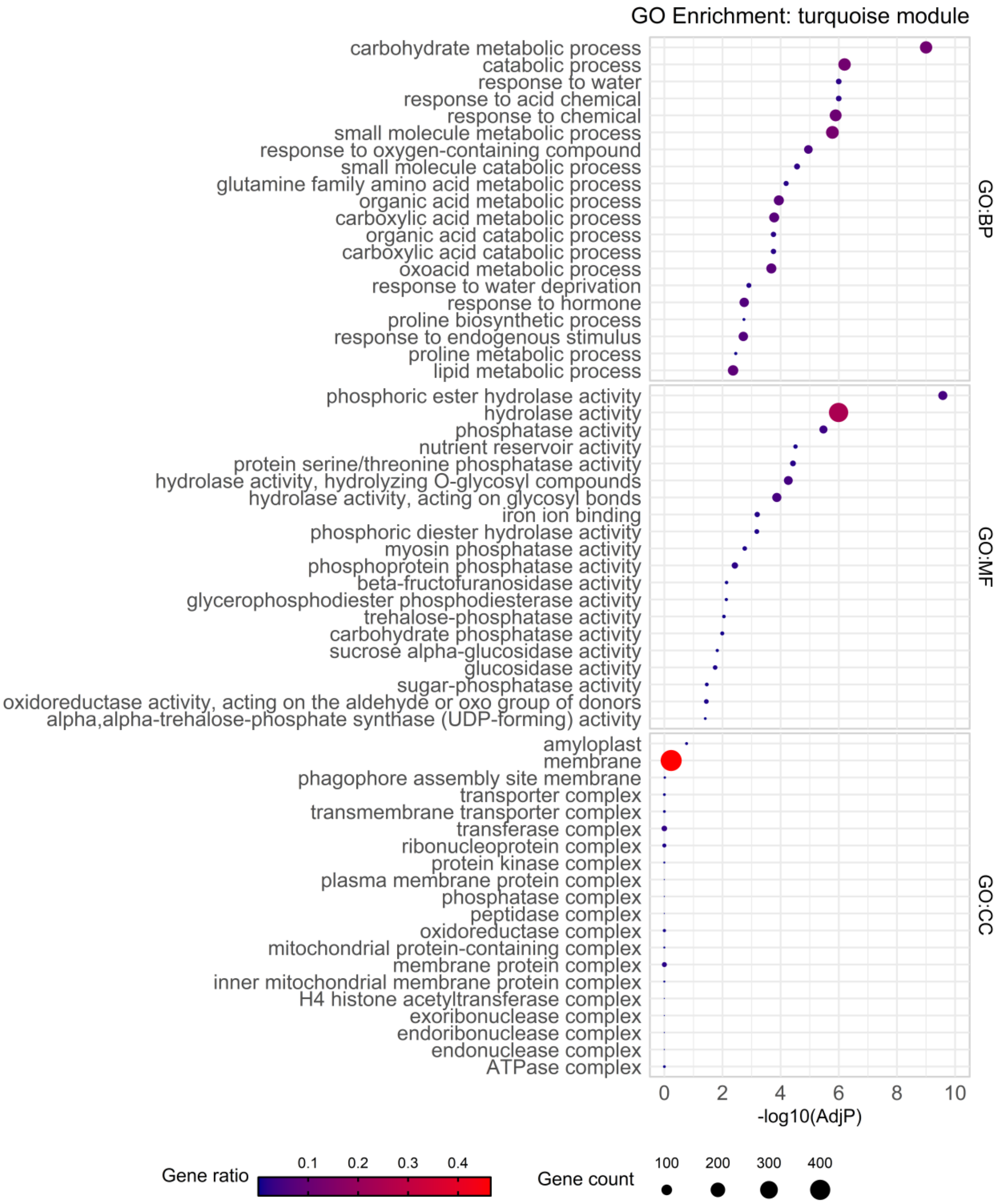


Figure S7: GO enrichment for DEGs found in WGCNA module turquoise for biological process (BP), molecular function (MF) and cellular component (CC).

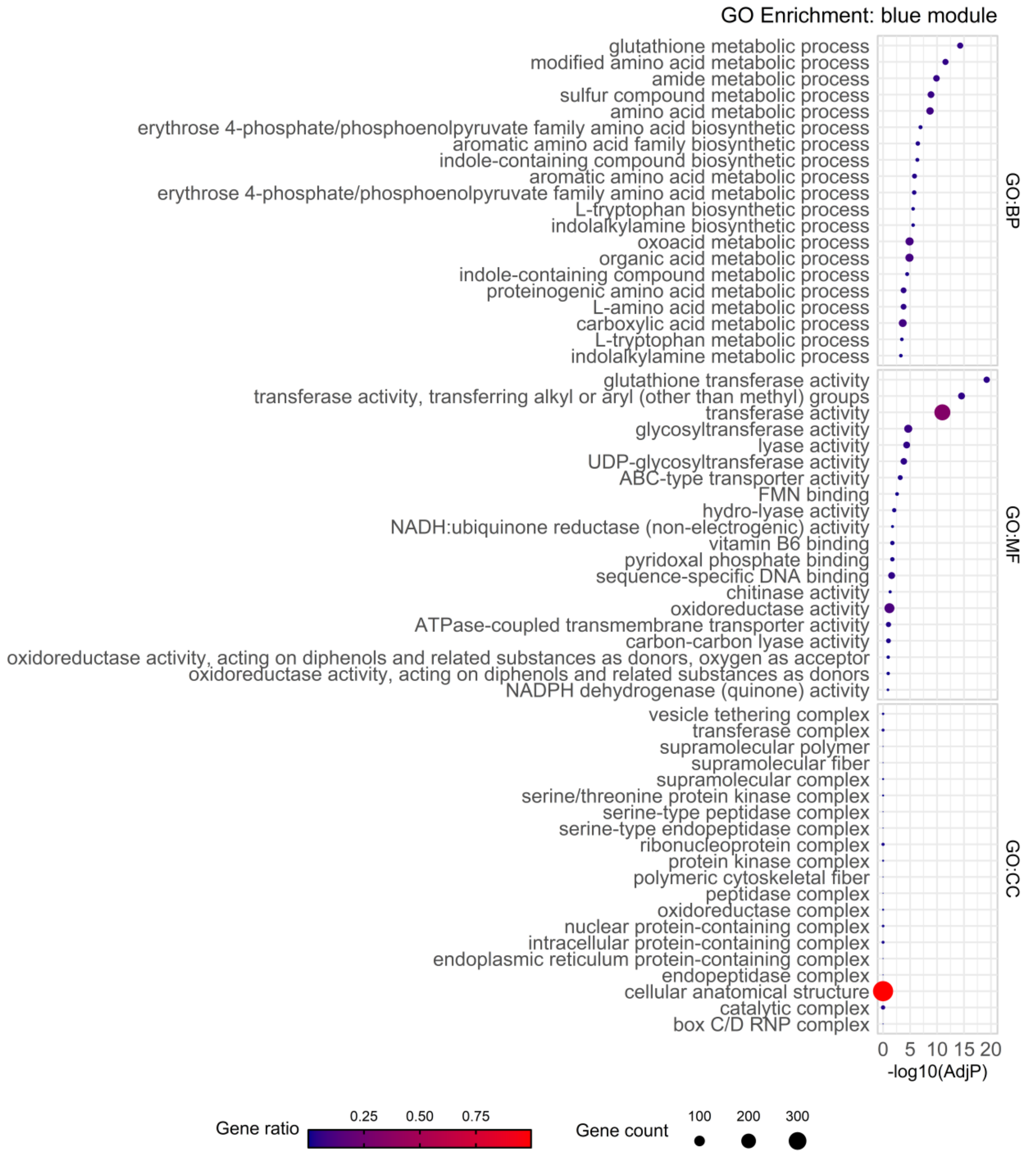


Figure S8: GO enrichment for DEGs found in WGCNA module blue for biological process (BP), molecular function (MF) and cellular component (CC).

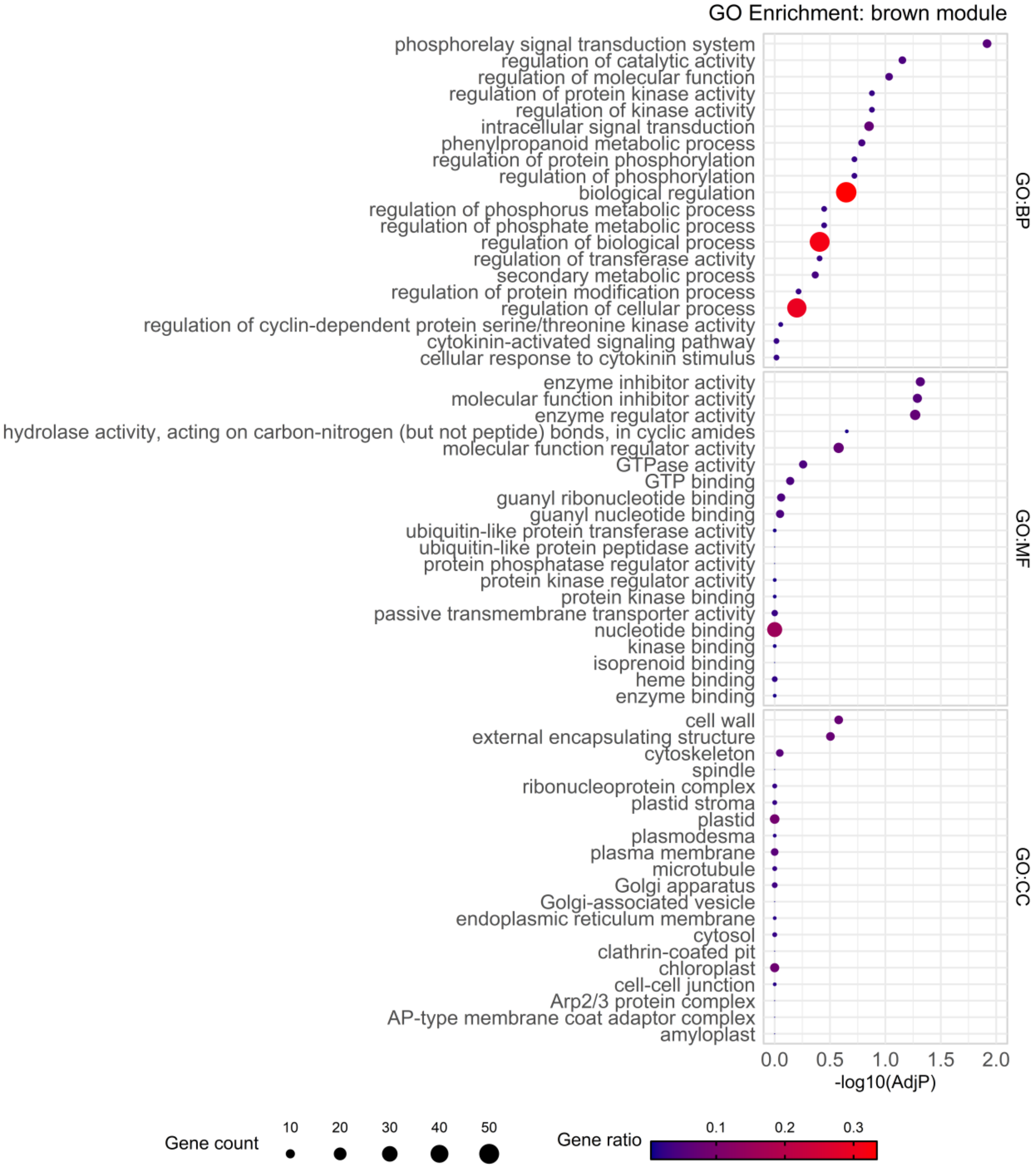


Figure S9: GO enrichment for DEGs found in WGCNA module brown for biological process (BP), molecular function (MF) and cellular component (CC).

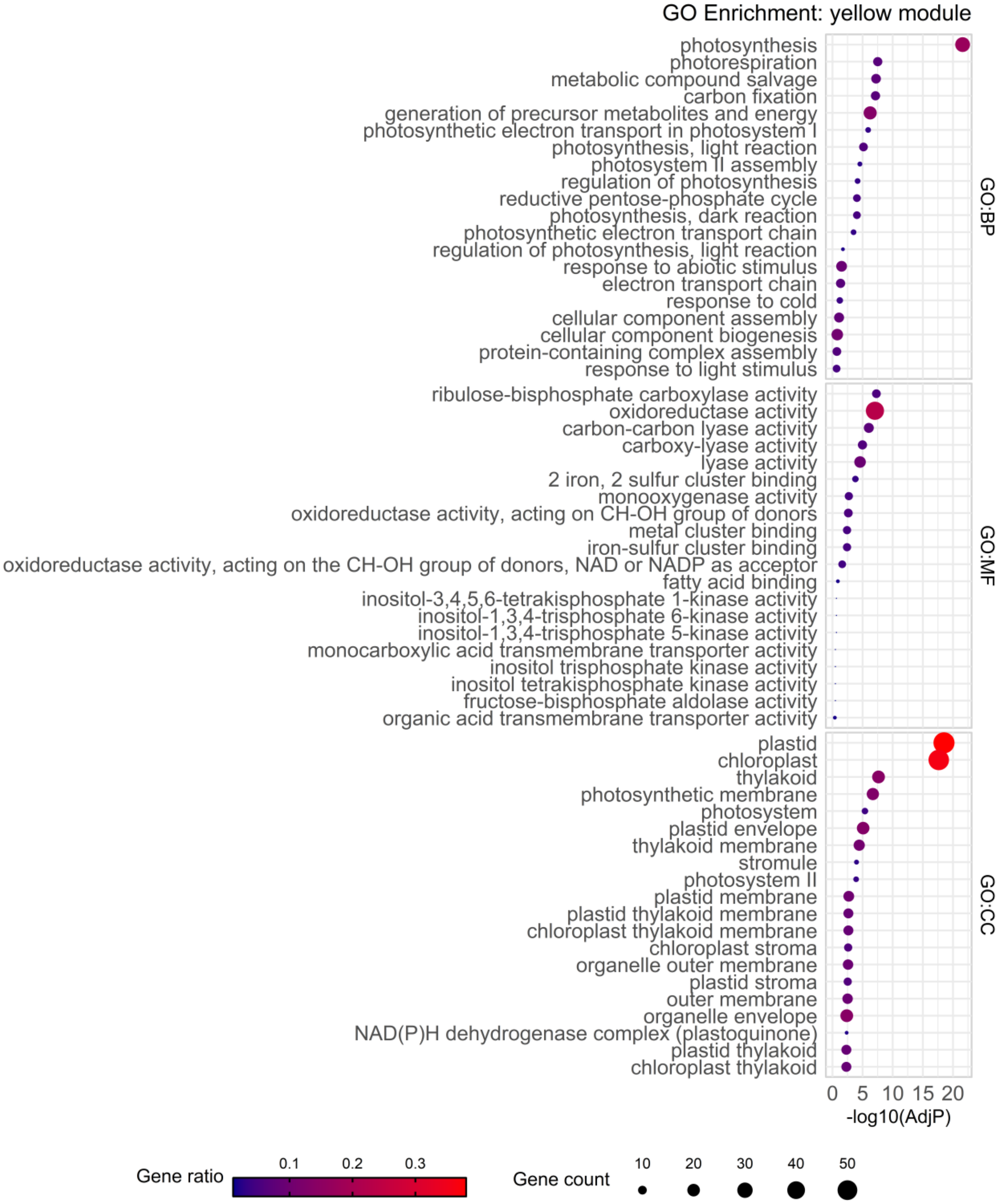


Figure S10: GO enrichment for DEGs found in WGCNA module yellow for biological process (BP), molecular function (MF) and cellular component (CC).

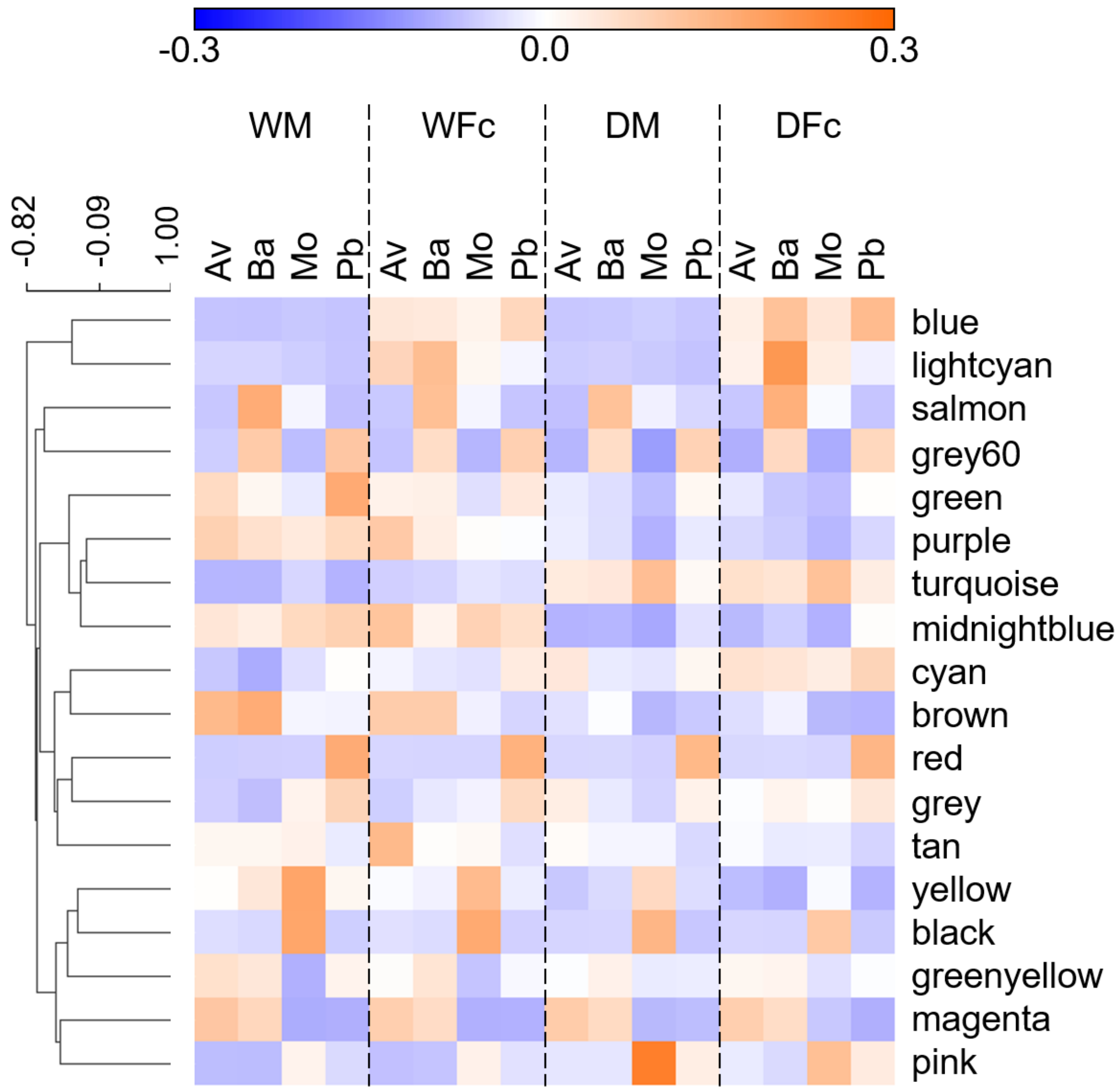


Figure S11: Clustered eigengene expression values of each WGCNA module across all genotypes and stress treatments. The heat map shows the hierarchically clustered means (4 and 7 dpi) of eigengene values for each WGCNA module.

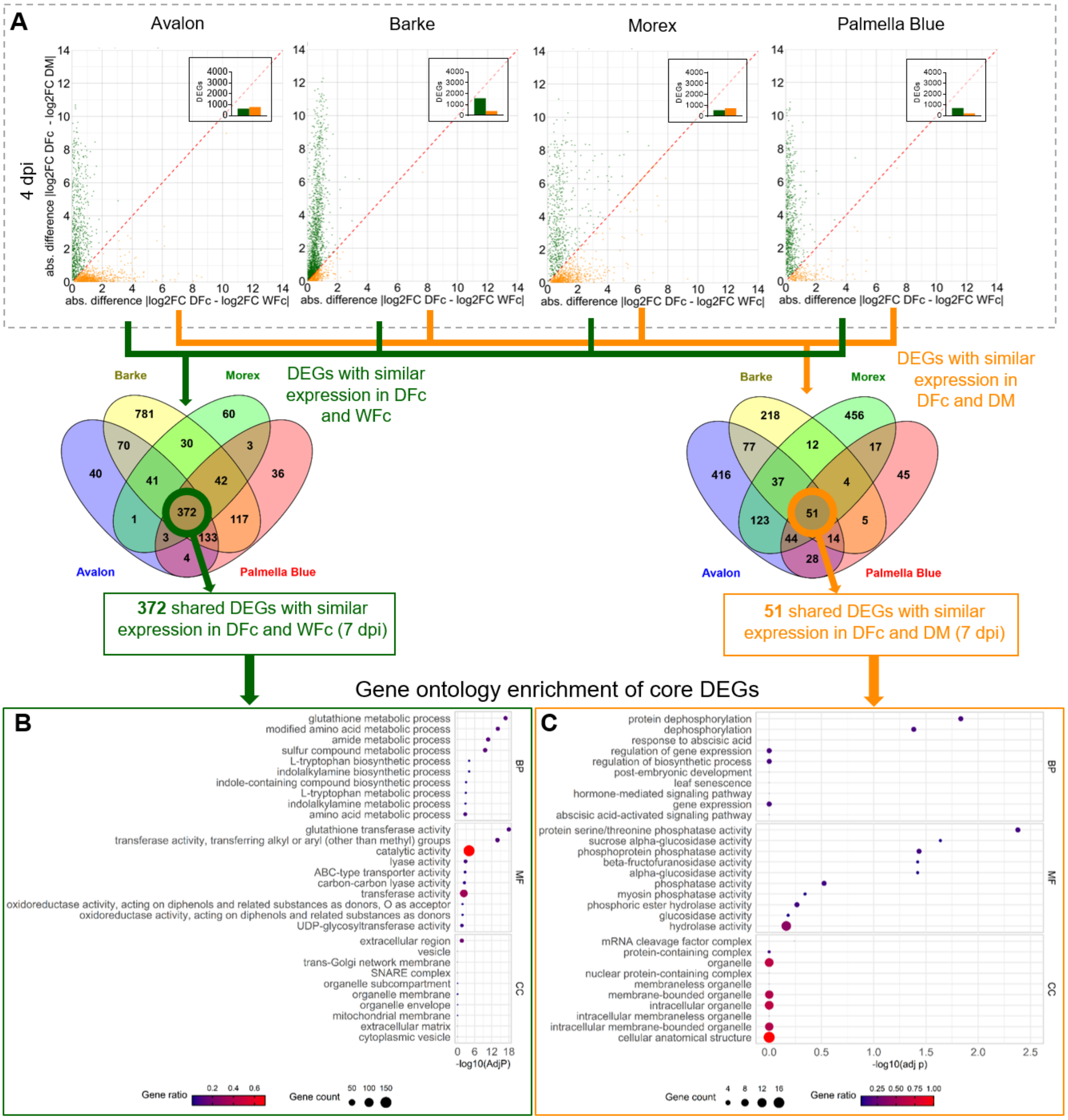


Figure S12: Expression Similarities and GO enrichment analyses of shared core DEGs at 4 dpi. (A) Expression similarity plots for significantly (FDR-corrected p < 0.01) regulated DEGs at 4 dpi. DEGs with similar expression under infection and combined stress are shown as green dots; those with similar expression under drought and combined stress are shown in orange. Bar graphs within each plot indicate total DEG counts. Venn diagrams show the number of shared DEGs with similar expression between DFc and WFc or DFc and DM across all four genotypes at 4 dpi. (B) GO enrichment for DEGs with similar expression in DFc and WFc. (C) GO enrichment for DEGs with similar expression under DFc and DM for biological process (BP), molecular function (MF) and chemical component (CC).

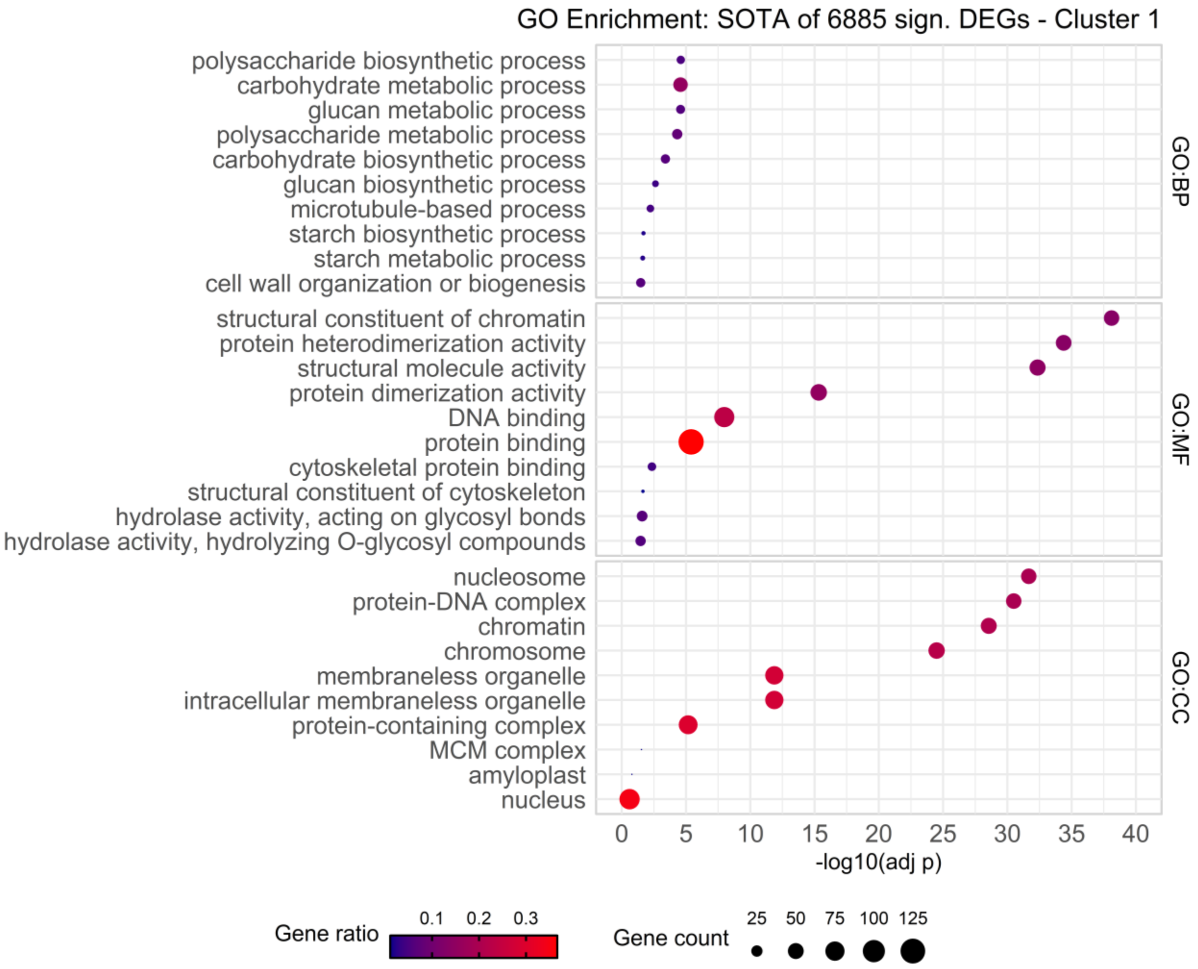


Figure S13: GO enrichment for DEGs found in SOTA cluster 1 for biological process (BP), molecular function (MF) and cellular component (CC).

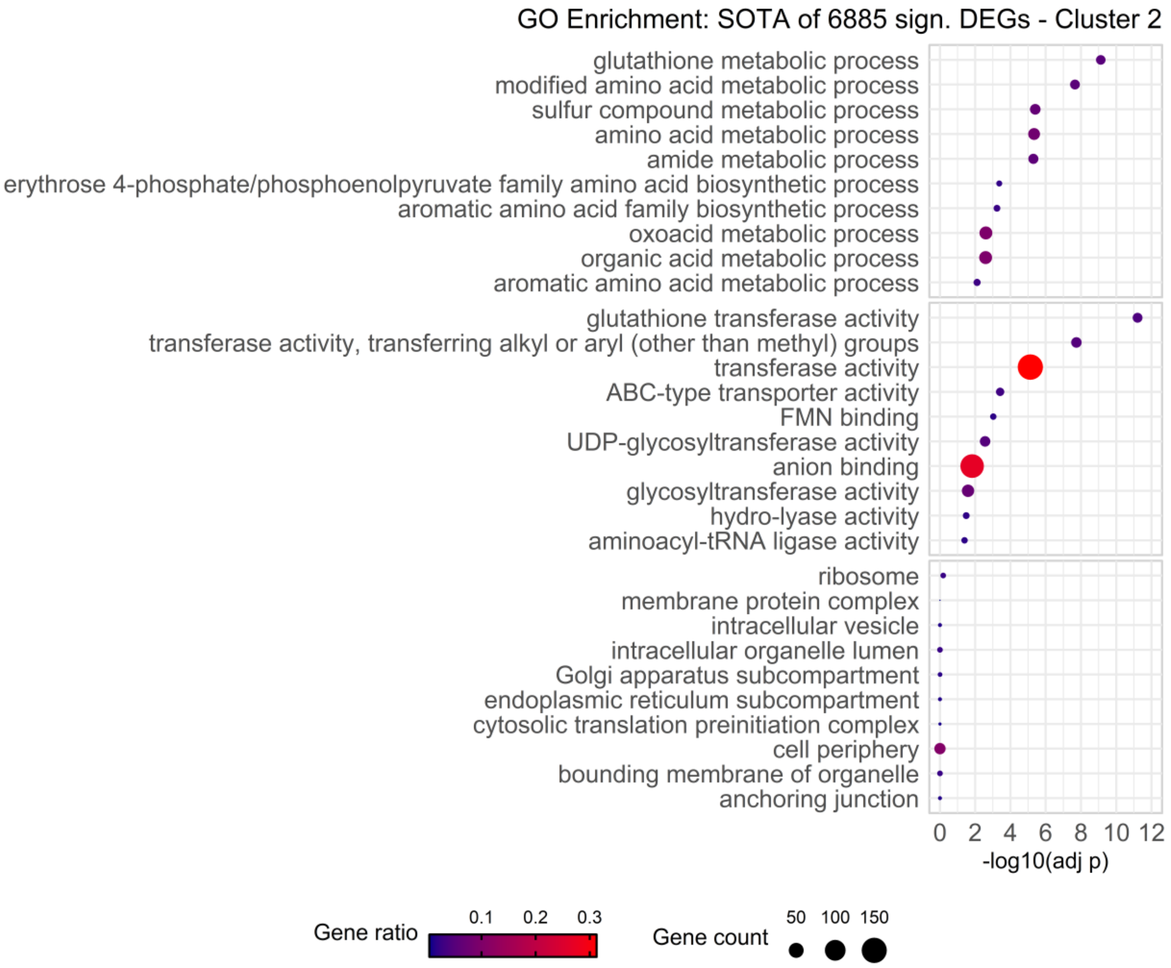


Figure S14: GO enrichment for DEGs found in SOTA cluster 2 for biological process (BP), molecular function (MF) and cellular component (CC).

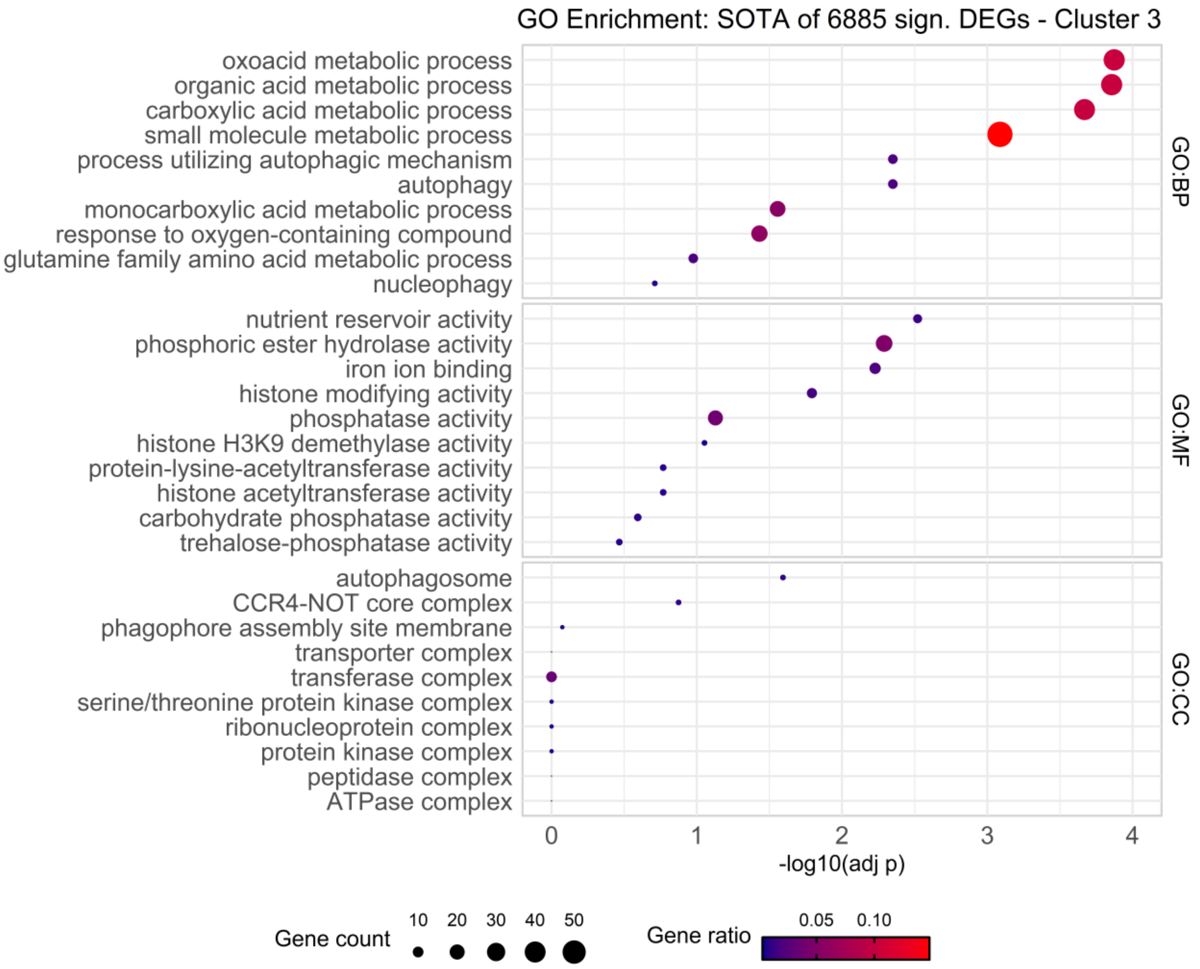


Figure S15: GO enrichment for DEGs found in SOTA cluster 3 for biological process (BP), molecular function (MF) and cellular component (CC).

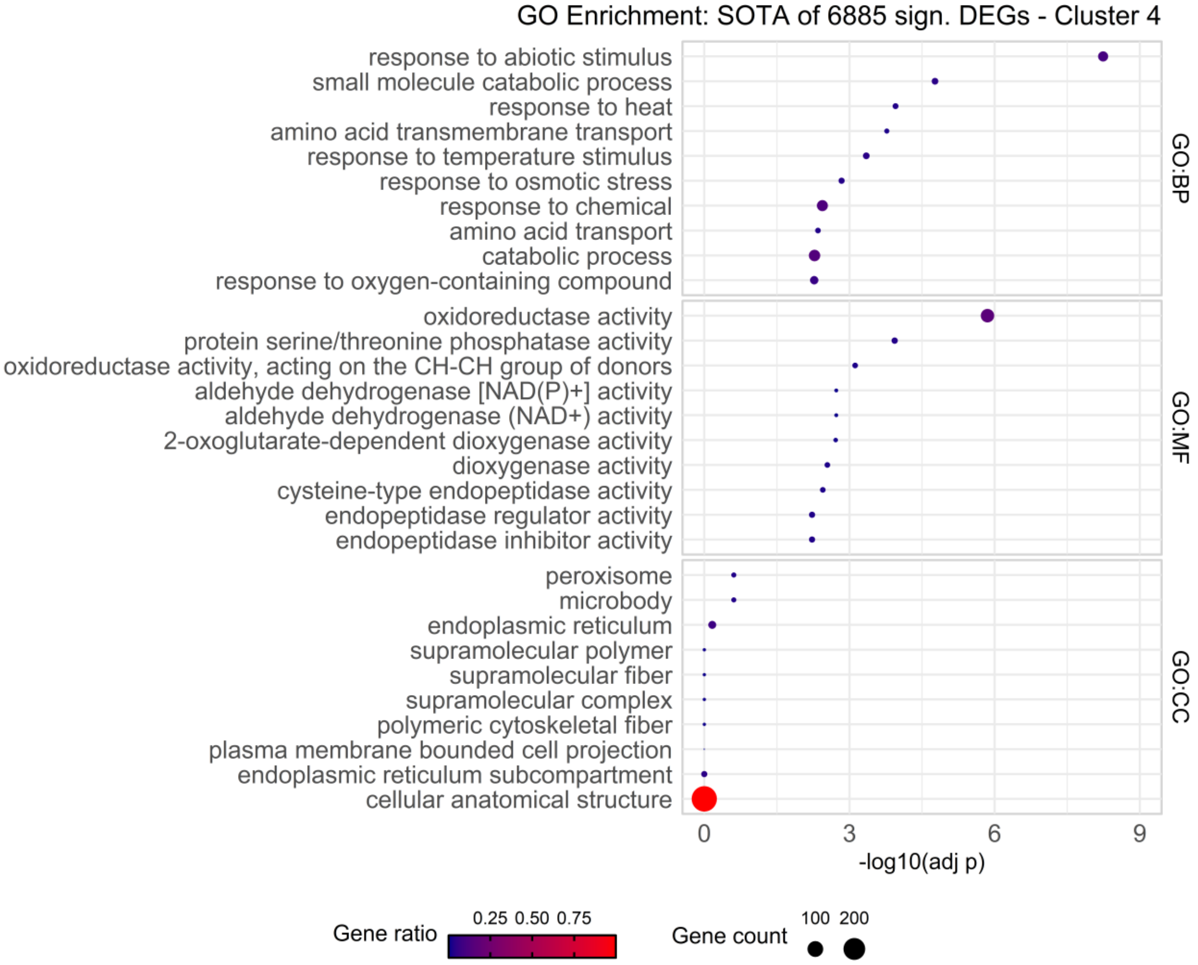


Figure S16: GO enrichment for DEGs found in SOTA cluster 4 for biological process (BP), molecular function (MF) and cellular component (CC).

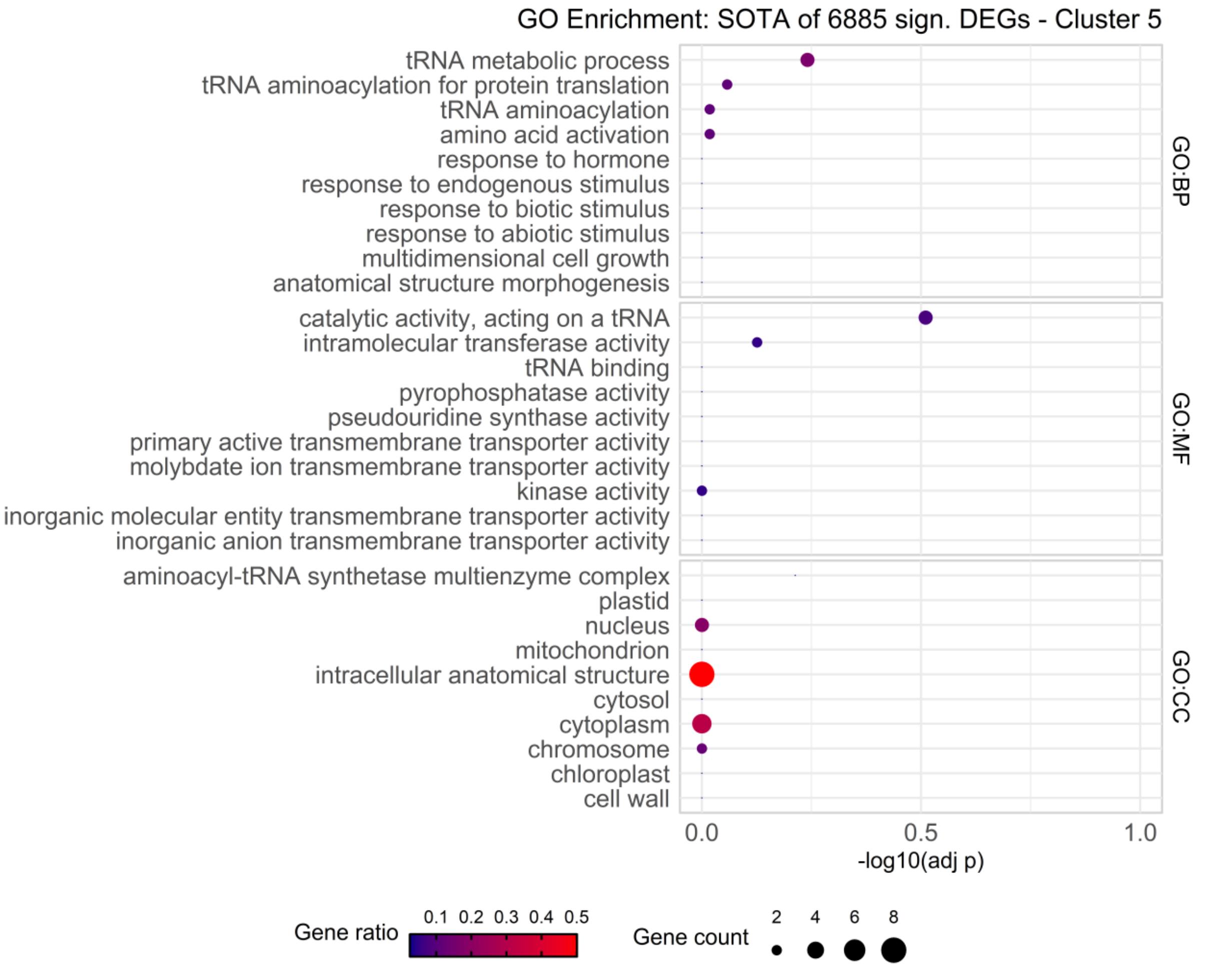


Figure S17: GO enrichment for DEGs found in SOTA cluster 5 for biological process (BP), molecular function (MF) and cellular component (CC).

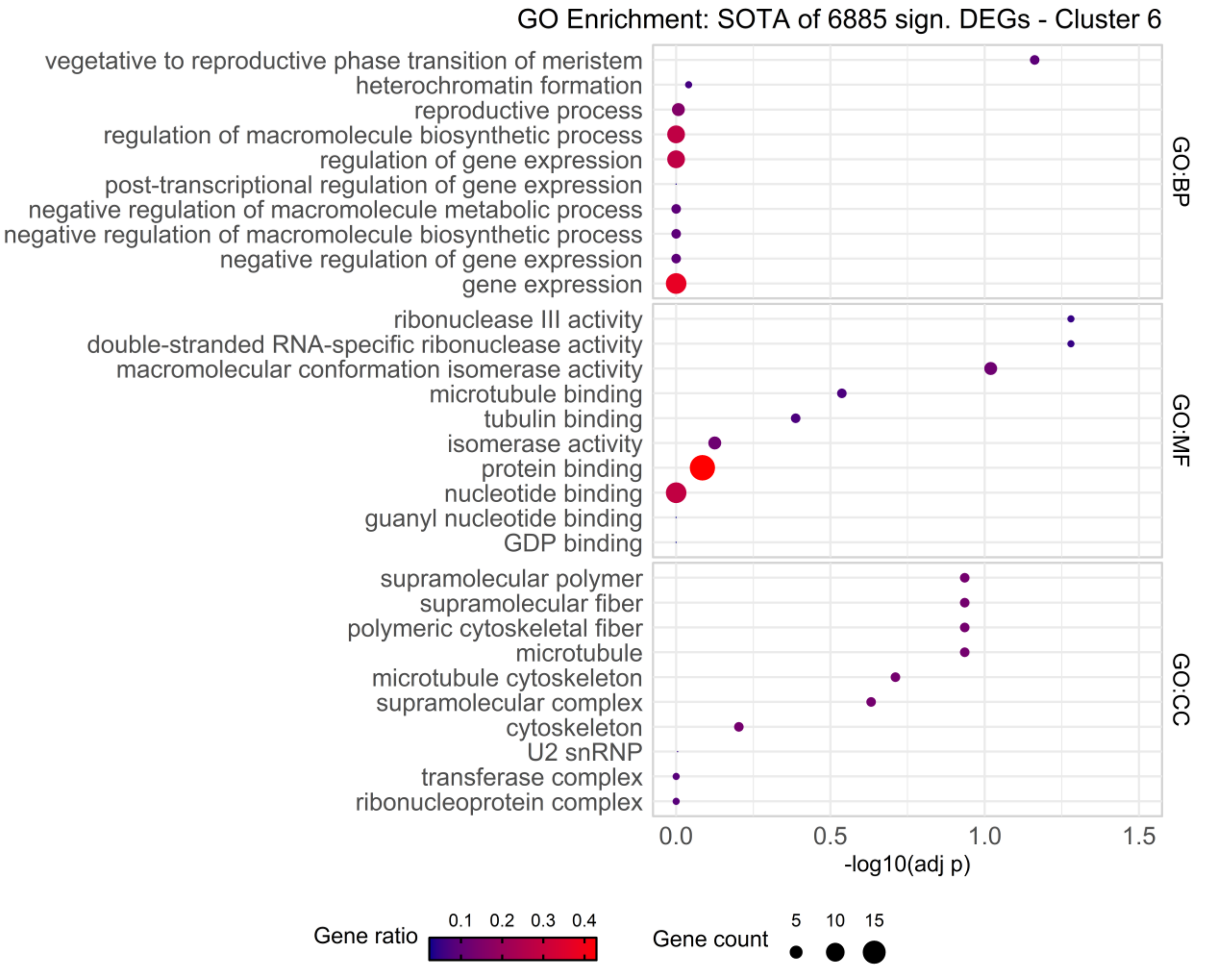


Figure S18: GO enrichment for DEGs found in SOTA cluster 6 for biological process (BP), molecular function (MF) and cellular component (CC).

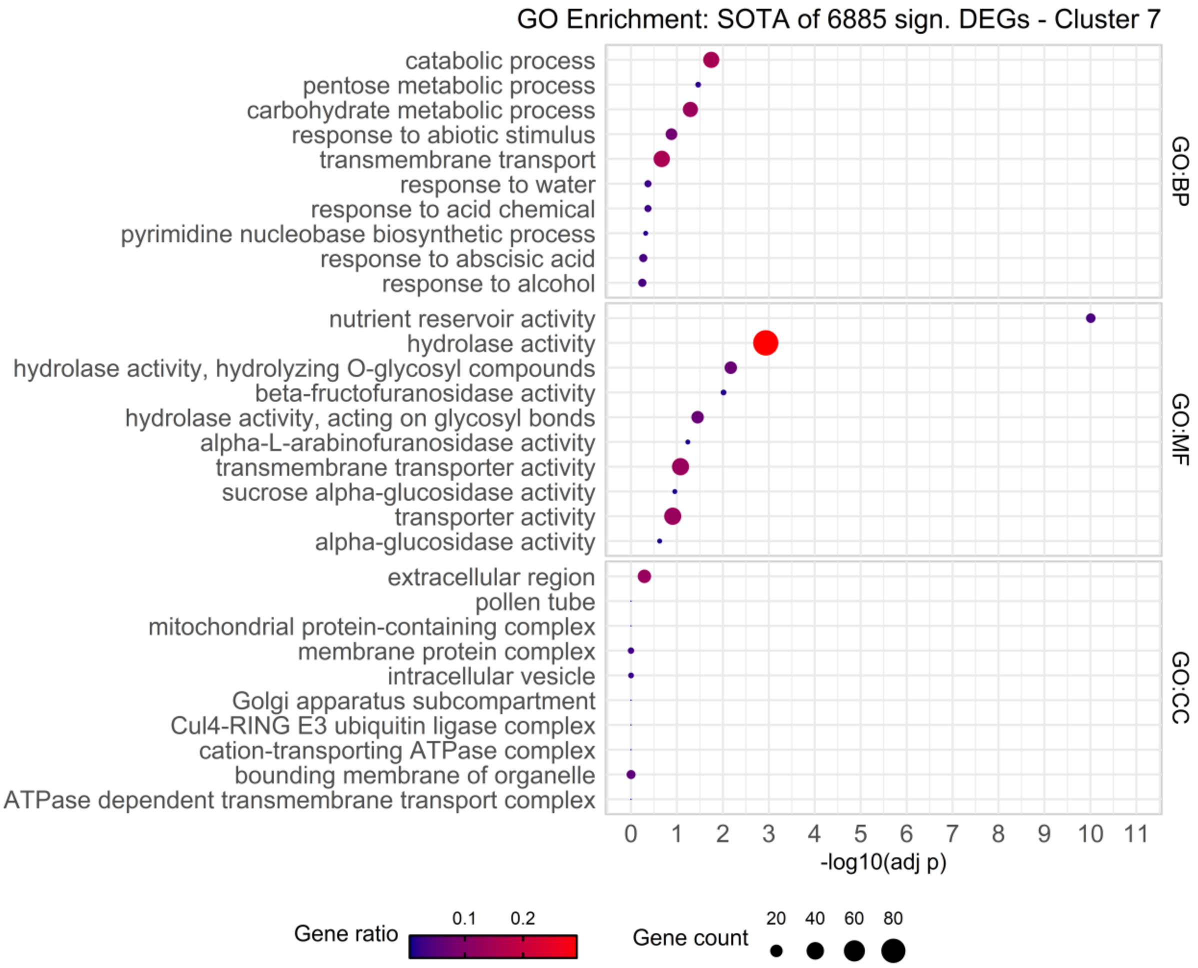


Figure S19: GO enrichment for DEGs found in SOTA cluster 7 for biological process (BP), molecular function (MF) and cellular component (CC).

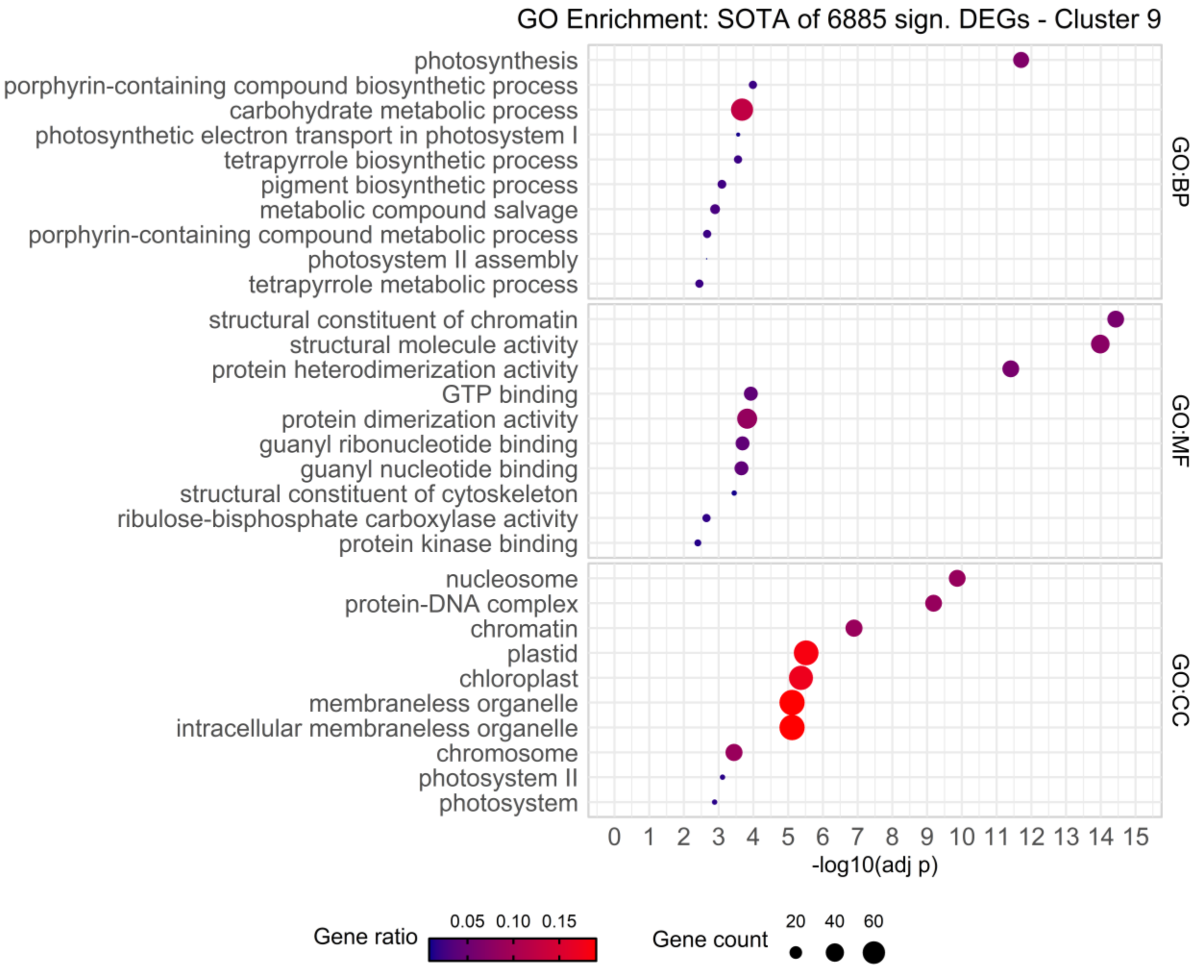


Figure S20: GO enrichment for DEGs found in SOTA cluster 9 for biological process (BP), molecular function (MF) and cellular component (CC).

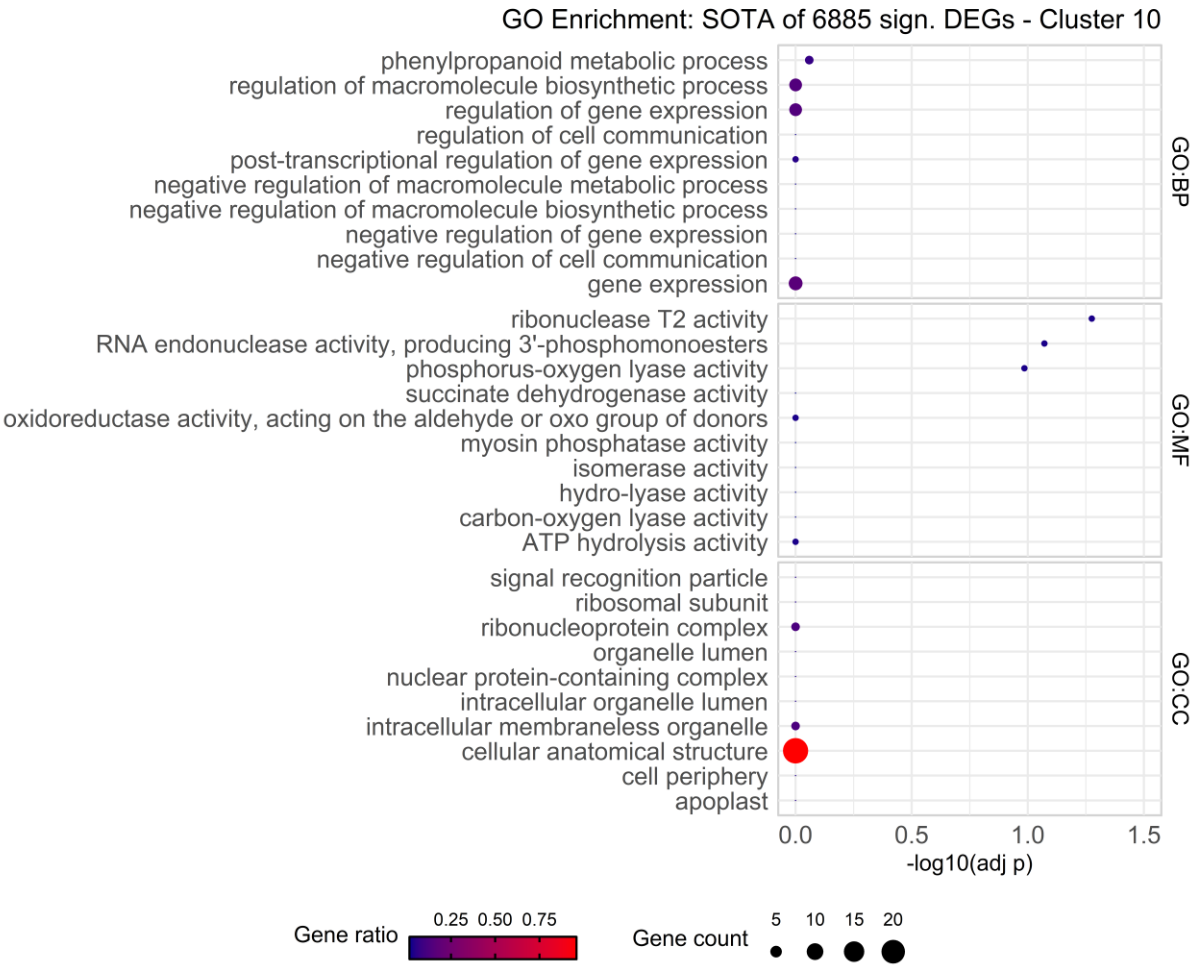


Figure S21: GO enrichment for DEGs found in SOTA cluster 10 for biological process (BP), molecular function (MF) and cellular component (CC).

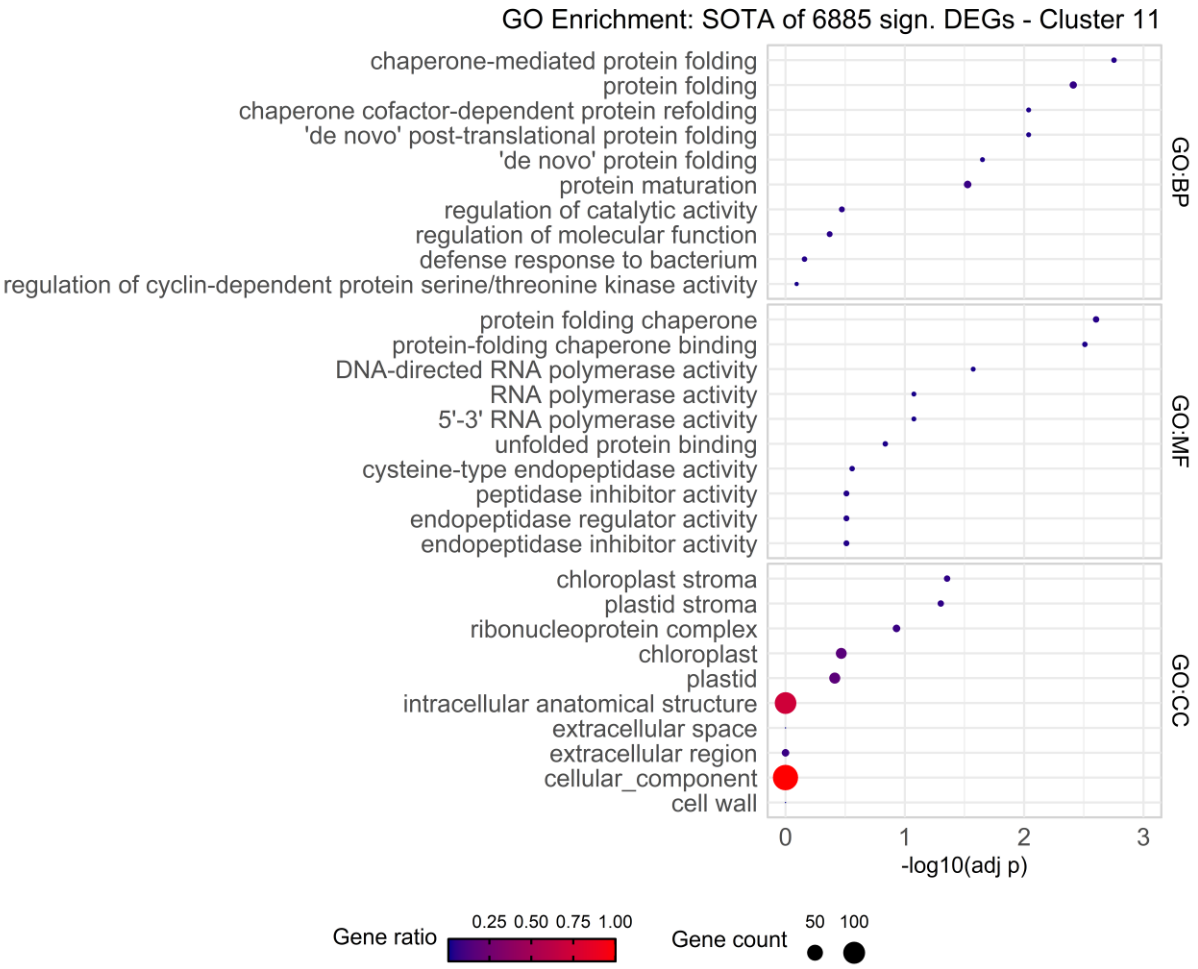


Figure S22: GO enrichment for DEGs found in SOTA cluster 11 for biological process (BP), molecular function (MF) and cellular component (CC).

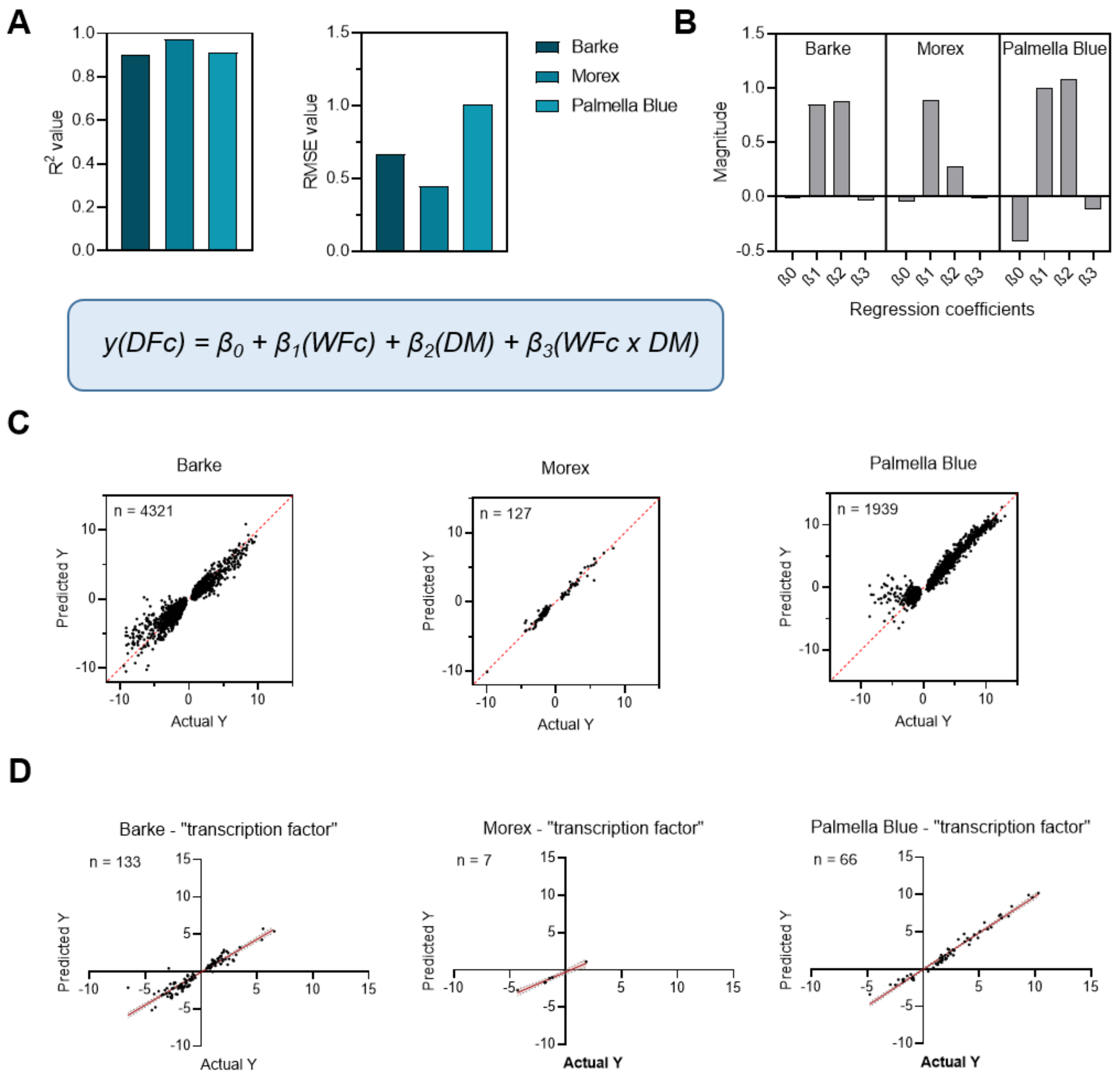


Figure S23: Multiple Linear Regression Analysis for prediction of gene expression under combination stress using all significantly regulated DEGs found in the transcriptomic study by Hoheneder et al. (2023). (A) displays the Goodness of fit ($R^2$) and respective Root Mean Squared Error obtained from the MLR model for significantly (FDR < 0.01) regulated DEGs under combination stress in each barley cultivar. The MLR model equation is given in the box. (B) shows the magnitude of regression coefficients for β0, β1 (main effect: WFc), β2 (main effect: DM) and β3 (interaction between WFc and DM) per cultivar and time point post infection. (C) represents the actual and predicted log2 fold changes for significantly regulated DEGs in DFc obtained from the MLR model for each barley cultivar and the two time points post-infection. (D) shows the actual and predicted log2 fold changes in combination stress for exemplary gene sets selected by keyword search in the obtained lists of differentially expressed genes for “transcription factor” in Barke, Morex and Palmella Blue.

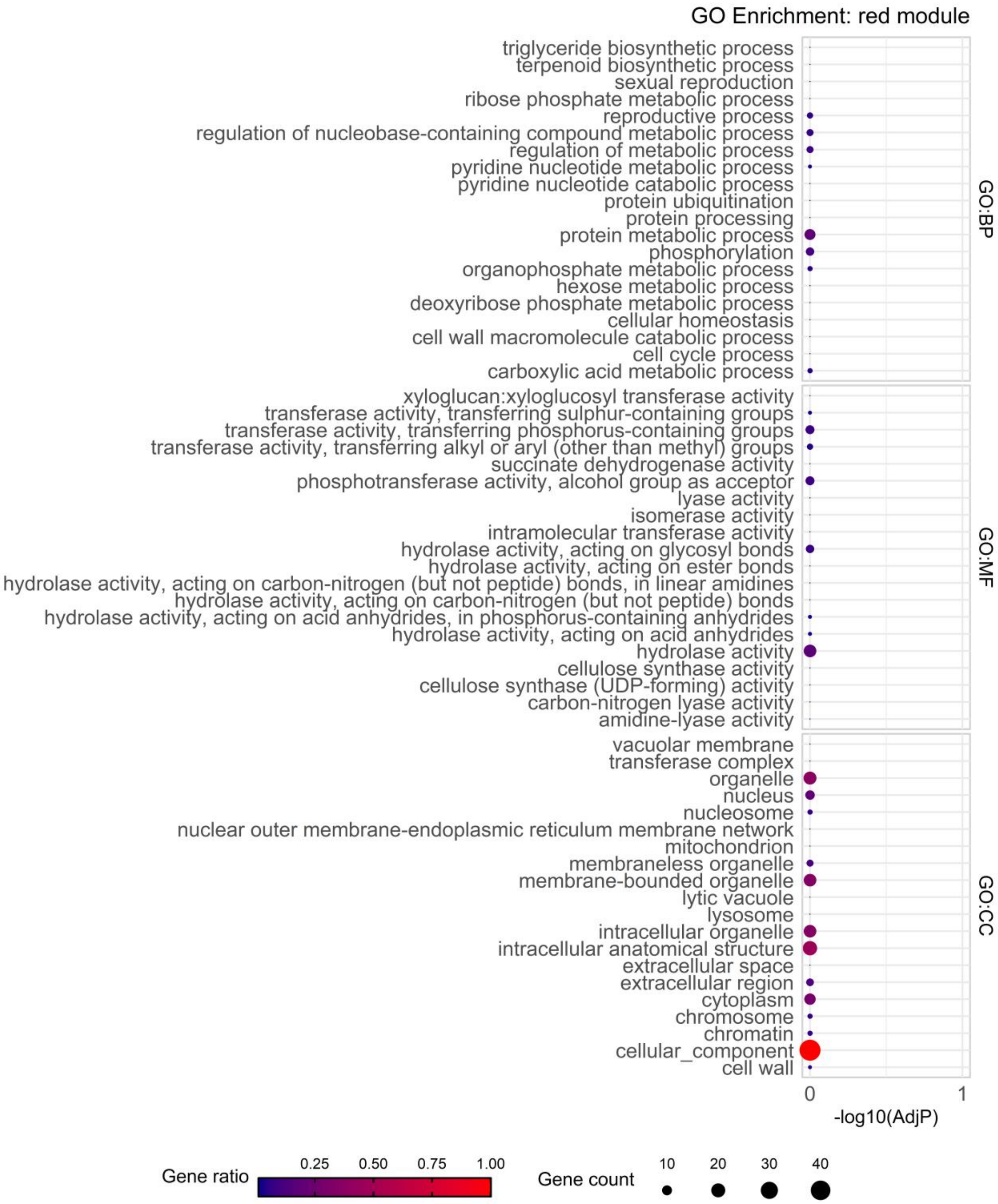


Figure S24: GO enrichment for DEGs found in WGCNA module red for biological process (BP), molecular function (MF) and cellular component (CC).

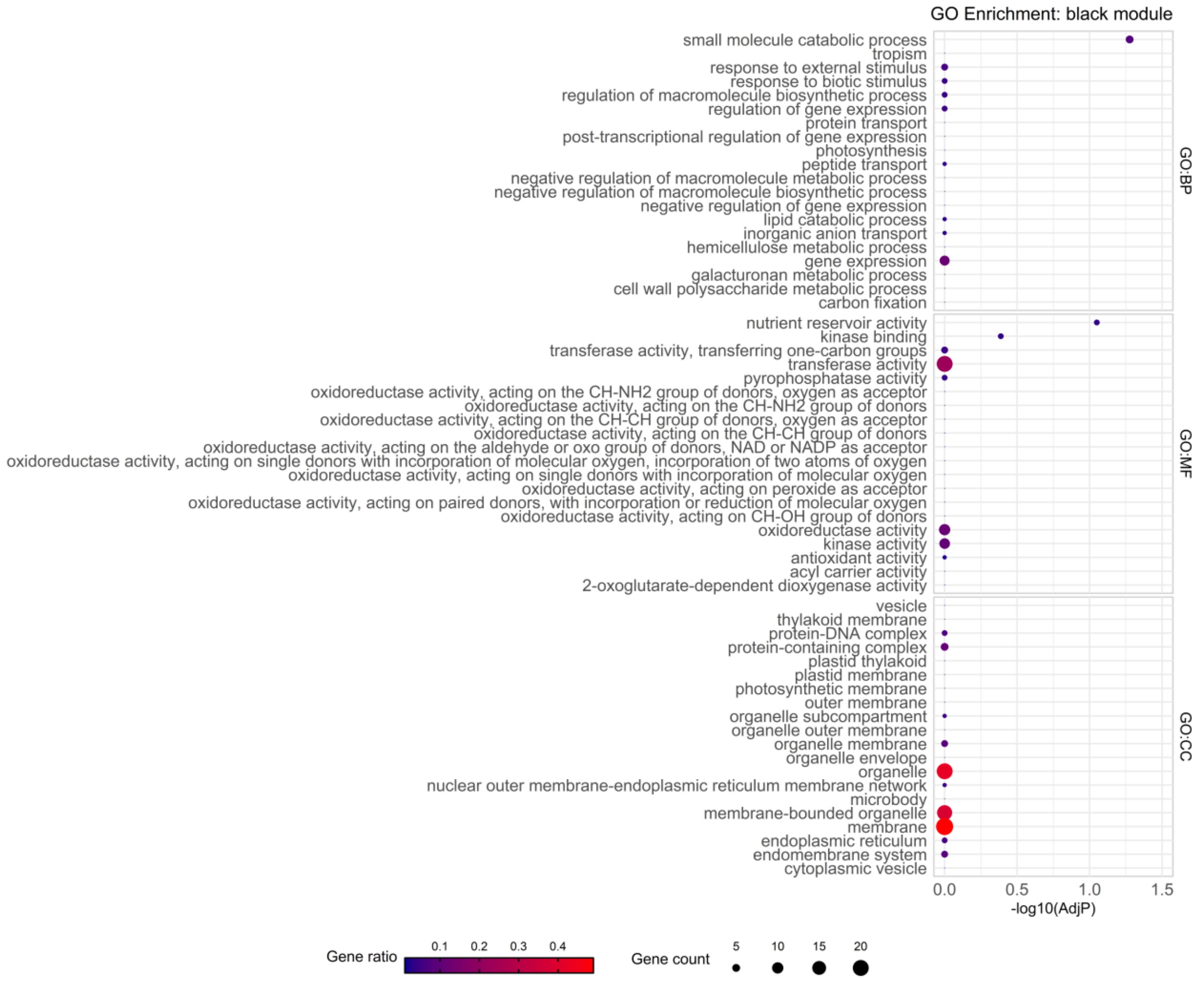


Figure S25: GO enrichment for DEGs found in WGCNA module black for biological process (BP), molecular function (MF) and cellular component (CC).

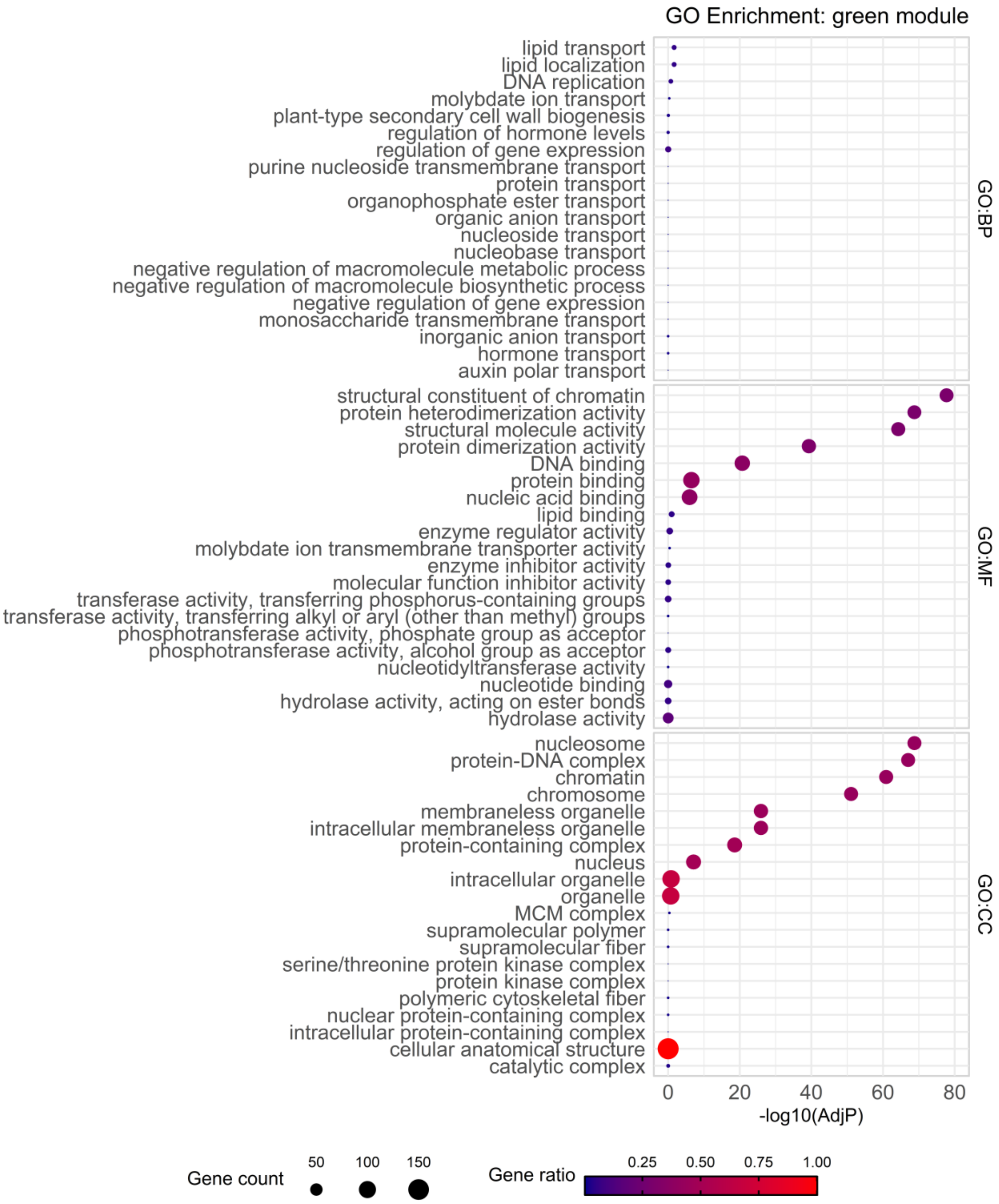


Figure S26: GO enrichment for DEGs found in WGCNA module green for biological process (BP), molecular function (MF) and cellular component (CC).

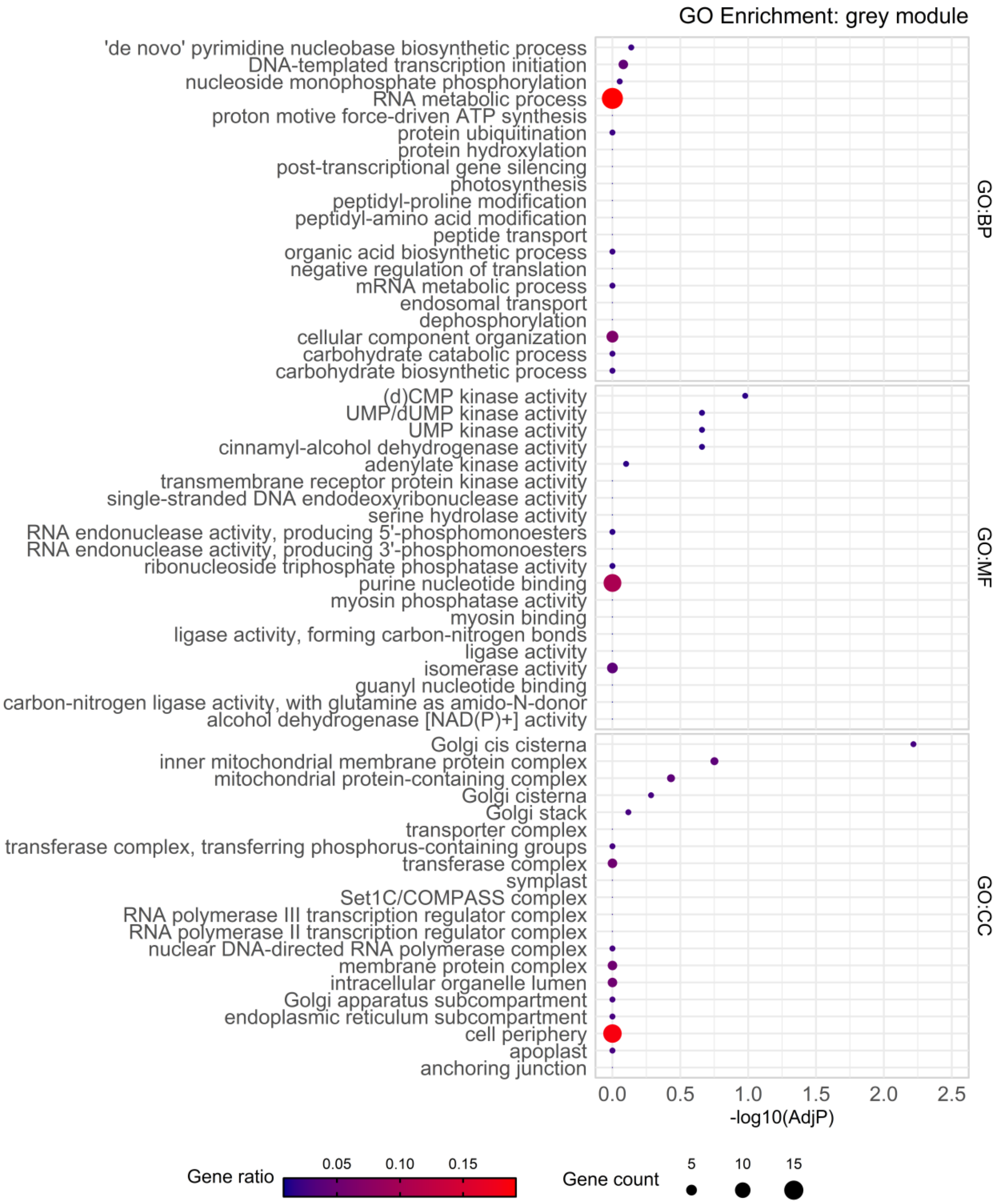


Figure S27: GO enrichment for DEGs found in cluster WGCNA module grey for biological process (BP), molecular function (MF) and cellular component (CC).

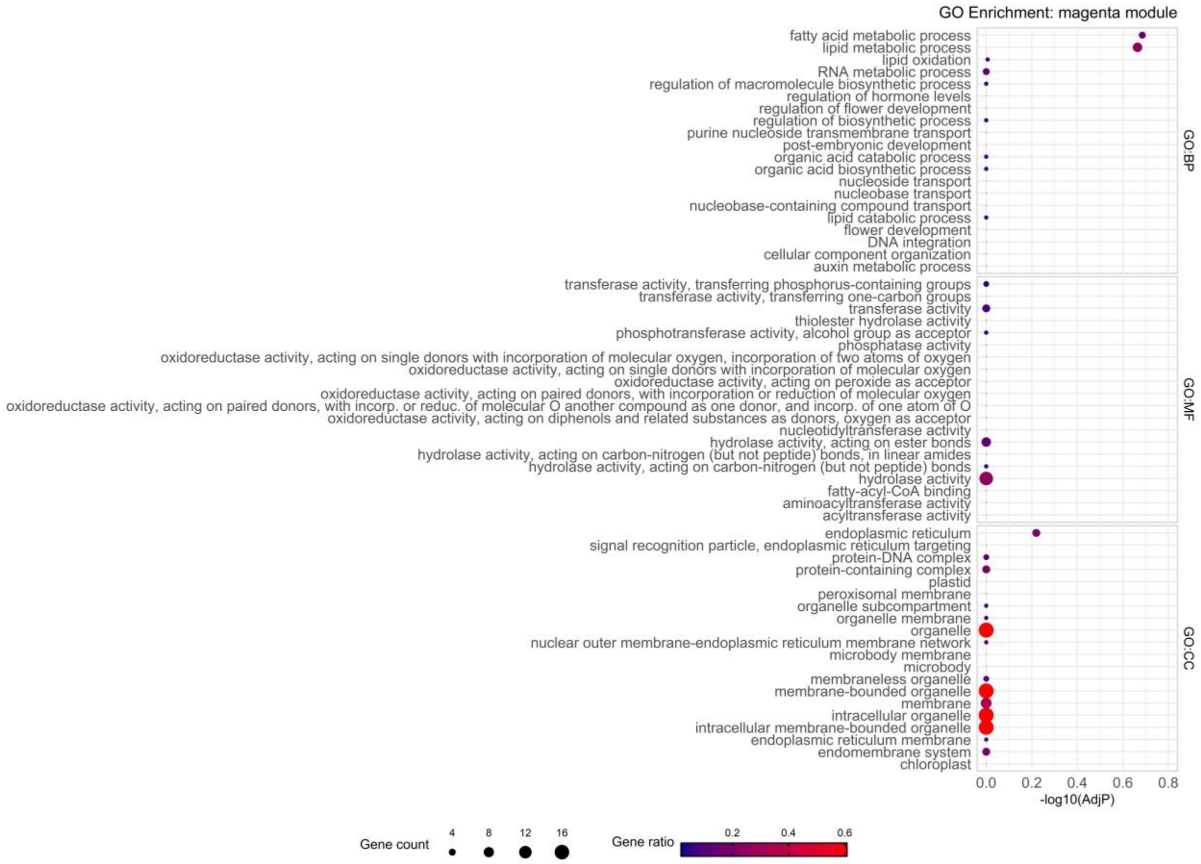


Figure S28: GO enrichment for DEGs found in WGCNA module magenta for biological process (BP), molecular function (MF) and cellular component (CC).

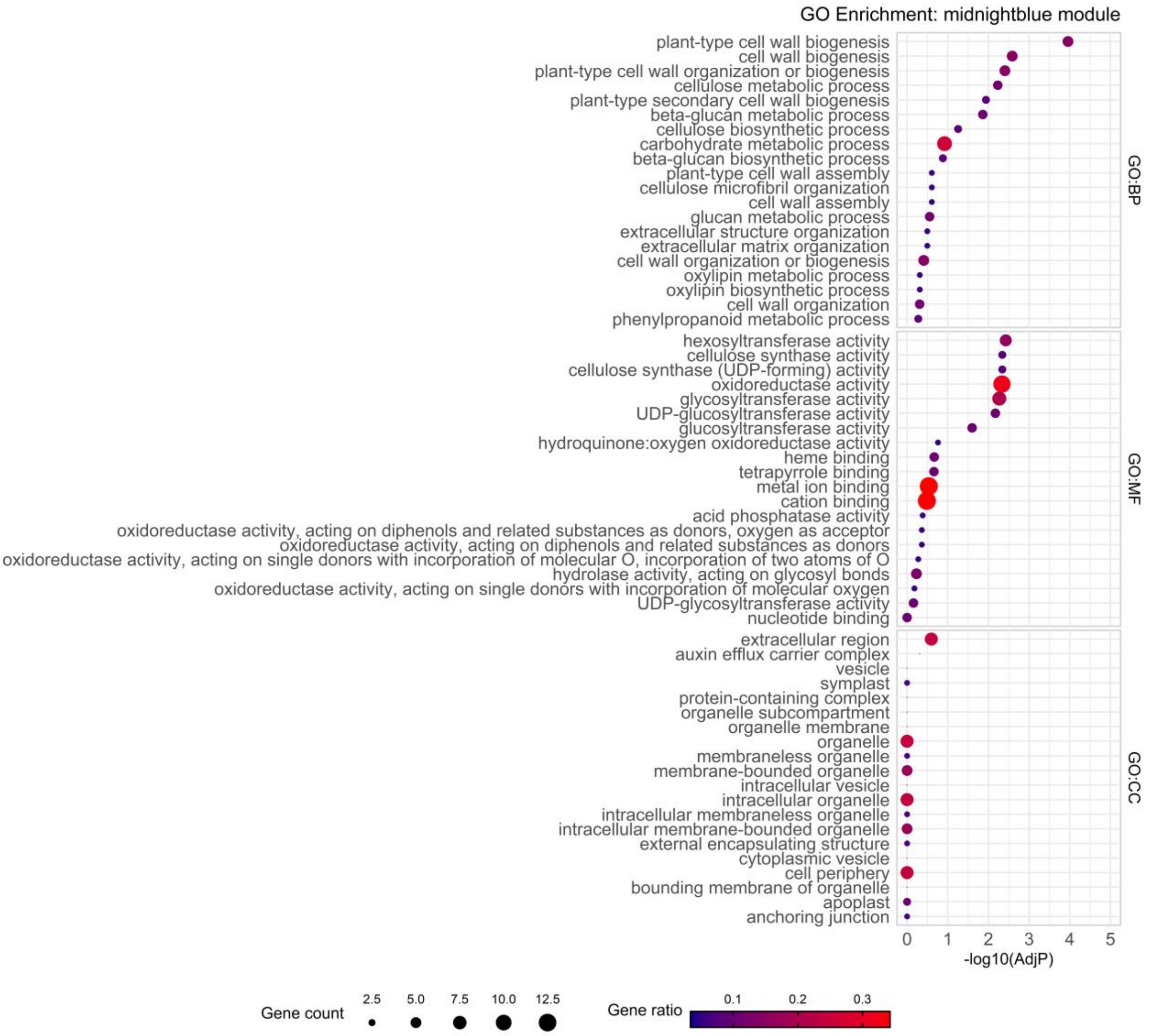


Figure S29: GO enrichment for DEGs found in WGCNA module midnightblue for biological process (BP), molecular function (MF) and cellular component (CC).

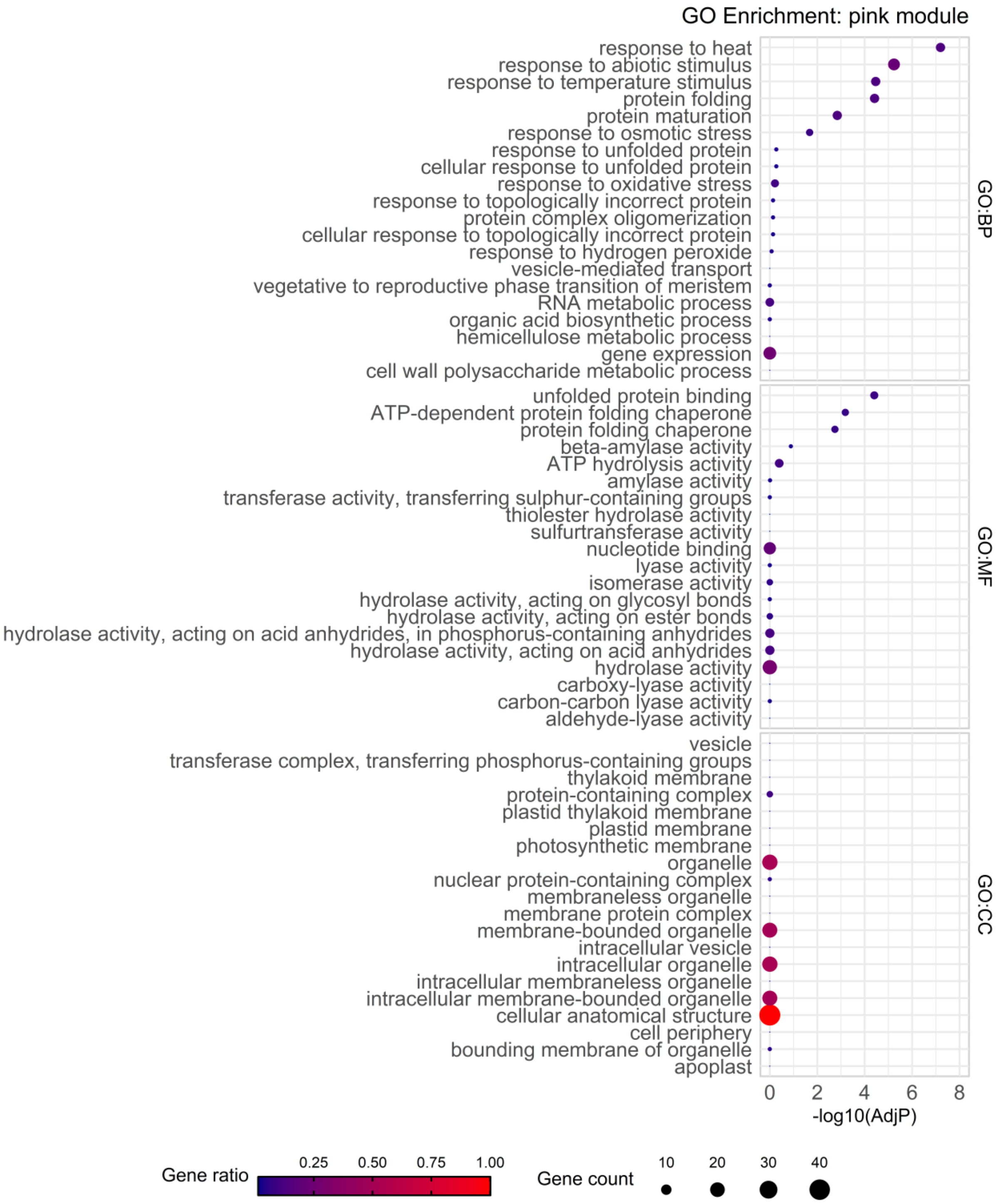


Figure S30: GO enrichment for DEGs found in WGCNA module pink for biological process (BP), molecular function (MF) and cellular component (CC).

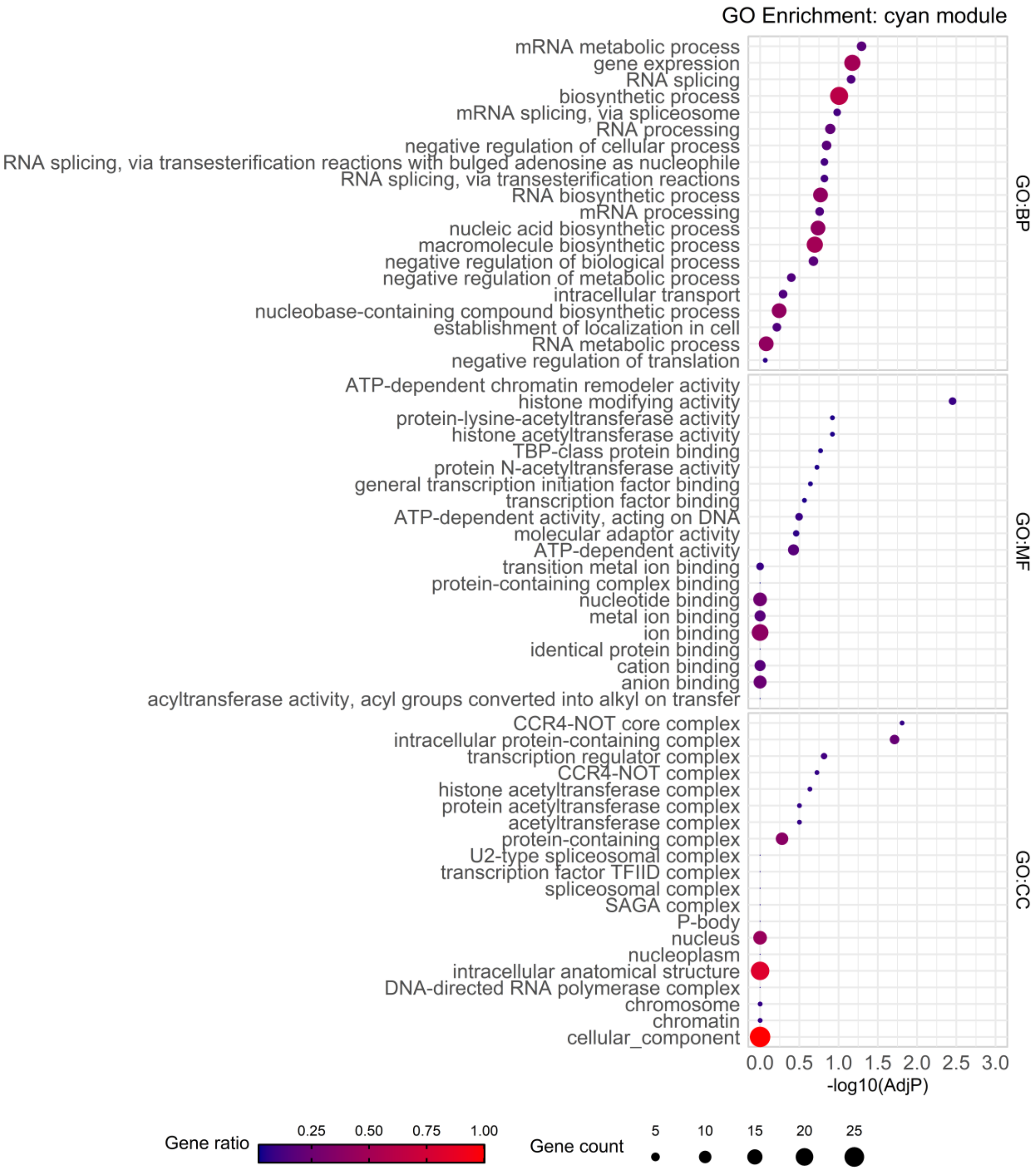


Figure S31: GO enrichment for DEGs found in WGCNA module cyan for biological process (BP), molecular function (MF) and cellular component (CC).

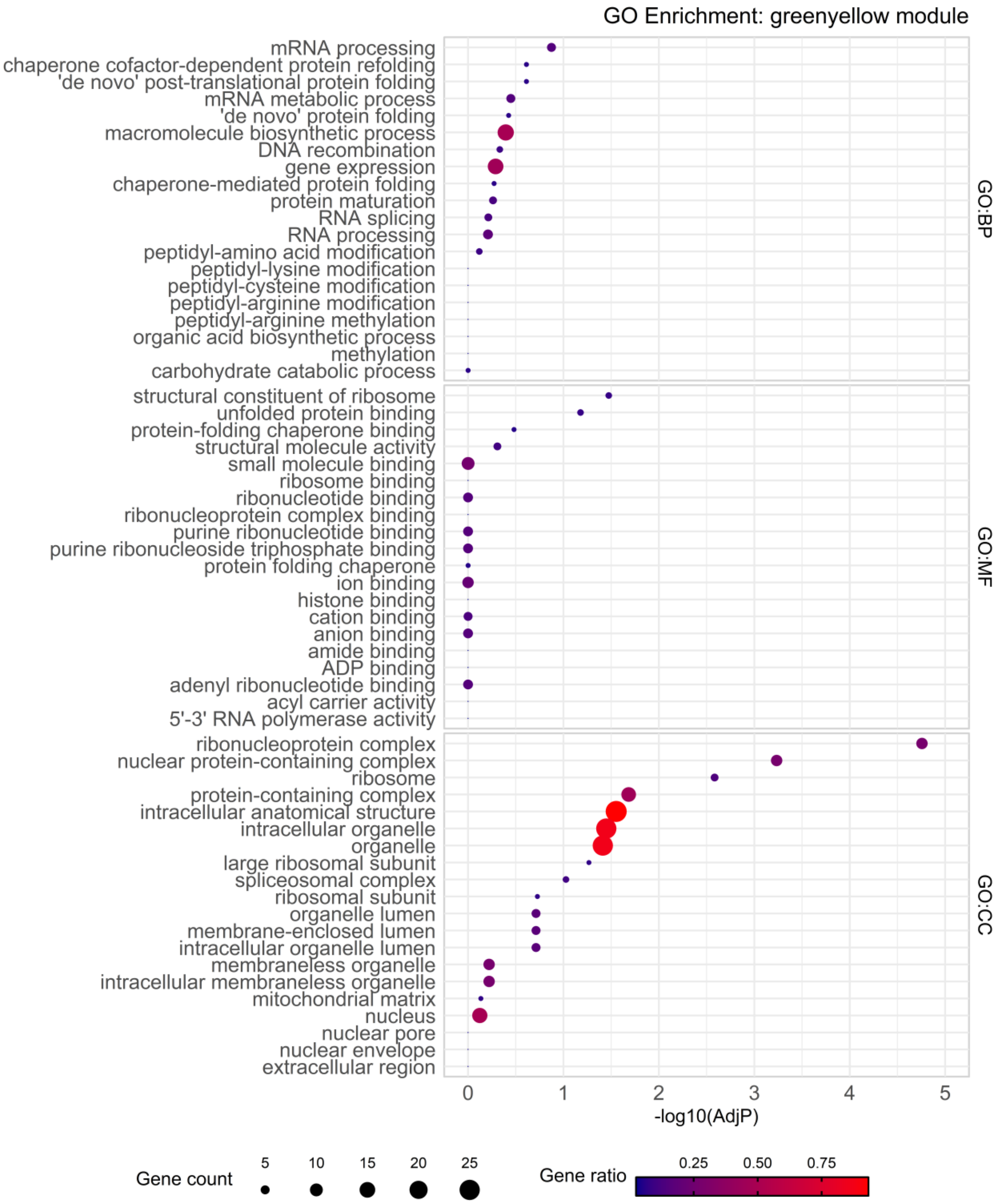


Figure S32: GO enrichment for DEGs found in WGCNA module greenyellow for biological process (BP), molecular function (MF) and cellular component (CC).

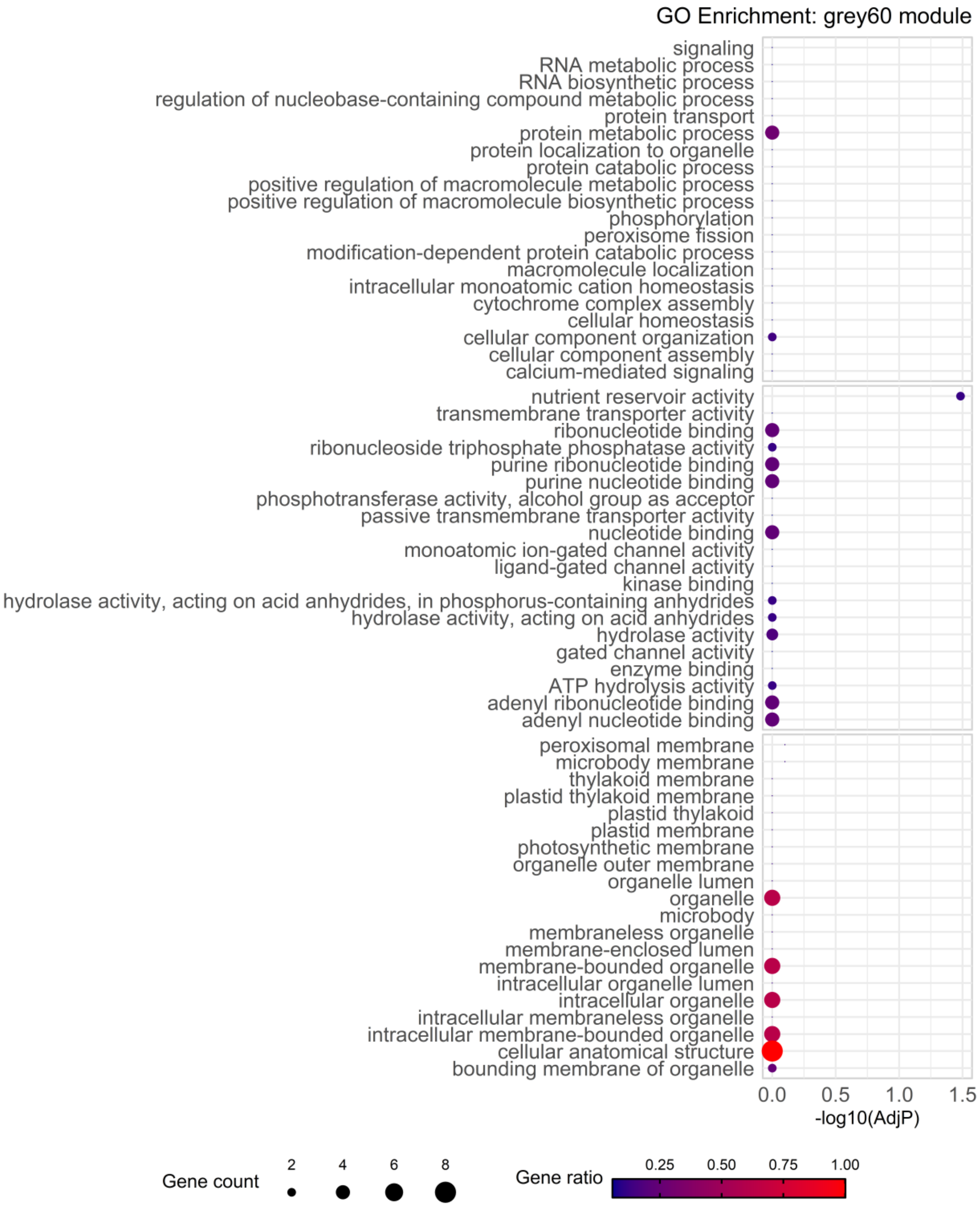


Figure S33: GO enrichment for DEGs found in WGCNA module grey60 for biological process (BP), molecular function (MF) and cellular component (CC).

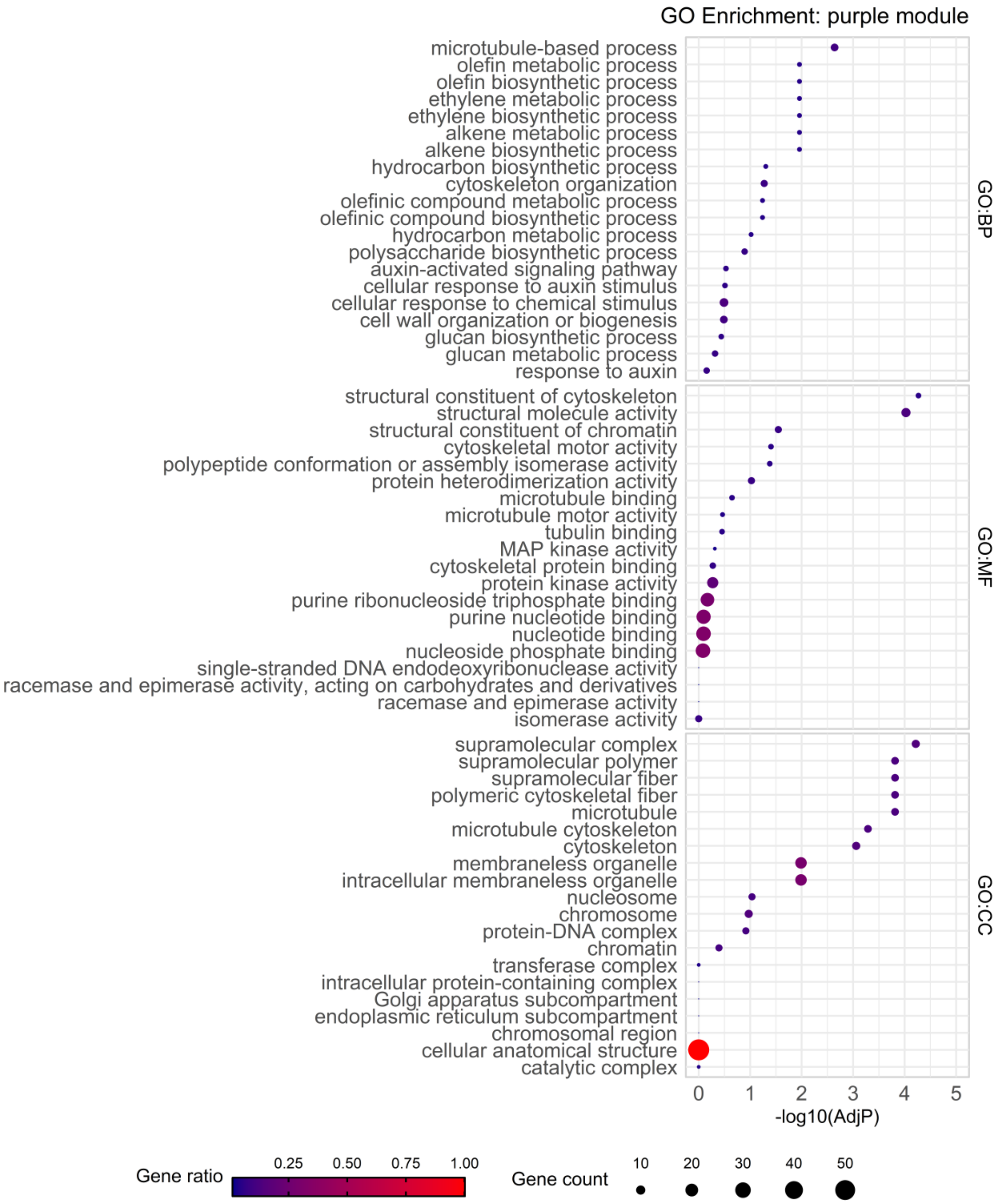


Figure S34: GO enrichment for DEGs found in WGCNA module purple for biological process (BP), molecular function (MF) and cellular component (CC).

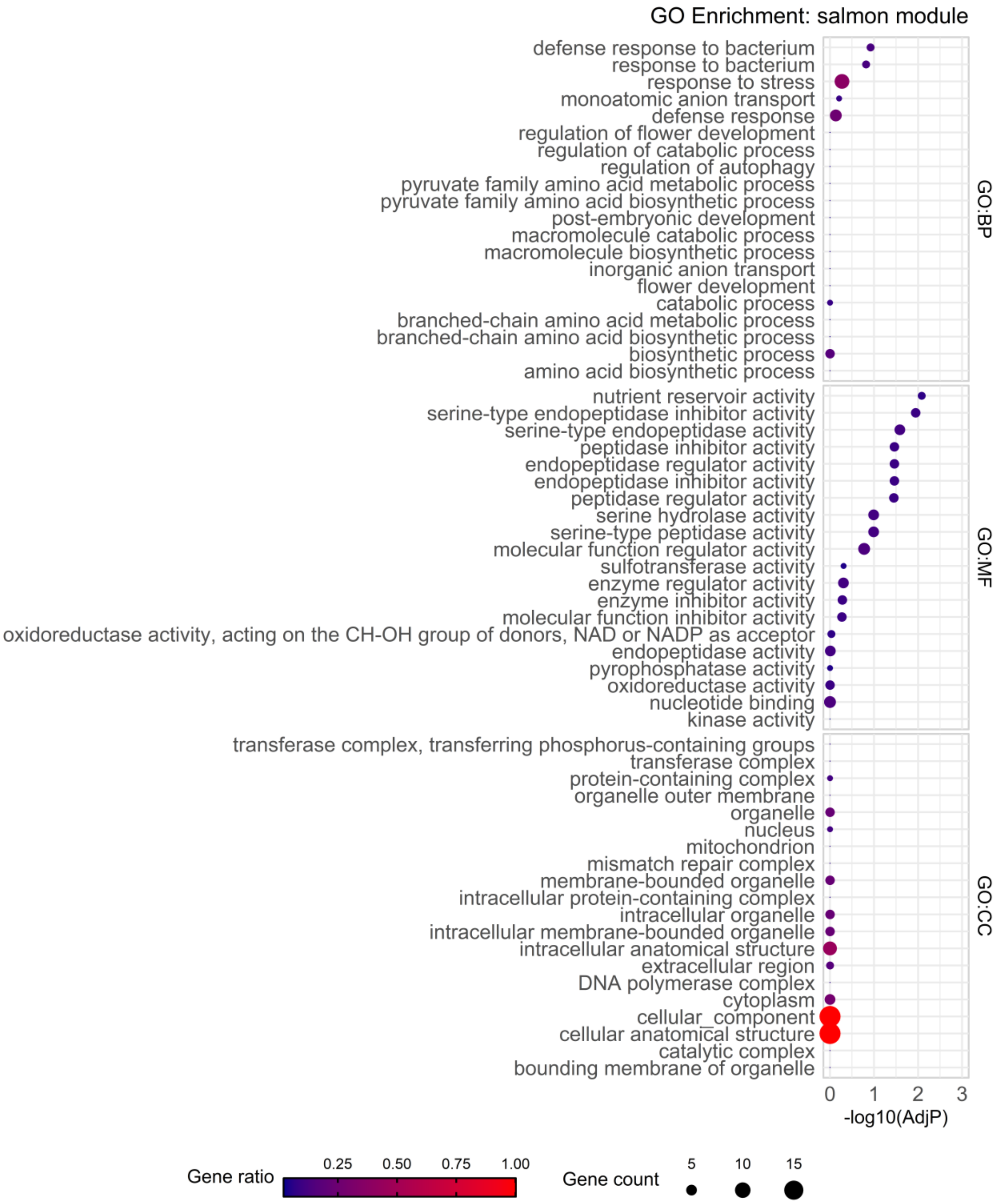


Figure S35: GO enrichment for DEGs found in WGCNA module salmon for biological process (BP), molecular function (MF) and cellular component (CC).

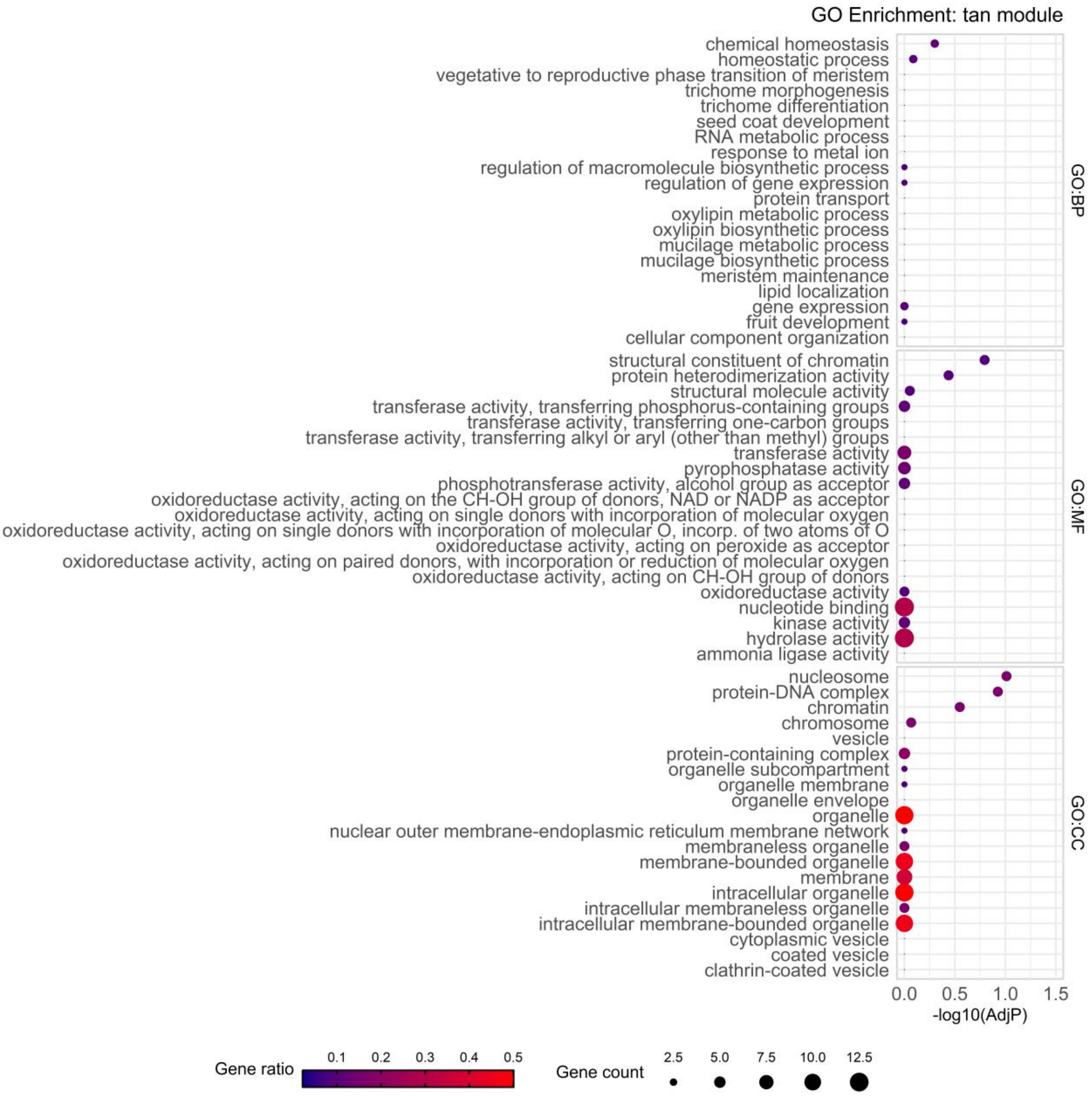


Figure S36: GO enrichment for DEGs found in WGCNA module tan for biological process (BP), molecular function (MF) and cellular component (CC).

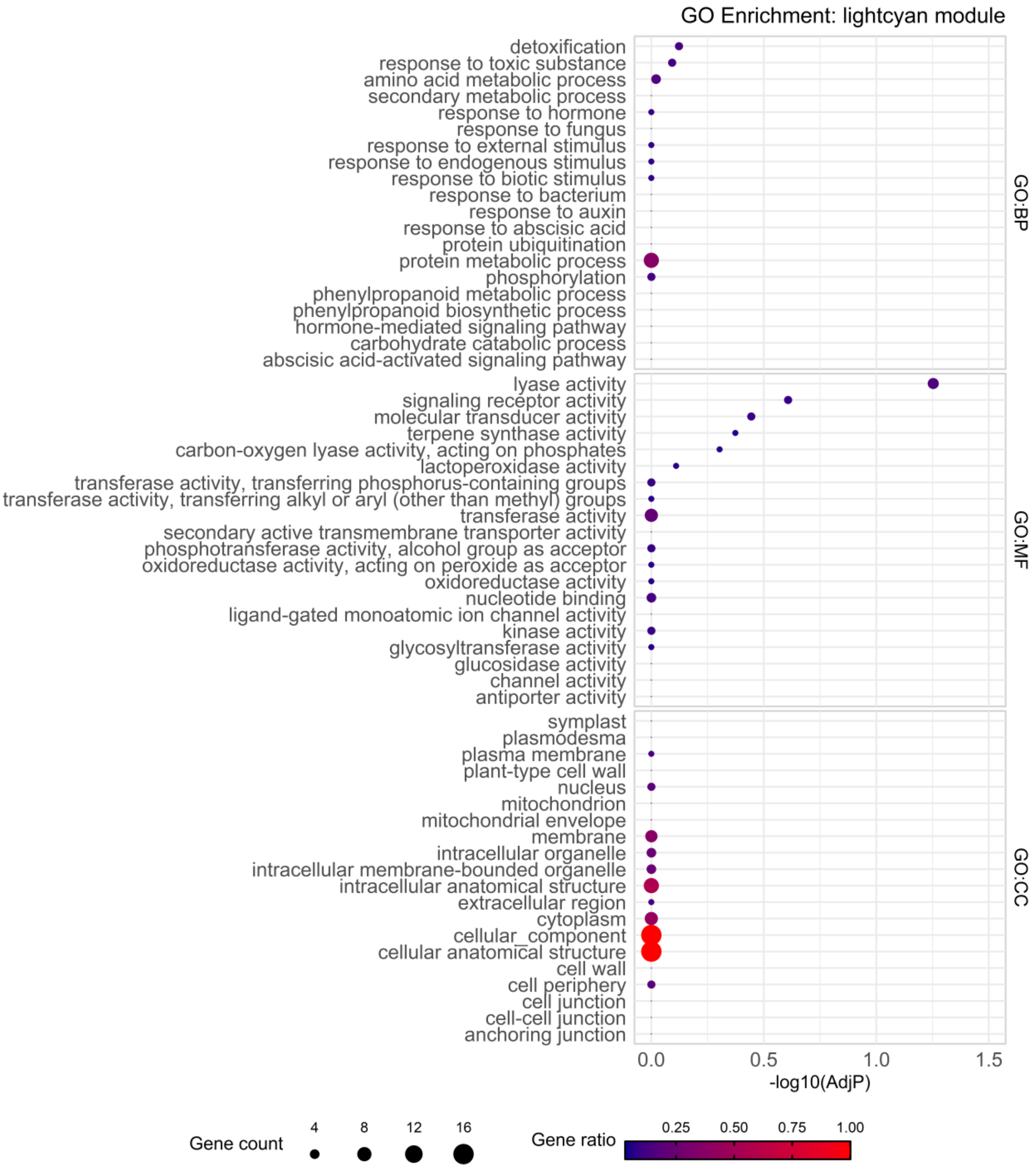


Figure S37: GO enrichment for DEGs found in WGCNA module lightcyan for biological process (BP), molecular function (MF) and cellular component (CC).
